# Identification of glucose-independent metabolic pathways associated with anti-proliferative effect of metformin, their coordinate derangement with cMyc downregulation and reversibility in liver cancer cells

**DOI:** 10.1101/2023.08.04.551931

**Authors:** Sk Ramiz Islam, Soumen Kanti Manna

## Abstract

Several studies indicated anti-cancer effects of metformin in liver cancer. This was attributed to the activation of LKB-AMPK axis, which is associated with anti-hyperglycaemic effect and cytotoxicity. However, despite lack of evidence on cytotoxic effect of physiological metformin concentrations and ability of cancer cells to survive under glucose-deprivation, no study has examined the glucose-independent effect of non-cytotoxic metformin or metabolic reprogramming associated with it. In addition, no study has ever been conducted on reversibility of anti-cancer effects of metformin. Here, the dose-dependent effects of metformin on HepG2 cells were examined in presence and absence of glucose. The longitudinal evolution of metabolome was analyzed along with gene and protein expression as well as their correlations with and reversibility of cellular phenotype and metabolic signatures. Metformin concentrations up to 2.5mM were found to be non-cytotoxic but anti-proliferative irrespective of presence of glucose. Apart from mitochondrial impairment, derangement of fatty acid desaturation, one-carbon, glutathione and polyamine metabolism were associated with non-cytotoxic metformin treatment irrespective of glucose supplementation. Depletion of pantothenic acid, downregulation of essential amino acid uptake, metabolism and purine salvage were identified as novel glucose-independent effects of metformin. These were significantly correlated with *cMyc* expression and reduction in proliferation. Rescue experiments established reversibility upon metformin withdrawal and tight association between proliferation, metabotype and *cMyc* expression. Taken together, derangement of novel glucose-independent metabolic pathways and concomitant cMyc downregulation co-ordinately contribute to anti-proliferative effect of metformin even at non-cytotoxic concentrations, which is reversible and may influence its therapeutic utility.

## Introduction

Liver cancer, which is the third leading cause of cancer-related deaths worldwide^1^, has poor prognosis and limited therapeutic options. Apart from viral hepatitis, non-alcoholic fatty liver disease (NAFLD) and type 2 diabetes, which are interlinked and increasing alarmingly, are major risk factors^2,3^. In epidemiological studies, metformin, an affordable and widely used anti-hyperglycaemic drug, has been found to decrease liver cancer incidence and increase overall survival^4,5^. While experimental studies showed its cytotoxic and anti-cancer effects, clinical trials largely remain non-conclusive on its utility in cancer^6^. *In vitro* studies that showed cytotoxic effects of metformin against variety of cancer cells (e.g., liver, breast and colon) ^7–11^ used metformin concentrations (up to 100mM) much higher than that found in plasma^12^. Most of these studies attributed the anti-cancer effect of metformin to the activation of LKB-AMPK axis which also contributes to its anti-hyperglycaemic effect^6,13^. But there is no direct evidence of cytotoxicity induced by metformin alone in tumors. Studies showed that metformin causes mitochondrial toxicity to compromise TCA cycle as well as entry of central carbons towards lipid biosynthesis^14–16^. Glucose and central carbon metabolism is the most important source of energy for cancer as well as other cells. Hence, inhibition of glucose metabolism by metformin can also affect activities of other cells including immune cells^17^ and may modulate efficacy of metformin in clinical trials. In fact, earlier studies showed that metformin repress mitochondrial metabolism and T-cell-mediated immune response^18,19^. However, cancer cells have been shown to reprogram several pathways other than glucose metabolism to harness energy as well as building blocks^20–23^. In fact, a subset of cancer cells can thrive even under glucose-deprived environment, which is found at the core of solid tumors^20–26^. These cells often exhibit stem-like properties and therapeutic resistance. Thus, understanding how metformin might affect pathways other than glucose metabolism is essential to rationalize its anti-cancer potential and may yield clues to develop therapeutic strategies that target non-central carbon pathways using metformin or its analogues. Till date, no comprehensive analysis of temporal evolution of metabolic landscape upon metformin treatment has been performed, which is essential to identify early and consistent metabolic signatures and pathways affected by metformin. It has also never been examined whether metformin-induced cellular phenotype and metabotype are reversible. This is essential to gauge the impact of stopping metformin use in patients at risk of liver cancer. Hence, this study examined the time-dependent effect of a non-cytotoxic metformin dose on cell fate, metabolic signature along with gene expression as well as the reversibility of the phenotype and molecular signatures upon metformin withdrawal in liver cancer cells.

## Experimental Procedures

### Chemical, solvents and reagents

All chemicals, MSTFA and GC-MS grade solvents were purchased from Sigma-Aldrich. LC-MS grade water and acetonitrile were purchased from Fischer Scientific. MOX reagent was purchased from Thermo Fischer. A detailed list of all authentic standards and their sources is provided in the supplementary table S1.

### Cell culture and experimental design

HepG2 cells were kind gifts from the laboratory of Prof. Debashis Mukhopadhyay (SINP, Kolkata). Cells were cultured with low glucose (5mM) DMEM media supplemented with 10% FBS, 100 I.U/ml penicillin and 50ug/ml streptomycin and maintained in a humidified 37°C incubator with 5% CO2. All experiments were performed within passage ten. Metformin (0.625 - 10mM) was added to DMEM with or without 5mM glucose and effects were analyzed at 3, 6-, 12-, 24- and 48-hour post-treatment. In the manuscript, treatment groups were designated as: **NG**- with 5mM glucose, **NT**- 2.5mM metformin with 5mM glucose, **WG**- no glucose supplementation, **WT**- 2.5mM metformin without glucose supplementation. For rescue experiment, cells were cultured under aforementioned conditions for 24 hours followed by media change and cultured for another 48hours as described below. The control (NG) group was continued with NG media (1). For NT group, either metformin was removed by replacing with NG media (2A) or exposed to only glucose deprivation using WG media (2B). For WG group, either glucose was replenished with NG media (3A) or exposed to metformin treatment in absence of glucose using WT media (3B). For WT group, either metformin was removed using WG media (4A) or glucose was replenished using NT media (4C) or both were done together using NG media (4B) or continued with WT media (4D) as control.

### CellTox green cytotoxicity assay

Cytotoxicity was measured using CellTox green cytotoxicity assay kit (Promega, Madison, USA) according to the manufacturer’s protocol. Fluorescence was measured using 96-well plates in BioTek Synergy HTX multimode reader (Agilent, Santa Clara, USA) with excitation and emission at 485nm and 520nm respectively.

### XTT assay

The cellular metabolic viability was measured by NADH-dependent mitochondrial dehydrogenase activity using Cell Proliferation Kit II (XTT) (Roche, Basel, Switzerland) according to the manufacturer’s protocol by measuring the absorbance at 492nm in BioTek Synergy HTX multimode reader. The XTT activities were presented as normalized with respect to the NG group.

### BrdU incorporation assay

Cellular proliferation was measured by Cell Proliferation ELISA, BrdU (colorimetric) kit (Roche, Basel, Switzerland) according to the manufacturer’s protocol with minor modification. Approximately 1000 cells/well were grown in a 96-well plate and incubated in specified conditions. 10uM of BrdU was added 3hr before the endpoint. After media removal, wells containing cells were dried in the air followed by fixed and incubated with the anti-BrdU-POD antibody for 90 min. After washing, the plate was incubated with substrate solution in the dark for colour development. The reaction was stopped by adding 1M sulphuric acid and absorbance was measured at 450nm. The BrdU incorporation was presented as normalized with respect to the NG group.

### Protein extraction and Western Blotting

Proteins from harvested cells were extracted with RIPA-lysis buffer and 15-30µg total protein was separated using 10% denaturing polyacrylamide gel followed by transfer methanol activated-PVDF membrane and was probed with anti-c-Myc antibody (ab32072, Abcam, Cambridge, UK). The blots were developed in X-ray film using an ECL kit (Invitrogen, Massachusetts, USA). Vinculin (#18799, Cell Signaling Technology, Massachusetts, USA) was used as a loading control. We used GelQuant.NET software provided by biochemlabsolutions.com for densiometric analysis.

### Total glutathione estimation

Total cellular glutathione was measured using Glutathione Assay Kit (Sigma-Aldrich, Missouri, USA) following the manufacturer’s protocol with minor modifications by colorimetric measurement of TNB dye at 412nm in BioTek Synergy HTX multimode reader.

### RNA isolation, RT-PCR

Total RNA from the cell was isolated using TRIzol™ Reagent (Invitrogen, Massachusetts, USA) following the given protocol cDNA was synthesized using High-Capacity cDNA Reverse Transcription Kit (Applied Biosystems™, Massachusetts, USA) according to the manufacturer’s protocol. In brief, 2µg of total RNA was reverse transcribed in a volume of 20ul.Relative expression was checked in the RT-PCR using PowerUp™ SYBR™ Green Master Mix (Applied Biosystems™, Massachusetts, USA) according to manufacturer’s protocol. The relative fold change of genes was calculated using the formula of 2^-(ΔΔCt). 18S rRNA was used as the reference gene. Sequence information of the primers are given in supplementary **table S2**.

### Network analysis

Network of genes were analyzed using STRING^27^ (https://string-db.org). Network edge thickness indicated the strength of data support. All active interaction sources were enabled. The minimum required interaction score was set to medium (0.400).

### Sample collection for metabolomics

Cells were grown in 6-well plates for up to 50-70% confluency before the treatment. After the indicated time points, cells were washed with ice-cold 150mM NaCl twice. 1 ml cold metabolite extraction solvent (water: acetonitrile: 2-propanol=2:3:3) containing 15uM homovalinic acid (internal standard) was added to each well. Cells were scraped and into 1.5ml polypropylene tubes (Eppendorf) and stored at -80°C freezer until further use.

### Sample preparation and derivatization

Samples were thawed in ice and vortexed vigorously, followed by three freeze-thaw cycle using liquid nitrogen. Supernatant containing metabolites were collected after centrifugation at 14000 x g RCF for 25min at 4°C. 100ul of supernatant was dried under vacuum followed by addition of 30ul MOX reagent and heated for 1 hour at 50°C. This was followed by addition of 50ul of MSTFA and heating at 65°C for 1hr. A pool of supernatants well as a blank sample were also prepared similarly for quality control.

### GC-MS Analysis

Derivatized samples were analyzed using Agilent 7890B gas chromatography (GC) system coupled with 5977B single-quadrupole mass spectrometer (MS). Samples were separated on a HP-5MS column (30m × 0.25mm × 0.25μm) (Agilent, Santa Clara, USA) using helium at a flow rate of 1ml/min. A retention time-locked data acquisition method (RT-lock) was developed against homovalinic acid for superior chromatographic alignment across the samples. The front inlet and MS transfer lines were set at 300°C. The oven temperature was set to 70°C, held for 2min followed by a ramp to 280°C at a rate of 5°C/min and held for 1min. The second ramp was set to 295°C at a rate of 10°C/min and held for 5min. A post-run equilibration was done at 300°C for 5min before coming down to the initial state. MS source and MS quad temperature were set to 230°C and 150°C respectively. Spectra were acquired in full scan mode in the m/z range of 50-600 with a focus m/z of 150 at a speed of 30Hz/sec. Before the individual sample injection, the pooled samples were injected five times for conditioning the column. Samples for each time point were prepared in a single batch and injected in GC-MS in randomized order with intermittent pooled and blank sample injections.

### Metabolomic data analysis

All chromatograms were manually checked for consistency. Feature extraction and integration was performed using MassHunter quantitative software (Agilent, Santa Clara, USA). Compound identity was based on NIST library score as well as matching with authentic standards wherever available (see supplementary information table S3). The integrated peak area table was used for univariate and multivariate data analysis. Features showing significant abundance in the blank were removed as well as those with CV > 30% for internal standard-normalized abundance in the pooled samples.

### Statistical analysis

Multivariate statistical analysis such as unsupervised principal components analysis (PCA), supervised volcano plot and heatmap analyses were performed using MetaboAnalyst5.0 (https://www.metaboanalyst.ca/MetaboAnalyst/home.xhtml)^28^ with sum-normalization, log transformation and Pareto scaling. A fold change cut-off > 1.5 and FDR correction was used to select metabolites of interest in volcano plot. MetaboAnalyst was also used for Pearson correlation analysis as well as for the identification of affected pathways with FDR correction. For individual metabolites, one-way ANOVA with Bonferroni correction was used to test statistical significance using GraphPad Prism 9 (https://www.graphpad.com). Changes between glucose concentration-matched pairs were considered to reflect the effect of metformin whereas those between metformin concentration-matched pairs were considered to reflect the effect of glucose. Throughout P < 0.05 was considered significant and denoted with asterisks as follows: <0.05 (*), <0.002(**), 0.001(***).

### Drawing tools

Microsoft PowerPoint 2022 (Washington, United States), Inkscape 1.2.2 (https://inkscape.org/), Adobe Illustrator 2022 (California, United States), and BioRender (https://app.biorender.com/) were been used for illustrations and composing figures.

## Results

### Metformin inhibits proliferation in time and dose-dependent manner irrespective of glucose supplementation

The XTT assay showed (Figure S1A) that 0.625-10mM metformin resulted in marginal decrease in viability in glucose-supplemented media with 12% decrease after 48hours with 10mM metformin. Glucose deprivation (WG) alone seemed to cause a significant decrease in viability at 12hours but recovered considerably by 24hours with marginal (12%) decrease at 48hours. The viability kept decreasing progressively upon metformin treatment (0.625-10mM) in glucose-deprived media after 24hours with 42% decrease at 48hours with 10mM metformin compared to WG. The BrdU incorporation (Figure S1B) was found to decrease after 24hours with ≥5mM metformin in glucose-supplemented media. At 48hours, it was found to decline at all metformin concentrations in dose-dependent manner. Glucose deprivation also decreased BrdU incorporation 24hour onwards. However, metformin caused dose-dependent decrease in BrdU incorporation after 12hours in glucose-deprived media compared to WG. These indicated that metformin caused dose-dependent progressive decrease in proliferation irrespective of glucose supplementation. However, it was found that there was significant cytotoxicity (Figure S1C) with metformin concentration ≥5mM. The cytotoxicity gradually increased 24hours onward, particularly with metformin concentrations ≥5mM in glucose-deprived media. There was no significant cytotoxicity irrespective of glucose supplementation with metformin ≤2.5mM. Taken together, these indicated that while metformin concentration ≤2.5mM did not cause any significant cytotoxicity, it affected viability and attenuated proliferative capacity irrespective of glucose-supplementation.

As shown in Figure 1A, in glucose-supplemented media while there was a 16% decrease in viability in presence of non-cytotoxic 2.5mM metformin (NT) at 48hours, cell proliferation decreased by 83% compared to NG (Figure 1B). Glucose deprivation (WG) alone resulted led to decrease in proliferation 24hour onwards with marginal change in viability (Figure 1A-B). 2.5mM metformin in glucose-deprived media (WT) not only caused 28% loss of viability compared to WG at 48hr but also decreased cell proliferation 12hour onwards without any cytotoxicity. At 12, 24 and 48hours, cell proliferation decreased by 45%, 79%, and 76%, respectively, in WT compared to WG. These indicated that metformin inhibited cell proliferation even without cytotoxicity irrespective of glucose supplementation. All subsequent analyses including metabolomics (Figure 1C) were carried out using 2.5mM metformin to identify glucose-independent pathways.

**Figure 1.**
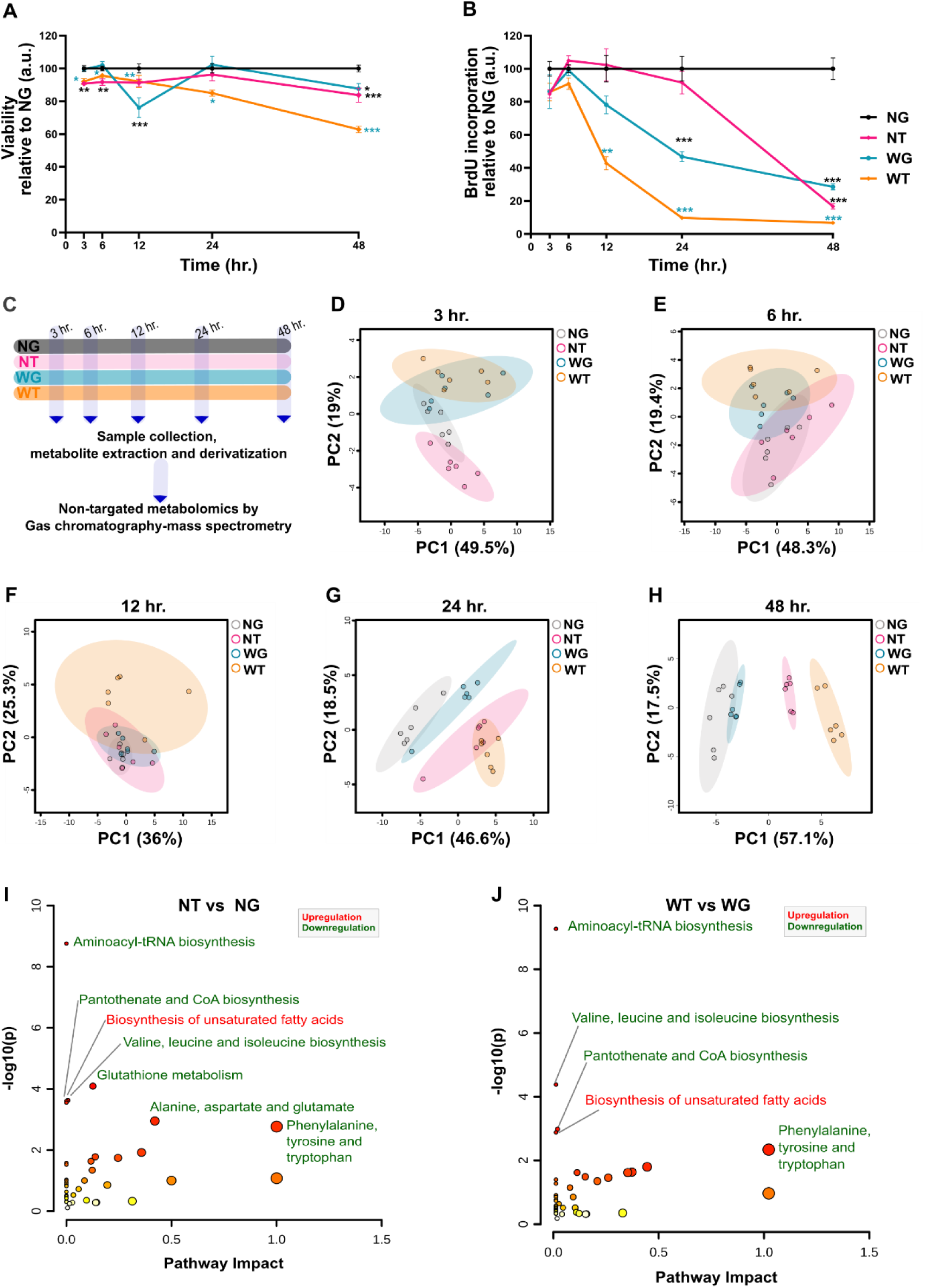
*Effects of metformin on cell viability, proliferation and global metabotype in with and without glucose supplementation. **A.** Effect on cell viability from 3 hour to 48 hour after treatment assessed by XTT assay. NG - with 5mM glucose, NT - 2.5mM metformin with 5mM glucose, WG - no glucose supplementation, WT - 2.5mM metformin without glucose supplementation. The absorbance at each time point was normalized with respect to mean absorbance in the NG group at that time point (data presented as mean ± SEM, n = 6) **B.** Effect on proliferation assessed by BrdU incorporation assay from 3 hour to 48 hour after treatment. The absorbance at each time point was normalized with respect to mean absorbance in the NG group at that time point (data presented as mean ± SEM, n = 6). One-way ANOVA with Bonferroni post-hoc test for multiple comparison was used to test statistical significance. P values <0.05(*), <0.002(**), <0.001(***) are indicated and color-coded for pair-wise comparisons (black: comparison with NG, cyan: comparison with WG and pink: comparison with NT). **C.** Experimental design for global metabolic profiling. **D-H.** 2D scores scatter plots for principal components analysis (PCA) showing temporal evolution of global metabotype with time points indicated for respective plots. **I, J.** Results of pathway analysis showing effect of metformin on metabolic pathways. Upregulated (red) and downregulated (green) pathways due to metformin treatment in presence of glucose (NT) and in absence of glucose (WT) compared to NG and WG, respectively, are highlighted. Multivariate analysis of the sum-normalized, log-transformed and Pareto-scaled metabolomic data was performed using MetaboAnalyst 5.0. Volcano plot analysis (with fold change and FDR-corrected P value cut off 1.5 and 005, respectively) was used identify differential metabolic signatures which were further used for pathway analysis using MetaboAnalyst.*

### Temporal evolution of metabolic signatures parallels cell proliferation

The PCA scores plots showed that there was no clear segregation of the overall metabotype upon glucose deprivation and/or metformin treatment at 3 or 6hours post-treatment along PC1 (Figure 1D, E). The WT showed a tendency of segregation from the rest after 12hour (Figure 1F), which coincided with decrease in proliferation. At 24hour, the metabotype of WG, which showed a decrease in proliferation, was slightly adrift from NG while NT and WT clustered away from them along PC1(Figure 1G). At 48hours, while NG and WG clustered closely, WT drifted furthest from them while NT was found in between along PC1 (Figure 1H), trends similar to that observed for proliferation. This indicated that the metformin treatment caused gradual and progressive derangement of metabolic signature 12hours onwards leading to distinct metabotypes. The separation of metabotypes at 48hours paralleled trends for proliferation showing dominant effect of metformin irrespective of glucose supplementation. Pathway analysis indicated that while aminoacyl-tRNA biosynthesis, glutathione metabolism, pantothenate and CoA biosynthesis, valine, leucine and isoleucine biosynthesis, alanine, aspartate and glutamate metabolism, phenylalanine, tyrosine and tryptophan biosynthesis were downregulated, biosynthesis of unsaturated fatty acids was upregulated upon metformin treatment in presence of glucose (Figure 1I, Table S4). In absence of glucose, amino-acyl tRNA biosynthesis, valine, leucine, isoleucine, phenylalanine, tyrosine, tryptophan biosynthesis as well as pantothenate and CoA biosynthesis was downregulated, whereas unsaturated fatty acid biosynthesis was upregulated (Figure1J, Table S5).

A comparison of metabolic signatures at 48hour (Figure S2A, Table S6) revealed decrease in valine, isoleucine, phenylalanine, tyrosine, cysteine, taurine, glycine, threonine, aspartic acid, tryptophan, pantothenic acid, adenine and increase in lactic acid, myristic acid, palmitic acid, oleic acid, linoleic acid, arachidonic acid, mono-palmitin, and hypoxanthine to be common in both NT and WT but absent in WG indicating them to be glucose-independent effects of metformin. These common signatures, indeed, represented aminoacyl-tRNA biosynthesis, pantothenate and CoA biosynthesis, valine, leucine and isoleucine biosynthesis, phenylalanine, tyrosine and tryptophan biosynthesis and biosynthesis of unsaturated fatty acids pathways (Table S7).

### Effect of metformin on central carbon pathway and fatty acid metabolism

Analysis of metabolic signature revealed significant impact on central carbon metabolism, which is connected to fatty acid and amino acid metabolism as shown in Figure 2A. Citric and malic acids remained low throughout in NT and WT compared to respective controls (Figure 2B-C). These metabolites were also lower in WG compared to NG. The expression of ACC1 and ACC2 were found to be unaltered in WG and depleted similarly in NT and WT compared to respective controls (Figure 2D-E). Palmitic, stearic, myristic acids were elevated similarly upon metformin treatment 24hour onwards (Figure 2F-H). Alongside, unsaturated oleic, linoleic and arachidonic acids were elevated upon metformin treatment (Figure 2I-K). In addition, monopalmitin also showed a trend of elevation 24hour onwards in WT and was elevated in both NT and WT 48hours (Figure 2L). Interestingly, pantothenic acid (Figure 2M) was found to be significantly depleted in NT 24hour onwards whereas it was depleted 6hour onwards in WT compared to respective controls, preceding decrease in proliferation. On the other hand, lactic acid (Figure 2N) was elevated in NT compared to NG throughout. Even in WT, lactic acid was found to be elevated 24hour onwards.

**Figure 2.**
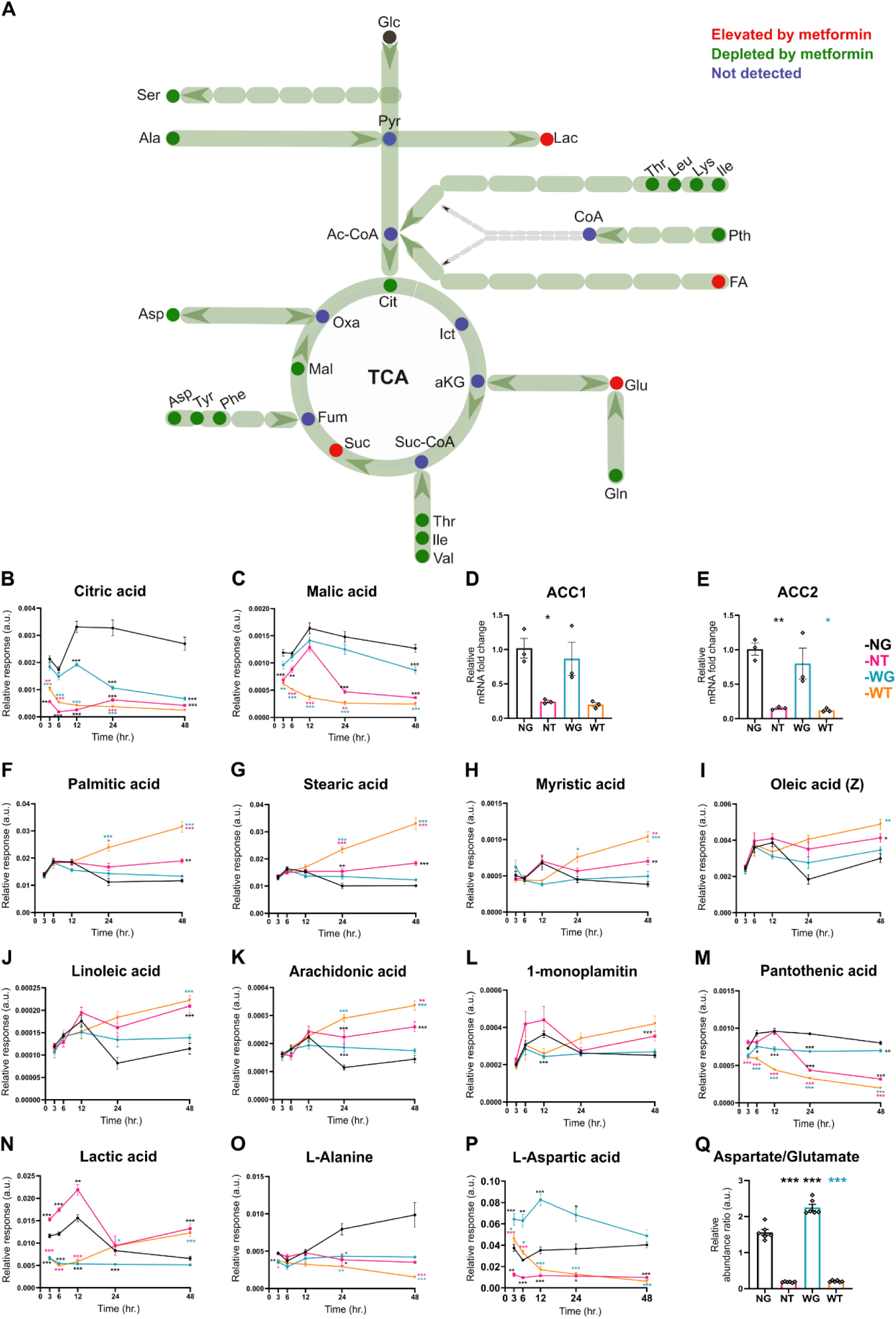
*Effects of metformin on central carbon pathway and fatty acid metabolism. A. Schema showing the effect of metformin on central carbon metabolism including connection to amino acid and fatty acid metabolism (abbreviations: Glc-Glucose, Pyr-Pyruvate, Ser-Serine, Ala-Alanine, Lac-Lactic acid, Thr-Threonine, Leu-Leucine, Lys-Lysine, Ile-Isoleucine, Pth-Pantothenic acid, CoA-Coenzyme A, Ac-CoA-Acetyl coenzyme A, FA-Fatty acids, Cit-Citric acid, Ict-Isocitric acid, aKG-alpha-ketoglutaric acid, Suc-CoA-Succinyl coenzyme A, Suc-Succinic acid, Fum-Fumaric acid, Mal-Malic acid, Oxa-Oxaloacetic acid, Glu-Glutamic acid, Gln-Glutamine, Val-Valine, Asp-Aspartic acid, Tyr-Tyrosine, Phe-Phenylalanine). The trend plots for longitudinal changes in relative abundances of citric acid (**B**), malic acid (**C**), palmitic acid (**F**), stearic acid (**G**), myristic acid (**H**), oleic acid (**I**), linoleic acid (**J**), arachidonic acid (**K**), 1-monopalmitin (**L**), pantothenic acid (**M**), lactic acid (**N**) L-alanine (**O**), and L-aspartic acid (**P**) are shown along with aspartate/glutamate ratio (Q) and expression of acetyl-CoA carboxylase alpha (ACC1, **D**) and acetyl-CoA carboxylase beta (ACC2, **E**) genes at 48 hours. All data presented as mean ± SEM with n = 6 for metabolites and 3 for ACC1 and ACC2. One-way ANOVA with Bonferroni post-hoc test for multiple comparisons was used to test statistical significance. P values <0.05(*), <0.002(**), <0.001(***) are indicated and color-coded for pair-wise comparisons (black: comparison with NG, cyan: comparison with WG and pink: comparison with NT).*

These indicated a general depression in TCA cycle along with impairment of fatty acid oxidation and increase in glycolytic flux upon metformin treatment. Alanine (Figure 2O), which can supply pyruvate for glycolysis with concomitant glutamate production, was found to decrease significantly in both NT and WT 24hour onwards. Alanine/glutamate ratio (Figure S2B) was found to be reduced to similar levels in both NT and WT at 48hours. Aspartic acid (Figure 2P), which may be produced from TCA intermediate oxaloacetate with concomitant consumption of glutamate, was found to be depleted throughout upon metformin treatment irrespective of glucose supplementation. Aspartate/glutamate ratio also reduced upon metformin treatment and was found at similar levels at 48hours in NT and WT (Figure 2Q).

### Effects of metformin on essential and non-essential amino acid uptake and metabolism

The abundance of several essential (EAA) as well as non-essential (NEAA) amino acids were found to be significantly affected upon metformin treatment. While they tended to increase in WG, the abundances of branched-chain amino acid (BCAA; isoleucine, leucine, and valine) tended to decrease upon metformin treatment and was significantly lower in both NT and WT at 48 hours (Figure 3A-C).

**Figure 3.**
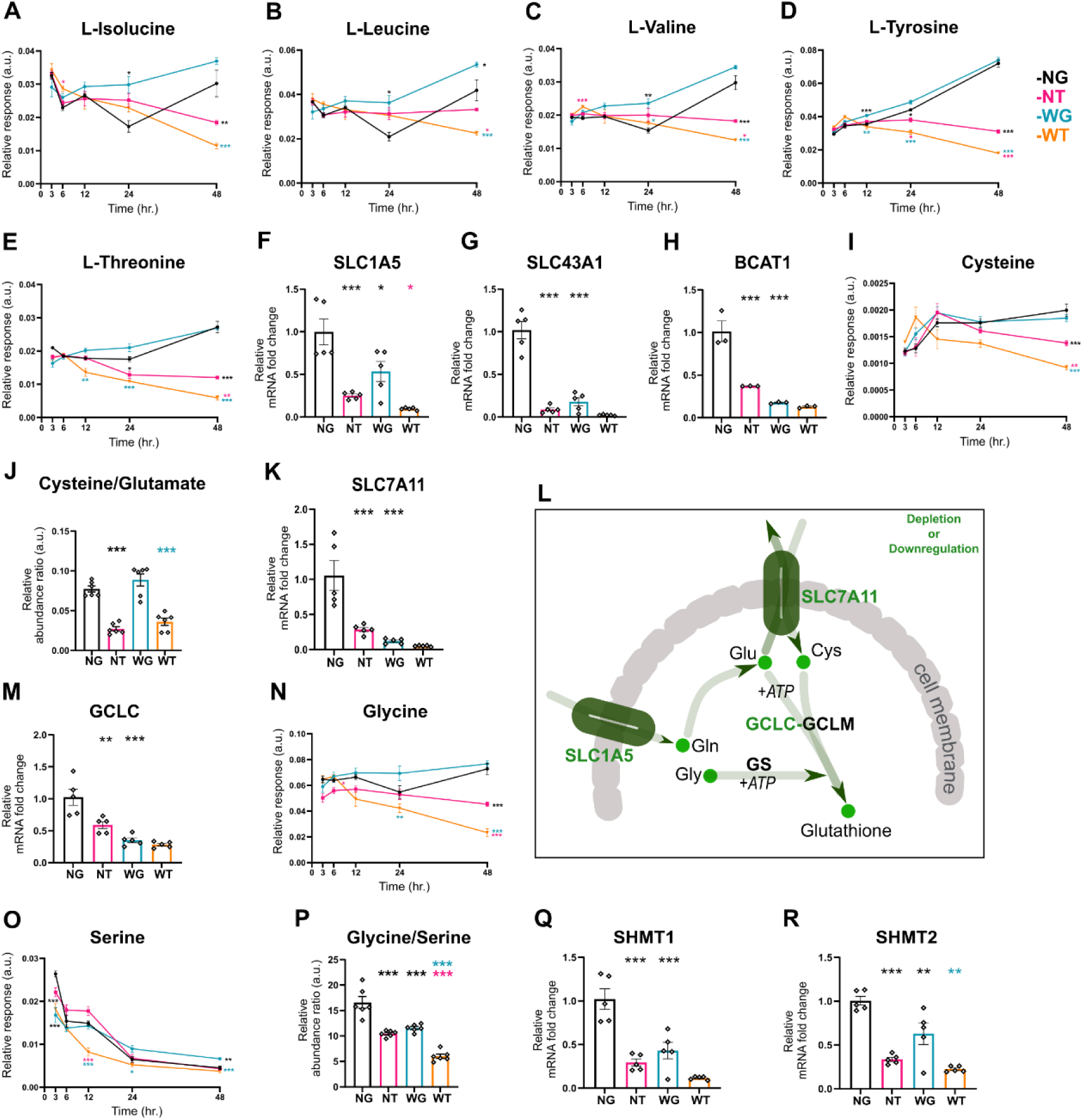
*Effects of metformin on amino acid uptake, metabolism and glutathione biosynthesis. The trend plots for longitudinal changes in relative abundances of isoleucine (**A**), leucine (**B**), valine (**C**), tyrosine (**D**), threonine (**E**), cysteine (**I**), glycine (**K**), and serine (**N**) are presented along with cysteine/glutamate (**J**) and serine/glycine (**P**) ratios at 48 hours. Expression of solute carrier family 1 member 5 (SLC1A5, **F**), solute carrier family 43 member 1 (SLC43A1, **G**), branched-chain amino acid transaminase 1 (BCAT1, **H**), solute carrier family 7 member 11 (SLC7A11, **K**), glutamate-cysteine ligase catalytic subunit (GCLC, **M**), serine hydroxymethyltransferase 1 (SHMT1, **Q**), and serine hydroxymethyltransferase 2 (SHMT2, **R**) genes at 48 hours are also presented. All data presented as mean ± SEM with n = 6 for metabolites and 5 for genes. One-way ANOVA with Bonferroni post-hoc test for multiple comparisons was used to test statistical significance. P values <0.05(*), <0.002(**), <0.001(***) are indicated and color-coded for pair-wise comparisons (black: comparison with NG, cyan: comparison with WG and pink: comparison with NT).*

A progressive decline, was also found for tyrosine and threonine (Figure 3D-E) 12 and 24hour onwards, respectively, in WT and NT, which either coincided or preceded decrease in proliferation. At 48hours (Figure S3A-G), abundance of BCAAs as well as other EAAs like threonine, phenylalanine, tyrosine and tryptophan were found depleted in both NT and WT whereas they either remained same or increased slightly (for leucine, phenylalanine and tryptophan) in WG. Expression of transporters of these amino acids (SLC1A5, SLC43A1, SLC3A2, LAT1) as well as BCAT1, which is involved in BCAA catabolism, were downregulated upon metformin treatment (Figure 3F-H and S3H, I). Among NEAAs, cysteine level was found to show a declining trend in WT at 24 hours and depleted in both NT and WT at 48hours (Figure3I). This was associated with slightly elevated level of glutamic acid in these conditions (Figure S3J). At 48hour the cysteine/glutamate ratio declined equally upon metformin treatment irrespective of presence of glucose and it did not change in WG (Figure 3J). The expression of the antiporter SLC7A11 (Figure 3K), which imports cystine in lieu of glutamate, was found to be depleted by10-fold in WG and 3.5, and 2-folds, respectively, in NT and WT compared to respective controls. The glutamine level tended to decrease in WT 12hour onwards as well as in NT and WG at 24hours (Figure S3K). At 48hours, it was found to be depleted similarly in WG and WT (Figure S3L). Cysteine metabolite taurine was also found to decrease significantly in NT and WT compared to respective controls at 48hours (Figure S3M). Expression of other amino acid transporters such as SLC16A10 (aromatic amino acids) and SLC7A1 (cationic amino acids) were also downregulated upon metformin treatment in presence of glucose as well as upon glucose deprivation alone (FigureS3N-O).

### Effects of metformin on one-carbon and glutathione metabolism

The glutamate and cysteine combine with glycine to produce glutathione as shown in Figure 3L. It was found that while there was no change in total glutathione in NT, it increased by 1.3- fold in WG but decreased by 2-fold in WT (Figure S3P). The expression of GCLC reduced 3.5-fold in WG and 1.8-folds in NT whereas it tended decreased further in WT (Figure 3M). The GCLM expression decreased similarly in both WG and WT (Figure S3Q). Glycine, which can either be taken up or produced via serine metabolism, tended to decrease upon metformin treatment, particularly, in WT, 24hour onwards (Figure 3N). While there was no change in glycine level in WG at 48hours, it was found to be depleted in NT and WT compared to respective controls. The serine level showed a decline in WT 12hour onwards whereas it tended to increase in WG 24hour onwards (Figure 3O). At 48hours, the glycine/serine ratio was found to be lower by1.4-fold in WG and 1.5, and 1.9-foldsin NT and WT compared to respective controls (Figure 3P). The expression of SHMT1, responsible for serine-glycine interconversion, reduced 3.7, 3-folds in NT and WT compared to respective controls (Figure 3Q). The expression of SHMT2 reduced 3.3, 2.3-folds in NT and WT compared to respective controls (Figure 3R). Serine and glycine participate in the one-carbon cycle via SHMT1/2 (Figure 4A), that also involves MTR and AHCYL1 genes. The expression of MTR reduced 5, 3.8-folds in NT and WT compared to respective controls (Figure S4A). The expression of AHCYL1 was reduced 2.5-fold in both metformin treatment and glucose deprivation alone (Figure S4B). Expression of genes involved in methionine salvage pathway (Figure 4A) such as MTAP reduced 2.5-fold in WG and 2.9-folds in NT (Figure S4C). Expression of MAT2A, which can also contribute to the adenine pool, was unchanged in NT whereas it was found to be 3.6 and 5-fold downregulated in WG and WT compared to NG and NT, respectively (Figure S4D).

**Figure 4.**
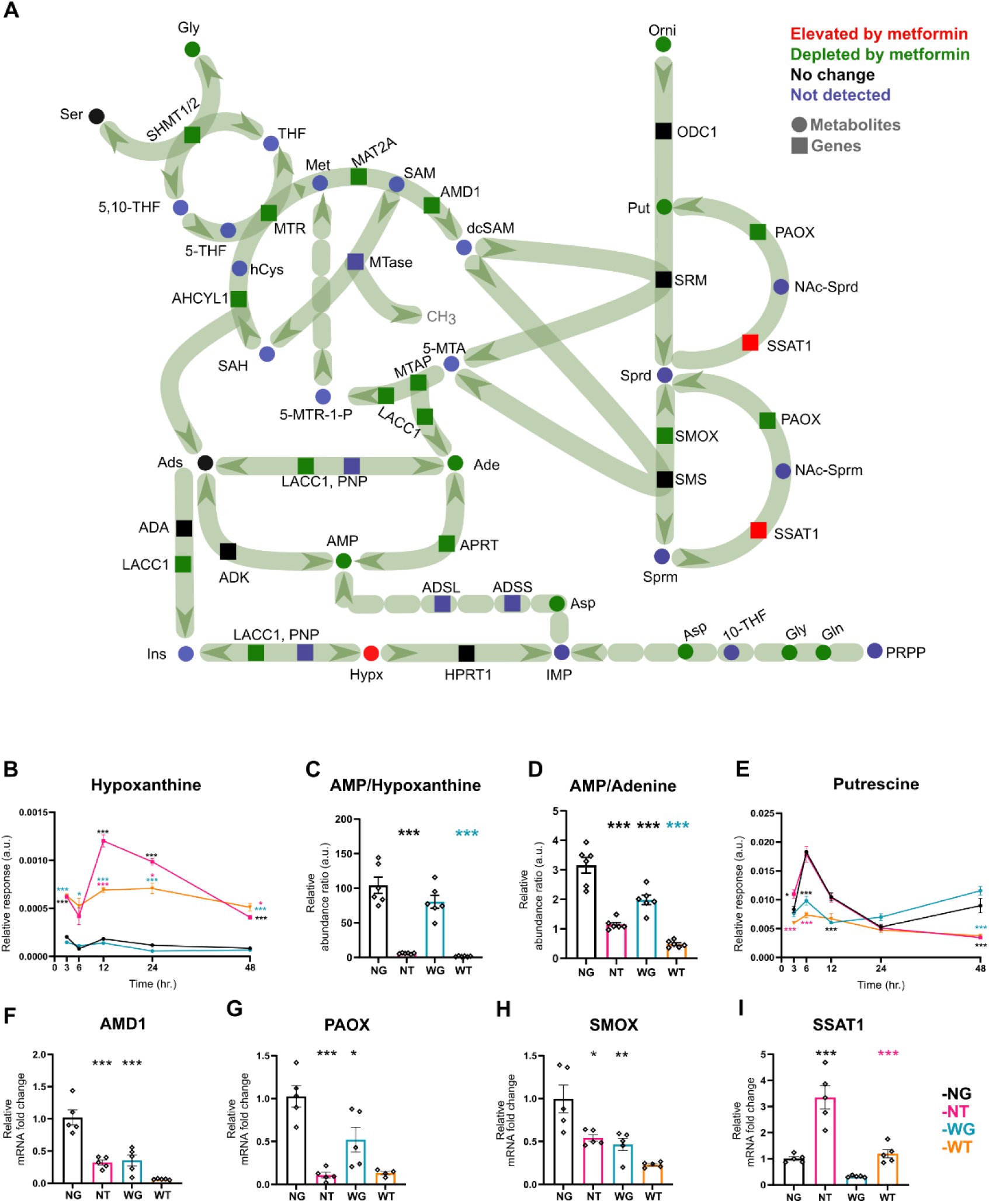
*Effects of metformin on purine and polyamine metabolism. **A.** Schematic depiction of effects of metformin on one-carbon cycle, purine and polyamine metabolism (abbreviations: Ser-Serine, Gly-Glycine, 5,10-THF- 5,10- tetrahydrofolate, THF-Tetrahydrofolate, Met-Methionine, SAM-S- adenosylmethionine, SAH-S-adenosylhomocysteine, hCys-Homocysteine, dcSAM-Decarboxylated S-adenosylmethionine, 5-MTA- 5-methylthioadenosine, 5-MTR-1-P-5-methylthioribose-1-phosphate, Ads-Adenosine, Ade-Adenine, AMP-Adenosine monophosphate, Ins-Inosine, Hypx-Hypoxanthine, IMP-Inosine monophosphate, Asp-Aspartic acid, PRPP-Phosphoribosyl diphosphate, Gln-Glutamine, 10-THF- 10- tetrahydrofolate, Orni-Ornithine, Put-Putrescine, Sprd-Spermidine, Sprm-Spermine, NAc-Sprd-N-acetylspermidine, NAc-Sprm-N-acetylspermine. Used gene abbreviations: ODC1- Ornithine decarboxylase 1, SRM-Spermidine synthase, SMS-Spermine synthase, SMOX-Spermine oxidase, PAOX-Polyamine Oxidase, SSAT1- Spermidine/spermine N1-acetyltransferase 1, AMD1- Adenosylmethionine decarboxylase 1, MAT2A-Methionine adenosyltransferase II alpha, MTR- 5- methyltetrahydrofolate-homocysteine, AHCYL1- S-adenosylhomocysteine hydrolase-like protein 1, MTAP-Methylthioadenosine Phosphorylase, SHMT1/2- Serine hydroxymethyltransferase 1/2, LACC1- Laccase Domain Containing 1, PNP-Purine nucleoside phosphorylase, ADA-Adenosine deaminase, ADK-Adenosine kinase, APRT-Adeninephosphoribosyltransferase,HPRT1-hypoxanthinephosphoribosyl-transferase 1, ADSS-adenylosuccinate synthase, ADSL-Adenylosuccinate lyase). The trend plots for longitudinal changes in relative abundances of hypoxanthine (**B**) and putrescine (**E**) is shown along with changes in AMP/hypoxanthine (**C**) and AMP/adenine (**D**) ratios at 48 hours. Effects on expression of adenosylmethionine decarboxylase 1 (AMD1, **F**), polyamine oxidase (PAOX, **G**), spermine oxidase (SMOX, **H**), and spermidine/spermine N1-acetyltransferase 1 (SSAT1, **I**) genes are also shown. All data presented as mean ± SEM with n = 6 for metabolites and 5 for genes. One-way ANOVA with Bonferroni post-hoc test for multiple comparisons was used to test statistical significance. P values <0.05(*), <0.002(**), <0.001(***) are indicated and color-coded for pair-wise comparisons (black: comparison with NG, cyan: comparison with WG and pink: comparison with NT).*

### Effects of metformin on purine and polyamine metabolism

Glutamine, glycine, aspartic acid and one-carbon units are required for *de novo* synthesis of purines, which can also be synthesized via salvage pathway from bases like hypoxanthine or adenine (Figure 4A). The hypoxanthine level was elevated 3hour onwards in both NT and WT while it remained unchanged in NG and WG (Figure 4B). At 48hours, the AMP/hypoxanthine ratio was 18 and 57-folds lower whereas AMP/adenine ratio was 2.7 and 4.1-folds lower, respectively, in NT and WT compared to respective controls (Figure 4C, D). But they were unaffected in WG. While, HPRT expression reduced 2-fold upon glucose deprivation (Figure S4E), APRT expression was reduced by 1.8-fold upon metformin treatment and glucose deprivation alone (Figure S4F). LACC1, a multifunctional gene that can contribute to purine salvage (Figure 4A), reduced 7.7-fold in WG and 2.6-fold in NT while it tended to decrease further in WT (Figure S4I). Expression of ADA was also downregulated upon metformin treatment and glucose deprivation (Figure S4G). However, there was no difference between NT and WT for HPRT, APRT, ADA or ADK (Figure S4E-H). The one-carbon metabolism also contributes to the polyamine biosynthesis (Figure 4A), which is initiated by conversion of ornithine to putrescine via ODC1. While ornithine was found to gradually increase in WG, it showed a tendency of decrease in NT at 48hours and was depleted in WT compared to WG after 24hour (Figure S4J). Putrescine was found significantly depleted in WT at 3hour itself. At 48hours, the putrescine level in NT and WT were found to be similar and significantly depleted compared to respective controls (Figure 4E). Expressions of AMD1, PAOX, SMOX involved in polyamine biosynthesis and interconversion were significantly attenuated by metformin treatment (Figure 4F-H) whereas ODC1, SRM and SMS were unaffected (Figure S4K-M). AMD1, which connects one-carbon and polyamine metabolism, reduced 2.8, 6.7- folds in NT and WT compared to respective controls. PAOX was significantly downregulated in both NT and WT by 6.2-fold and 4.5-fold compared to respective controls. SMOX expression reduced 2- and 1.8- in WG and NT, while it declined further in WT. On the other hand, expression of SSAT1, involved in polyamine acetylation, was 3.3 and 4-fold upregulated in NT and WT compared to respective controls (Figure 4I).

### Effect of metformin on cMyc expression and its correlation with metabolic and gene expression signatures

The BrdU incorporation was found to be very highly correlated (0.6<Pearson correlation coefficient <-0.6) with several metabolites (Figure 5A, Table S8), and genes (Figure 5B, Table S9) that were found to be affected by metformin treatment at 48hours. While TCA cycle intermediates, EAA as well as NEAAs, pantothenic acid, taurine, adenine, AMP and putrescine were positively correlated, saturated and unsaturated fatty acids, monopalmitin and hypoxanthine were negatively correlated. EAAs formed a highly correlated cluster and were also positively correlated with serine, glycine, ornithine, putrescine and pantothenic acid while these were negatively correlated with fatty acids and hypoxanthine (Figure 5A, Table S10).

**Figure 5.**
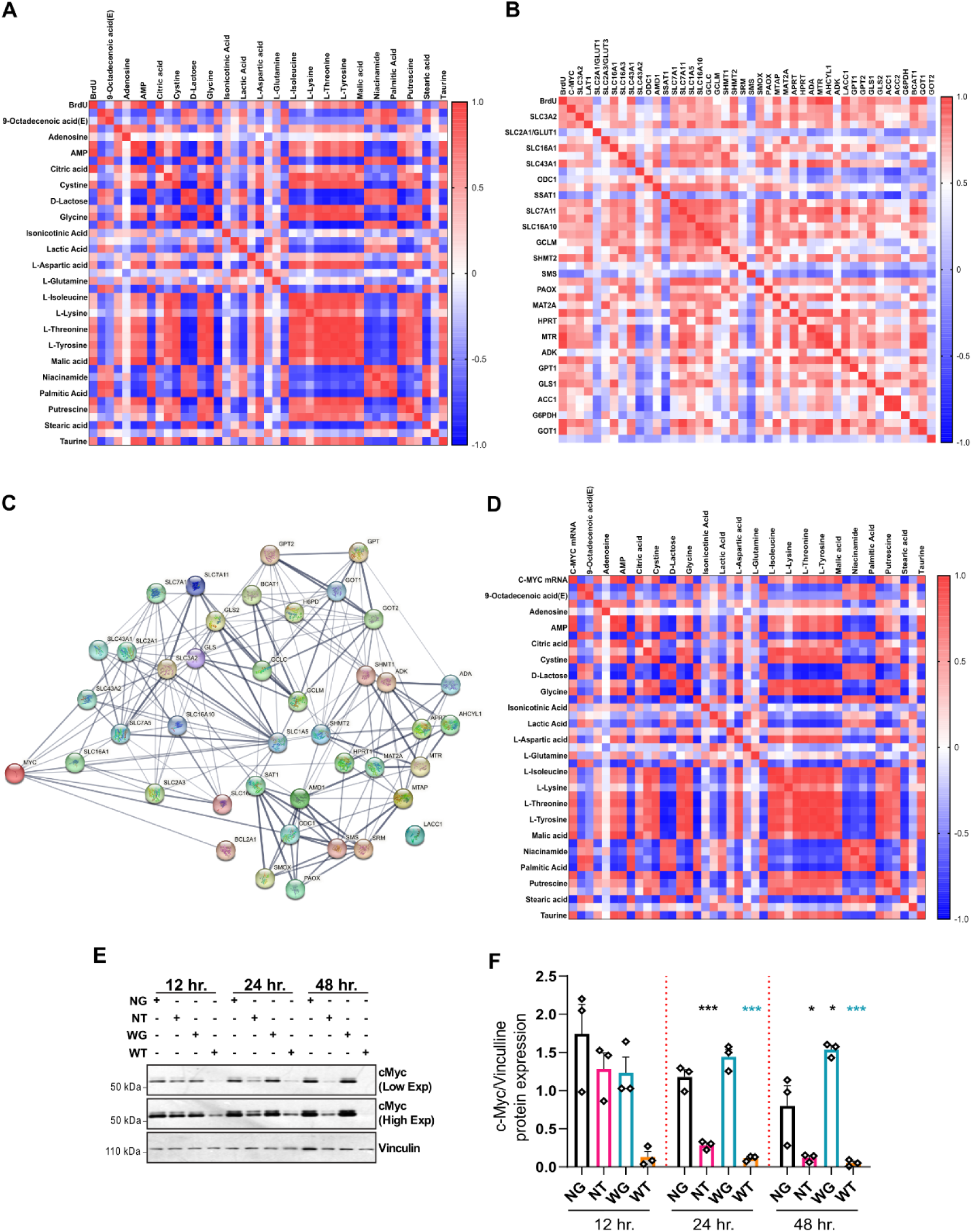
*Correlation between proliferation, metabolites, gene expression signatures and cMyc expression. Correlation matrix plot showing (**A**) association between BrdU incorporation and metabolite levels, (**B**) BrdU incorporation and gene expression levels at 48 hours. (**C**) Association between c-MYC and genes studied here obtained via STRING network analysis (https://string-db.org/cgi/about). **D.** Correlation matrix plot showing association between expression of cMYC gene and metabolite levels at 48 hours. Scale bar on the left side of the plot is indicative of the Pearson correlation coefficient (r). Red and blue color indicates positive or negative correlations respectively. **E.** Immunoblot showing effect of metformin treatment and glucose deprivation on cMyc protein expression at 12-, 24-, and 48-hours post-treatment. Vinculin was used as loading control. **F.** Densitometric quantification of the immunoblot signals from **E** (data presented as mean ± SEM with n = 3). One-way ANOVA with Bonferroni post-hoc test for multiple comparisons was used to test statistical significance. P-value <0.05(*), <0.002(**), <0.001(***) and color-coded for pairwise comparisons. Black stars: comparison with NG, pink stars: comparison with NT, cyan stars: comparison with WG.*

On the other hand, amino acid transporters (SLC1A5, SLC7A11, SLC16A1, SLC3A2, LAT1, SLC7A1, SLC16A10, SLC43A1), amino acid and central carbon metabolism genes (BCAT1, GOT1, GPT1, GPT2, GLS1, GLS2, ACC1, ACC2), one-carbon and GSH biosynthesis genes (SHMT1, SHMT2, AHCYL1, MTR, MTAP, GCLC), polyamine pathway and nucleotide salvage genes (AMD1, SMOX, PAOX, MTAP, ADA, LACC1, HPRT, APRT)showed significant positive correlations with BrdU incorporation (Figure 5B,Table S9). Given that several genes (Figure5C) and pathways, which were found to be affected upon metformin treatment, are known to be regulated by cMyc, the cMYC expression was analyzed at both mRNA (Figure S4N) and protein levels. cMYC mRNA level was not only highly correlated with BrdU incorporation (Figure 5B) but also tightly correlated with changes in metabolite abundances with a correlation pattern similar to that for BrdU (Figure 5D, Table S11). cMYC was also significantly correlated with the expression of amino acid transporters and genes involved in amino acid and central carbon metabolism, one-carbon metabolism and GSH biosynthesis, polyamine pathway and nucleotide salvage (Figure 5B, Table S12). While there was no change in *cMyc* protein expression upon glucose deprivation alone, it was found to be attenuated at 24hours in glucose-supplemented media and preceded the reduction of proliferation (Figure 5E-F). It was attenuated 12hour onwards upon metformin treatment in glucose-deprived media, which coincided with the reduction in BrdU incorporation.

### Reversibility of the effect of metformin on proliferation, metabotype and cMyc expression

At 24hours, when there was no change in viability but a significant reduction in BrdU incorporation in WG and WT (FigureS5A-B) the reversibility of the effect of metformin was tested by rescue experiment as depicted in Figure 6A. It was observed that at 48hours after media replacement, there was no difference between BrdU incorporation (Figure 6B) of 3B and 4D, which were ∼12 and ∼14-fold lower, respectively, than those in control media (1, 2A, 3A, 4B). The BrdU incorporation recovered fully in 4B where WT media was replaced with NG and was no different from the control (1). Even upon removal of metformin alone (4A), 10- and 3.7-fold increase in BrdU incorporation occurred compared to cells in WT (4D) or those continuing with metformin (4C), respectively, and it was found at 70% of the control. But, when glucose was replenished and metformin was continued (4C), there was no significant proliferative recovery. This indicated that while metformin is the dominant driver of the decrease in proliferation, the effect is quite reversible upon removal of metformin. The XTT assay (Figure S5C) also showed near-complete recovery in viability in 4B.

**Figure 6.**
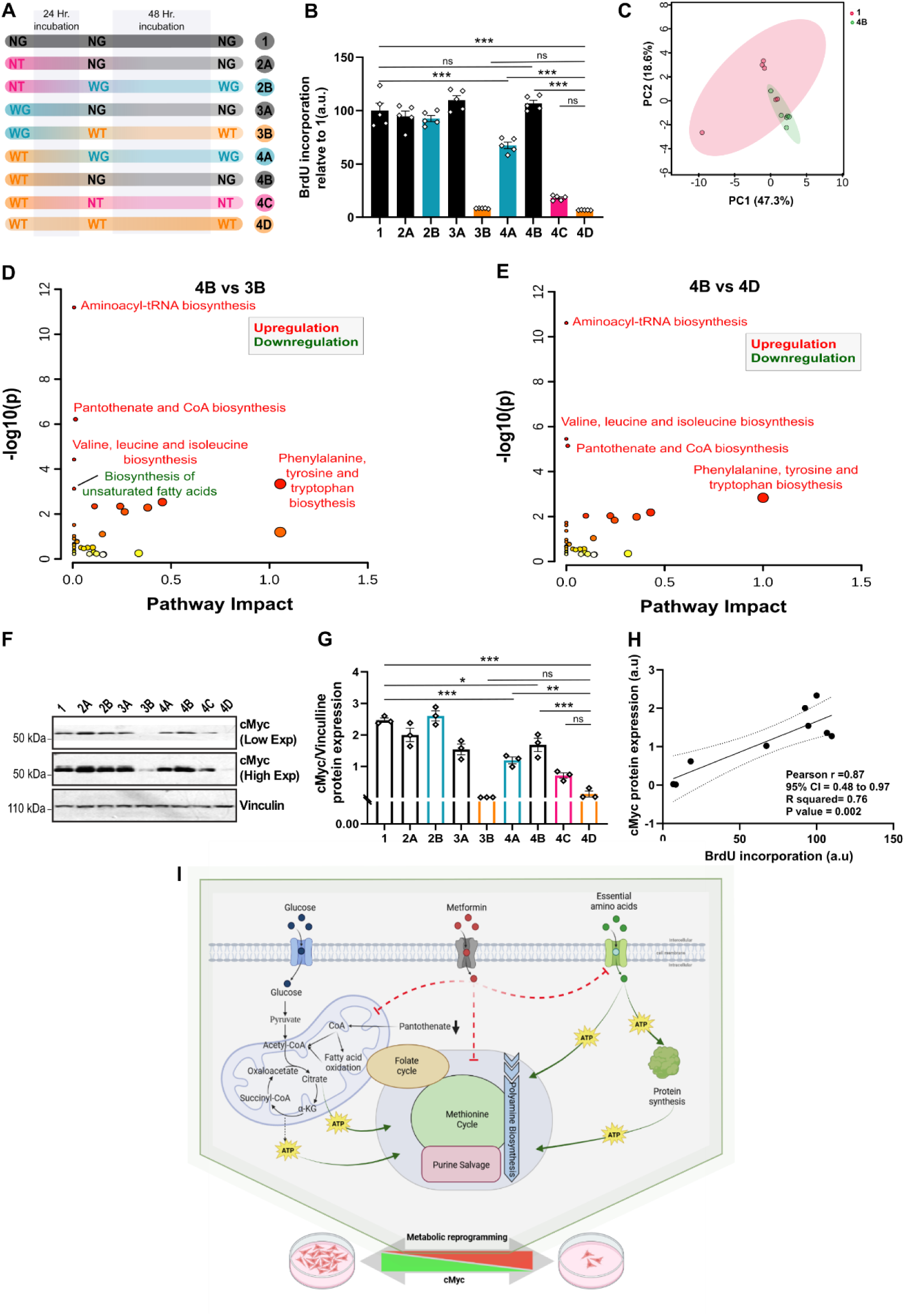
*Reversal of proliferation, metabolic signature, and cMyc expression upon metformin withdrawal. A. Experimental design of the rescue experiment. cells were cultured under aforementioned conditions for 24 hours followed by media change and cultured for another 48hours as described below. The control (NG) group was continued with NG media (1). For NT group, either metformin was removed by replacing with NG media (2A) or exposed to only glucose deprivation using WG media (2B). For WG group, either glucose was replenished with NG media (3A) or exposed to metformin treatment in absence of glucose using WT media (3B). For WT group, either metformin was removed using WG media (4A) or glucose was replenished using NT media (4C) or both were done together using NG media (4B) or continued with WT media (4D) as control. B. BrdU incorporation in rescue experiment. C. 2D scores scatter plots from principal components analysis (PCA) showing no difference in overall metabotype of control cells (1) and those rescued from WT with NG media (4B). Differentially regulated metabolic pathways in 3B (D) and 4B (E) compared to 4D indicating reversal of metabolic derangements upon metformin withdrawal. F. Immunoblot showing cMyc expression levels in different groups in rescue experiment. Vinculin was used as loading control. G. Densitometric quantification of the immunoblot signals from F (data presented as mean ± SEM with n = 3). One-way ANOVA with Bonferroni post-hoc test for multiple comparisons was used to test statistical significance. P-value <0.05(*), <0.002(**), <0.001(***) are indicated above lines between respective pairs. H. Correlation between average cMyc expression and BrdU incorporation in rescue experiments. I. The schematic representation of the glucose-dependent and independent metabolic derangements contributing to reversible anti-proliferative effect of metformin and its association with cMyc expression in liver cancer cells.*

The PCA scores plot of metabolic signatures showed that while there was no difference between 3B and 4D (Figure S5D), there was also no separation between 4B and 1 (Figure 6C) indicating a recovery of metabolic signature was associated with rescue of the proliferative phenotype. The comparison of metabolic signature between 4B and 3B or 4D showed upregulation or downregulation of same metabolic pathways (Figure 6D, Table S13 and Figure 6E, Table S14) that were found to be, respectively, downregulated or upregulated upon metformin treatment (Figure 1I-J) indicating reversal of derangements. Interestingly, the differential signatures between 4B and 4C were also quite similar (Figure S5E, Table S15) indicating these pathways to be reversibly metformin-responsive.

Similar to that observed for the BrdU incorporation, 3B and 4D were found to have lowest and similar *cMyc* expression whereas there was no difference between *cMyc* levels in 4B, 2A or 3A that were rescued with NG media (Figure 6F, G). Figure 6H showed strong correlation (r = 0.87, P < 0.002) between *cMyc* protein level and BrdU incorporation.

## Discussion

This is the first study that examined longitudinal effect of metformin on liver cancer cells in tandem with metabolic signature. The 2.5mM dose, which is less than the reported IC_50_ value (7mM) for HepG2^16^ was chosen to simulate the non-cytotoxic effect of metformin since there is no direct evidence of any significant cytotoxic effect of metformin alone on tumors *in vivo*. The anti-proliferative effect of metformin at this concentration is in line with well-documented tumor growth inhibition by metformin via inhibition of proliferation. This study showed that the anti-proliferative effect of metformin and the accompanying metabolic derangements involve novel non-central carbon pathways. Interestingly, it also showed these are reversible and associated with *cMyc* downregulation irrespective of presence of glucose.

Glycolysis and TCA cycle-mediated glucose metabolism is the primary source of carbon and energy (ATP) in cells. In absence of glucose, cancer cells can utilize glutamine as an alternative nutrient to drive anaplerotic TCA cycle^23^. This was reflected by depletion of glutamine in absence of glucose. Further reduction of glutamine upon metformin treatment is in line with earlier observation in oesophageal squamous cell carcinoma^29^. However, in presence of metformin, there was a marked decrease in TCA cycle intermediates indicating compromised mitochondrial function, which was also shown in earlier studies in both cellular models and patient samples^15,16,30^. This might force cells to be more reliant on glycolysis, which was indicated by increase in lactate in presence as well as in absence of glucose. In glucose-deprived media, the pyruvate for lactate production may come from alanine, resulting in gradual decrease in alanine observed in WT. The compromised TCA cycle upon metformin treatment might also explain the decrease in aspartate - a product of transamination of TCA intermediate oxaloacetate. Non-utilization via TCA cycle, however, would increase glutamate as was observed here and consistent with earlier reports on the effect of metformin^31^.

Fatty acids can be produced from citrate whereas their oxidation, which primarily occurs in mitochondria, can also supply carbons to TCA cycle. Fatty acid oxidation (FAO) was found to be elevated in cancer stem cells that contribute to therapeutic resistance and relapse^32–35^. Increase in fatty acids even with decreased citrate and genes essential for *de novo* fatty acid synthesis (ACC1 and ACC2) indicated reduced FAO due to mitochondrial toxicity. Interestingly, pantothenic acid, which is required for synthesis of CoA required for TCA cycle and fatty acid metabolism, was found to be depleted much earlier than increase in fatty acids and decrease in proliferation in metformin treated cells. This could compromise TCA cycle, FAO^36^as well as other pathways dependent on CoA such as essential amino acid metabolism^37^. The abundances of a number of unsaturated fatty acids including linoleic, arachidonic acid, which were elevated upon metformin treatment in plasma samples^38,39^, were elevated here as well. An earlier study showed that NAD^+^, which is required for fatty acid desaturation, was elevated upon metformin treatment in HepG2 cells^40^. Studies involving patient samples and MCF7 cells showed that metformin reduced FAO and increased FA desaturation in AMPK-independent manner^41^. However, the longitudinal impact of metformin on these metabolic pathways in liver cancer cells is reported here for the first time irrespective of glucose supplementation.

Serine contributes not only to ATP production under glucose deprivation but also to *de novo* synthesis of nucleotides and polyamines via one-carbon cycle in cancer^42^. There was a decrease in serine transporter (SLC1A5) expression and serine abundance, particularly, under glucose deprivation. SHMT1 and SHMT2, which convert it to glycine to drive the one-carbon cycle and are *cMyc* targets^43^, were significantly downregulated with concomitant decrease in glycine upon metformin treatment irrespective of glucose supplementation. In addition, other genes involved in one-carbon metabolism (AHCYL1, MTR, MAT2A) were also downregulated and indicated that metformin widely targeted one-carbon metabolism. While this was shown in breast cancer cells^10^, this study reports such effect in liver cancer cells for the first time. Cysteine was also found to be depleted upon metformin treatment along with decrease of expression of SLC7A11 and SLC3A2, which combines to form the xCT antiporter system^44^. These combined with dearth of energy and decrease in expression of GCLC and GCLM to decrease in total glutathione level, particularly, under glucose deprivation.

Polyamines are crucial for proliferation and their acetylation was shown to arrest it^45,46^. Metformin was found to decrease both putrescine and putrescine/ornithine ratio indicating reduced flux towards polyamine biosynthesis although the ODC1 expression was unchanged. However, AMD1, which is a *cMyc* target and essential for higher polyamine biosynthesis, was downregulated with concomitant increase in SSAT1, which leads to acetylation, upon metformin treatment irrespective of glucose level. This can decrease free polyamines to inhibit to proliferation. Such derangement in polyamine metabolism upon metformin treatment was never reported in liver cancer cells although metformin-induced attenuation of putrescine was recently reported in colorectal cancer xenografts^47^.

Cancer cells have been found to utilize branched-chain amino acids not only for regeneration of glutamine via BCATs, but also for energy generation via mitochondrial catabolism^48^. For the first time, we report that metformin reduces expression of transporters of branched chain as well as other EAAs leading to their depletion in cancer cell. It also reduced expression of BCAT1, which has been found to be upregulated and associated with poor prognosis in several cancers^48,49^. Along with mitochondrial dysfunction and reduction in pantothenate, these would compromise glutamine replenishment as well as energy metabolism from essential amino acids. In addition, low EAAs would also hinder translational machinery towards protein synthesis, which is essential for proliferation. These unravel a hitherto undiscovered mechanism of metformin-induced anti-proliferative effect via depletion of essential amino acids in cancer cells.

Proliferation requires nucleotides for DNA and RNA synthesis. Metformin treatment not only decreased DNA synthesis (BrdU incorporation) but also resulted in reduction in level of several mRNAs. Given that levels of glutamine, aspartate, glycine as well as one-carbon metabolism were downregulated in conjunction with energy crisis upon metformin treatment, its intuitive that *de novo* synthesis of nucleotides would be difficult. Cancer cells have been shown to resort to nucleotide salvage, which is associated with resistance^50,51^, and targeting nucleotide salvage pathways have been found to sensitize breast cancer cells to antimetabolite therapy^52^. Salvage pathway, being less energy and resource demanding, is important to maintain proliferation under nutrient limiting condition. Our results indicated that metformin treatment tended to reduce expression of purine salvage genes as well as AMP/adenine ratio. It also increased hypoxanthine concentration well ahead of the observed decrease in proliferation. Hypoxanthine is well known to act as substrate for nucleotide salvage to promote proliferation^53^. These indicated that attenuation of salvage pathway by metformin might further contribute to inhibition of proliferation.

Taken together, the simultaneous reduction in amino acids, pantothenic acid, TCA cycle and fatty acid oxidation impaired the energy and redox balance upon metformin treatment. The decrease in TCA intermediates or amino acids and increase in fatty acids were more pronounced in WT indicating that the energy crisis got exaggerated under glucose deprivation. Thus, our results provide further biochemical basis for the synergistic effect of hypoglycaemia and metformin treatment that was found to reduce tumor load in mouse model of colorectal cancer^54^. The derangement of fatty acid desaturation was found to be uniquely associated with metformin treatment irrespective of glucose deprivation. It also impaired the one-carbon cycle to attenuate biosynthesis of polyamines, purines as well as purine salvage pathway. These were co-ordinately associated with downregulation of essential amino acid transport and *cMyc* expression to result in a reduction in proliferation as shown in Figure 6I. cMyc, which is a master regulator of proliferation, has earlier been shown to control EAA transport via a positive feed-forward loop involving *Lat1* and *Slc3a2*^55,56^. In addition, cMyc is a well know regulator of ribosome biogenesis and protein synthesis^57^. Thus, its downregulation in tandem with coordinate derangement of aforementioned metabolic pathways involved in energy production, one-carbon cycle, polyamine metabolism, protein and nucleic acid synthesis would restrict the proliferative capacity of cancer cells. On the other hand, cancer cells were shown to reduce *cMy*c expression as a survival strategy under nutritional stress^58^. Earlier studies also showed downregulation of *cMyc* upon metformin treatment via both post-transcriptional and proteasomal mechanisms^59,60^.

This begged the question whether such reduction in *cMyc* expression and concomitant metabolic derangements in cancer cells are irreversible. The rescue experiments showed that these changes under non-cytotoxic metformin concentration were actually reversible. The complete reversal of changes in central carbon metabolites and fatty acids indicated that the observed mitochondrial toxicity was also reversible. Given that the physiological concentrations of metformin are lower than the concentration used here, these observations invite further examination of reversibility of metformin-induced anti-cancer effects *in vivo*. If validated, that would indicate that metformin withdrawal in patients or those at risk of developing liver cancer (e.g., diabetics, NAFLD) may increase the risk of disease progression. On the other hand, pathways like serine, EAA catabolism, fatty acid oxidation and nucleotide salvage are sparingly used by normal cells. Thus, development of metformin analogues that preserve/improve inhibitory effect on these pathways but sans deleterious effects on central carbon metabolism, may allow dose escalation without significantly affecting other cells. The fact that downregulation of these pathways, often utilized by therapy-resistant and stem-like cancer cells, were found to be glucose-independent, indicates to the possibility of using metformin/analogues to target such cells. Targeting multiple complimentary pathways based on genetic and metabolic vulnerability of cancer cells has been shown to trigger synthetic lethality^61–63^. Thus, metformin/analogues may be coupled with therapeutic agents targeting complementary pathways (e.g., glycolysis, fatty acid desaturation, *de novo* nucleotide synthesis) and/or intermittent fasting to induce synergistic or even synthetic lethality to increase preventive and therapeutic efficacy and personalize cancer treatment. Although several questions including the precise molecular mechanism underlying *cMyc* downregulation and metabolic derangements remain subject to further investigations, our study not only elucidates the temporal evolution of biochemical landscape leading to metformin-induced anti-proliferative effects but also provides a glimmer of hope to exploit some of these unique pathways to develop novel preventive and therapeutic strategies.

## Conclusions

This study showed that metformin can exert its effect on liver cancer cells even without eliciting cytotoxicity via attenuation of proliferation irrespective of glucose level. Apart from targeting TCA cycle, metformin was found to attenuate essential amino acid uptake and metabolism, fatty acid oxidation, one-carbon metabolism, polyamine metabolism, and purine salvage pathways to concomitantly attenuate proliferation of cancer cells. Increase in fatty acid desaturation was found to be a novel signature, whose role needs further investigation. Such attenuation of proliferation was found to be co-ordinately associated with downregulation of cMyc expression. These indicate to the possibility of use of metformin or its analogues in tandem with intermittent fasting and drugs targeting complimentary pathways in treatment of liver cancer. However, the anti-proliferative effect and associated changes in metabolic derangements and cMyc expression were found to be reversible irrespective of presence of glucose indicating that metformin withdrawal may have consequences on disease progression and warrants further studies.

## Supporting Information

Supplementary figures including figure legends (pdf). List of authentic standards used for metabolite identification (Table S1, xlsx). List of primers used for mRNA expression analysis (Table S2, xlsx). Table of annotated metabolites including retention time, selected ions, match score, and level of authentication (Table S3, xlsx). Table of enriched pathway in NT, and WT group compared to NG, and WG respectively (Table S4-5, xlsx). Table of overlapping compounds in NT, WG, and WT groups as found by Venn diagram analysis (Table S6, xlsx). Table of pathways commonly affected in both NT and WT (Table S7, xlsx). Table of metabolites, and genes correlated with BrdU incorporation at 48hours of treatment (Table S8-10, xlsx). Table of metabolites, and genes correlated with cMYC expression at 48hours of treatment (Table S11-12, xlsx). Table of rescued pathway in 4B group compared to 3B, 4D, and 4C (Table S13-15, xlsx). Extended supporting data of immunoblots (pdf).

## Authors’ Contributions

**S.R.I.:** Conceptualization, methodology, validation, formal analysis, investigation, data curation, writing - original draft, and visualization. **S.K.M:** Conceptualization, writing - review & editing, supervision, project administration, and funding acquisition.

## Notes

### Availability of data and materials

All data needed to evaluate the conclusions in the paper are present in the paper and/or the Supplementary Materials. The raw metabolomic data have been deposited to the Metabolights database (https://www.ebi.ac.uk/metabolights/, ID: MTBLS7760).

## Acknowledgements

Prof. Debashis Mukhopadhyay for HepG2 cell line and Mr. Sebabrata Maity for helping with immunoblotting.

## Abbreviations

ACC1: Acetyl CoA carboxylase alpha aka ACACA
ACC2: Acetyl CoA carboxylase beta aka ACACB
ADA: Adenosine deaminase
ADK: Adenosine kinase
AHCYL1: S adenosylhomocysteine hydrolase like protein
AMD1: Adenosylmethionine decarboxylase
AMPK: AMP-activated protein kinase
ANOVA: Analysis of Variance
APRT: Adenine phosphoribosyltransferase
BCAA: Branched-chain amino acids
BCAT1: Branched chain amino acid transaminase
BrdU: Bromodeoxyuridine
cMyc: Cellular myelocytomatosis oncogene
CoA: Coenzyme A
DMEM: Dulbecco’s Modified Eagle Medium
EAA: Essential Amino Acids
GCLC: Glutamate Cysteine Ligase Catalytic Subunit
GCLM: Glutamate Cysteine Ligase Modifier Subunit
GLS1: Glutaminase
GLS2: Glutaminase
GOT1: Glutamic oxaloacetic transaminase 1 cytosolic
GOT2: Glutamic oxaloacetic transaminase 2 mitochondrial
GPT1: Glutamic pyruvic transaminase
GPT2: Glutamic pyruvic transaminase
HPRT1: Hypoxanthine phosphoribosyltransferase 1
LACC 1: Laccase Domain Containing
LAT1: L type amino acid transporter 1 aka SLC7A5
LKB: Liver Kinase B aka STK11.
MAT2A: Methionine adenosyltransferase II alpha
MTAP: Methylthioadenosine Phosphorylase
MTR: 5 methyltetrahydrofolate homocysteine methyltransferase
NAFLD: Non-alcoholic fatty liver disease
NEAA: Non-essential Amino Acids
ODC1: Ornithine decarboxylase
PAOX: Polyamine Oxidase
SHMT1: Serine hydroxymethyltransferase 1
SHMT2: Serine hydroxymethyltransferase 2
SLC16A1: Solute Carrier Family 16 Member 1
SLC16A10: Solute Carrier Family 16 Member 10
SLC16A3: Solute Carrier Family 16 Member 3
SLC1A5: Solute Carrier Family 1 Member 5 aka ASCT2 Glutamine Transporter
SLC2A1: Solute carrier family 2 facilitated glucose transporter member 1 aka GLUT1
SLC2A3: Solute carrier family 2 facilitated glucose transporter member 3 aka GLUT3
SLC3A2: Solute carrier family 3 member 2 aka CD98
SLC43A1: Solute Carrier Family 43 Member1
SLC43A2: Solute Carrier Family 43 Member 2
SLC7A1: Solute Carrier Family 7 Member 1
SLC7A11: Solute Carrier Family 7 Member 11
SMOX: Spermine oxidase
SMS: Spermine synthase
SRM: Spermidine synthase
SSAT1: Spermidine /spermine N1 acetyltransferase 1 aka SAT1
TCA: Tricarboxylic acid cycle
XTT: Methoxynitrosulfophenyl-tetrazolium carboxanilide.

