## Supplementary material for "Identification of glucose-independent metabolic pathways associated with anti-proliferative effect of metformin, their coordinate derangement with cMyc downregulation and reversibility in liver cancer cells"

### Supplementary Figure Legends

**Figure S1.** Dose- and time-dependent effect of metformin and glucose deprivation on cell fate and metabolism.

**A.** Dose and time-dependent (3-48 hours) effect of metformin and glucose restriction on cell viability assessed by XTT-cell viability assay. The absorbance at each time point was normalized with respect to mean absorbance in the NG group at that time point (data presented as mean ± SEM, n = 6). **B.** Dose and time-dependent effect of metformin and glucose restriction on cell proliferation assessed by BrdU-incorporation assay. The absorbance at each time point was normalized with respect to mean absorbance in the NG group at that time point (data presented as mean ± SEM, n = 4). **C.** Dose and time-dependent cytotoxic effect of metformin and glucose restriction on HepG2 cells assessed by CellTox Green cytotoxicity assay (data are mean ± SEM; n = 5). One-way ANOVA with Bonferroni post-hoc test for multiple comparison was used to test statistical significance. P values <0.05(*), <0.002(**), <0.001(***) are indicated and color-coded for pair-wise comparisons (black: comparison with NG, cyan: comparison with WG and pink: comparison with NT).

**Figure S2.** Effect of metformin in on metabolic signatures

**A.** Venn diagram of differentially expressed metabolites (fold change > 1.5 and P < 0.05) in NT, WG, and WT at 48 hours. of treatment with 2.5mM metformin compared to NG. Numbers indicate the number of differentially altered metabolites shared by experimental groups. Diagram was made using an online tool (https://bioinformatics.psb.ugent.be/webtools/Venn/). B. Alanine/glutamate ration at 48 hours (data presented as mean ± SEM, n = 6). One-way ANOVA with Bonferroni post-hoc test for multiple comparison was used to test statistical significance. P values <0.05(*), <0.002(**), <0.001(***) are indicated and color-coded for pair-wise comparisons (black: comparison with NG, cyan: comparison with WG and pink: comparison with NT).

**Figure S3.** Effect of metformin on amino acids, glutathione and amino acid transporters.

**A-G, L and P.** Relative abundance of isoleucine, leucine, valine, threonine, phenylalanine, tyrosine, tryptophan, glutamine and glutathione at 48hrs along with trend plots for longitudinal changes in relative abundances of glutamic acid (**J**), glutamine (**K**) and taurine (**M**) (data presented as mean ± SEM, n = 6). Expression of solute carrier family 3 member 2 (SLC3A2, **H**), L-type amino acid transporter 1 (LAT1, **I**), Solute Carrier Family 16 Member 10 (SLC16A10, **N**), Solute Carrier Family 7 Member 1 (SLC7A1, **O**), and Glutamate-Cysteine Ligase Modifier Subunit (GCLM, **Q**) genes at 48hours (data presented as mean ± SEM; n = 5). One-way ANOVA with Bonferroni post-hoc test for multiple comparison was used to test statistical significance. P values <0.05(*), <0.002(**), <0.001(***) are indicated and color-coded for pair-wise comparisons (black: comparison with NG, cyan: comparison with WG and pink: comparison with NT).

**Figure S4**. Effect of metformin on purine and polyamine metabolism.

**A-I, K-N.** Expression of 5-methyltetrahydrofolate-homocysteine (MTR), S-adenosylhomocysteine hydrolase-like protein 1 (AHCYL1), Methylthioadenosine Phosphorylase (MTAP), Methionine adenosyltransferase II alpha (MAT2A), Hypoxanthine phosphoribosyltransferase 1 (HPRT), Adenine phosphoribosyltransferase (APRT), Adenosine deaminase (ADA), Adenosine kinase (ADK), Laccase Domain Containing 1 (LACC1), Ornithine decarboxylase 1 (ODC1), Spermidine synthase (SRM), Spermine synthase (SMS), Cellular myelocytomatosis oncogene (C-MYC) genes at 48hours (data presented as mean ± SEM.; n = 3-5). **J.** Trend plots for longitudinal changes in relative abundance of ornithine (data presented as mean ± SEM, n = 6). One-way ANOVA with Bonferroni post-hoc test for multiple comparison was used to test statistical significance. P values <0.05(*), <0.002(**), <0.001(***) are indicated and color-coded for pair-wise comparisons (black: comparison with NG, cyan: comparison with WG and pink: comparison with NT).

**Figure S5.** Reversibility of metabolic signature in rescue experiments.

**A.** Cell viability by XTT-assay at 24 hours of treatment (mean ± SEM, n = 6). **B.** Cell proliferation by BrdU-incorporation assay at 24 hours of treatment (mean ± SEM, n = 4). **C.** XTT activity in different groups in rescue experiments showing reversal of cell viability upon metformin withdrawal (data mean ± SEM, n = 6). One-way ANOVA with Bonferroni post-hoc test for multiple comparisons was used to test statistical significance. P-value <0.05(*), <0.002(**), <0.001(***) are indicated above lines between respective pairs. **D.** 2D scores scatter plot from PCA showing indistinguishable metabotype of 3B and 4D. **E.** Metabolic pathway analysis showing recovery of three indicated metabolic pathways in 4B compared to 4C.

Figure S1

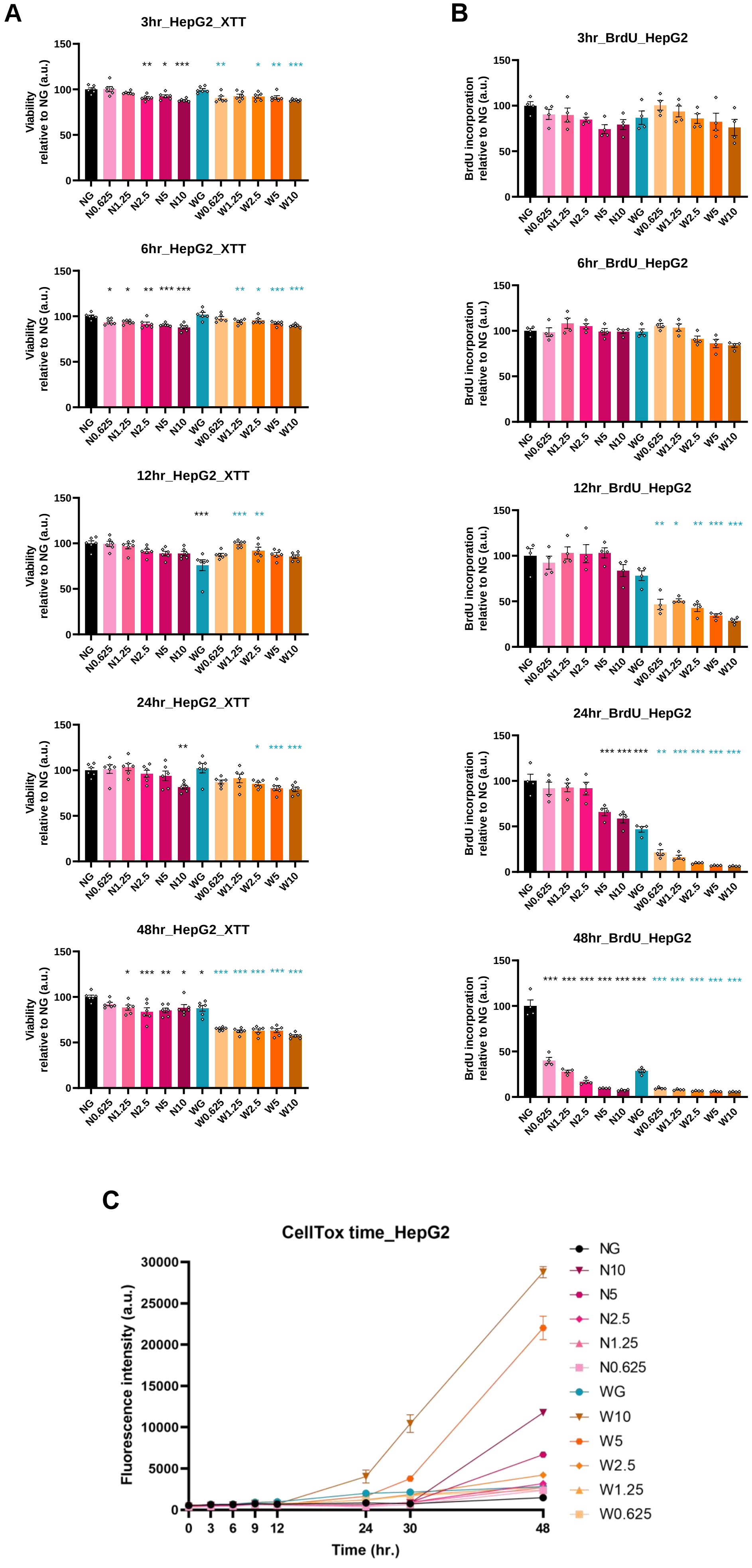
A
3hr_HepG2_XTT
6hr_HepG2_XTT
12hr_HepG2_XTT
24hr_HepG2_XTT
48hr_HepG2_XTT
Viability relative to NG (a.u.)
NG N0.625 N1.25 N2.5 N5 N10 WG W0.625 W1.25 W2.5 W5 W10
B
3hr_BrdU_HepG2
6hr_BrdU_HepG2
12hr_BrdU_HepG2
24hr_BrdU_HepG2
48hr_BrdU_HepG2
BrdU incorporation relative to NG (a.u.)
C
CellTox time_HepG2
Fluorescence intensity (a.u.)
Time (hr.)
NG
N10
N5
N2.5
N1.25
N0.625
WG
W10
W5
W2.5
W1.25
W0.625

Figure S2

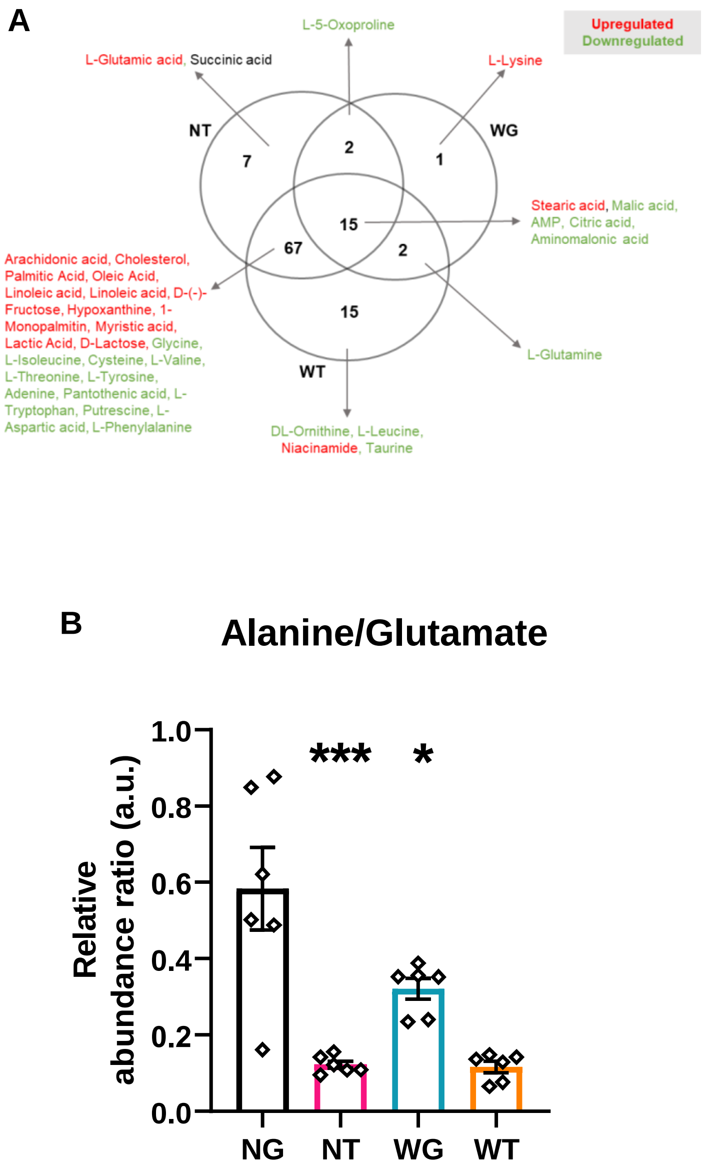
A
Upregulated
Downregulated
L-5-Oxoproline
L-Glutamic acid, Succinic acid
L-Lysine
NT
WG
WT
7
2
1
15
67
2
15
Stearic acid, Malic acid, AMP, Citric acid, Aminomalonic acid
Arachidonic acid, Cholesterol, Palmitic Acid, Oleic Acid, Linoleic acid, Linoleic acid, D-(-)-Fructose, Hypoxanthine, 1-Monopalmitin, Myristic acid, Lactic Acid, D-Lactose, Glycine, L-Isoleucine, Cysteine, L-Valine, L-Threonine, L-Tyrosine, Adenine, Pantothenic acid, L-Tryptophan, Putrescine, L-Aspartic acid, L-Phenylalanine
L-Glutamine
DL-Ornithine, L-Leucine, Niacinamide, Taurine
B
Alanine/Glutamate
Relative abundance ratio (a.u.)
1.0
0.8
0.6
0.4
0.2
0.0
***
*
NG
NT
WG
WT

Figure S3

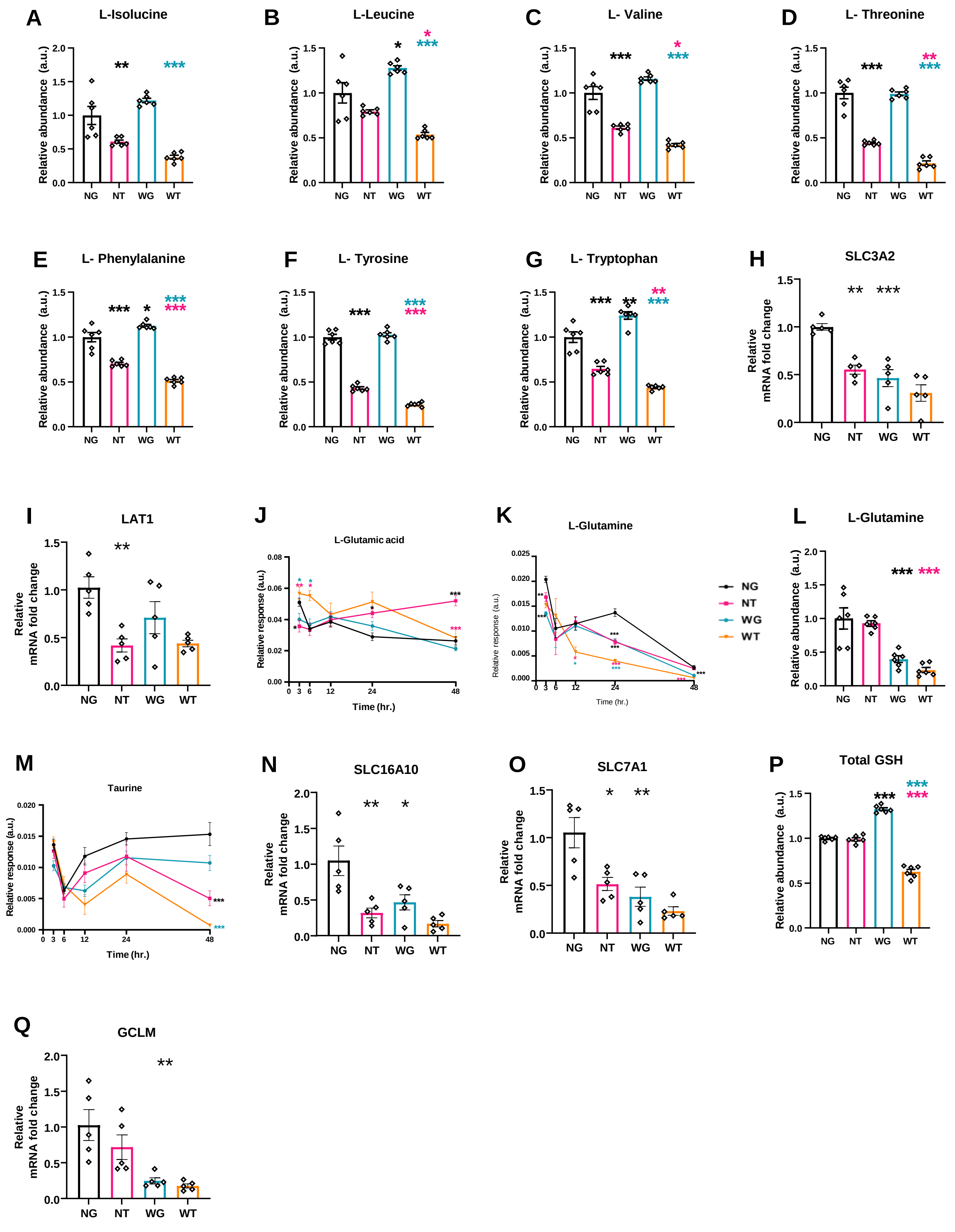
A
L-Isolucine
Relative abundance (a.u.)
B
L-Leucine
C
L- Valine
D
L- Threonine
E
L- Phenylalanine
F
L- Tyrosine
G
L- Tryptophan
H
SLC3A2
Relative mRNA fold change
I
LAT1
J
L-Glutamic acid
Relative response (a.u.)
Time (hr.)
K
L-Glutamine
NG
NT
WG
WT
L
L-Glutamine
M
Taurine
N
SLC16A10
O
SLC7A1
P
Total GSH
Q
GCLM

Figure S4

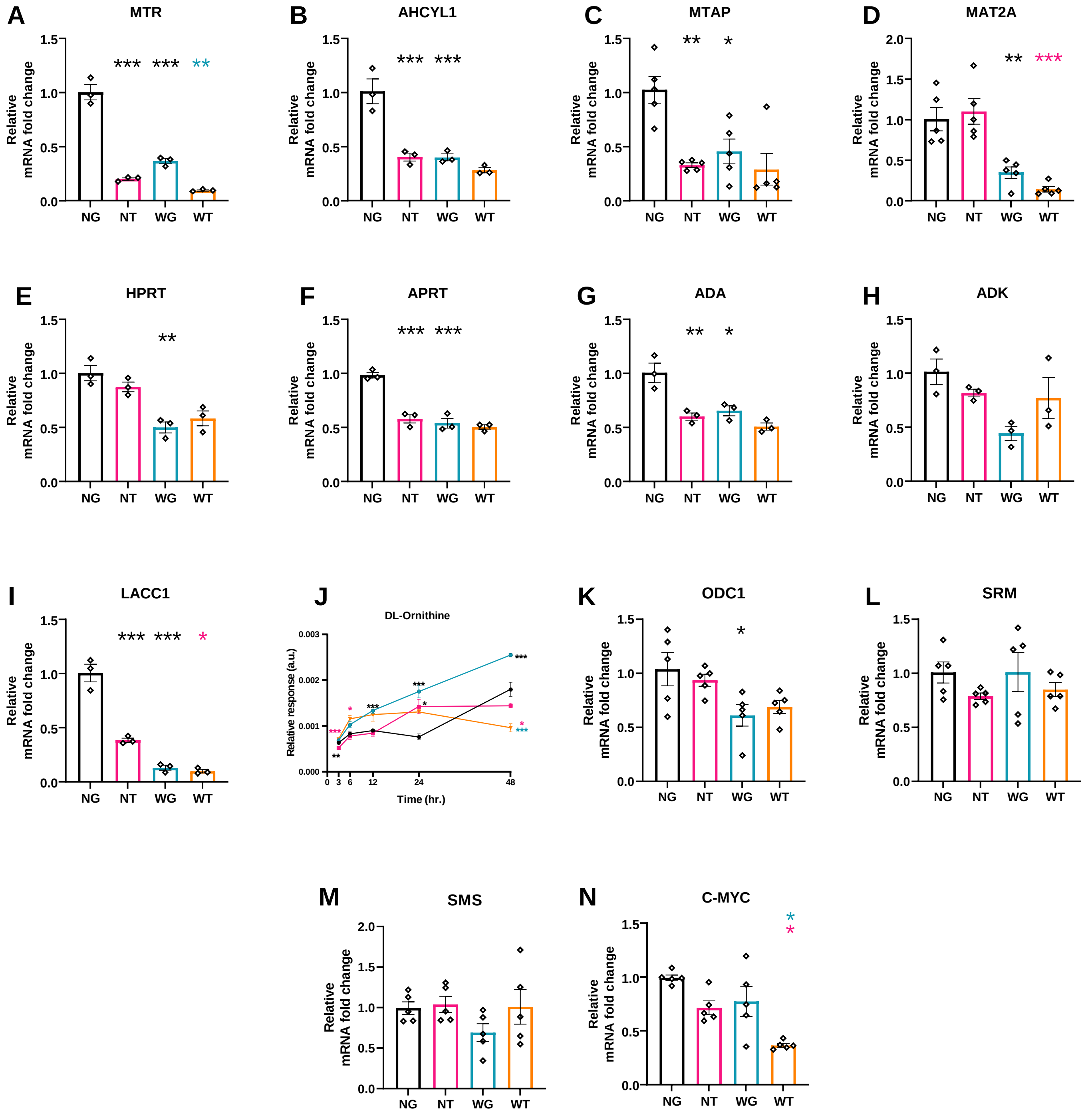

A
MTR
B
AHCYL1
C
MTAP
D
MAT2A
E
HPRT
F
APRT
G
ADA
H
ADK
I
LACC1
J
DL-Ornithine
Relative response (a.u.)
Time (hr.)
K
ODC1
L
SRM
M
SMS
N
C-MYC
Relative
mRNA fold change
NG
NT
WG
WT

Figure S5

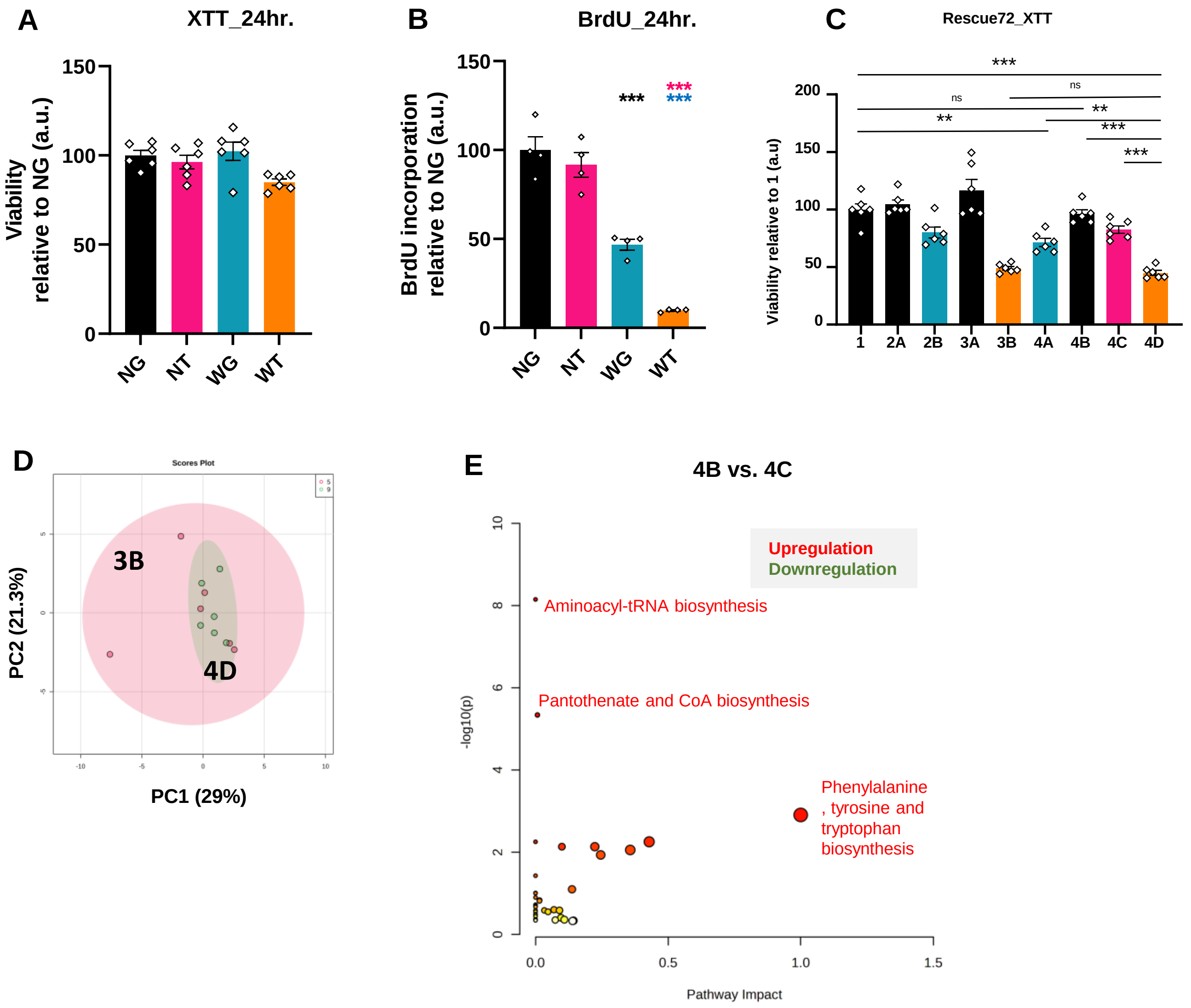

A
XTT_24hr.
Viability relative to NG (a.u.)
150
100
50
0
NG
NT
WG
WT
B
BrdU_24hr.
BrdU incorporation relative to NG (a.u.)
***
***
***
NG
NT
WG
WT
C
Rescue72_XTT
Viability relative to 1 (a.u)
200
150
100
50
0
***
ns
ns
**
**
***
***
1
2A
2B
3A
3B
4A
4B
4C
4D
D
Scores Plot
3B
4D
PC2 (21.3%)
PC1 (29%)
E
4B vs. 4C
Upregulation
Downregulation
Aminoacyl-tRNA biosynthesis
Pantothenate and CoA biosynthesis
Phenylalanine
, tyrosine and
tryptophan
biosynthesis
-log10(p)
Pathway Impact
0.0
0.5
1.0
1.5

**Extended data figure 1**

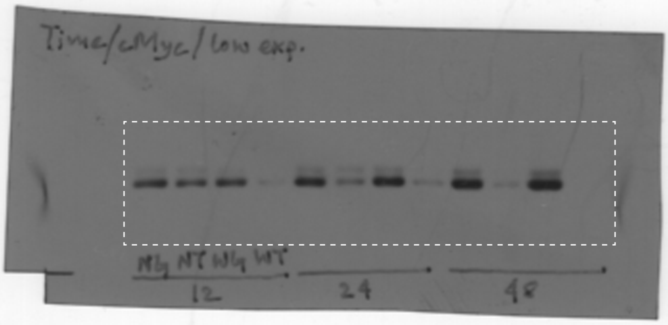


A. Extended data for Figure 5E.
cMyc (low exposure, upper panel)

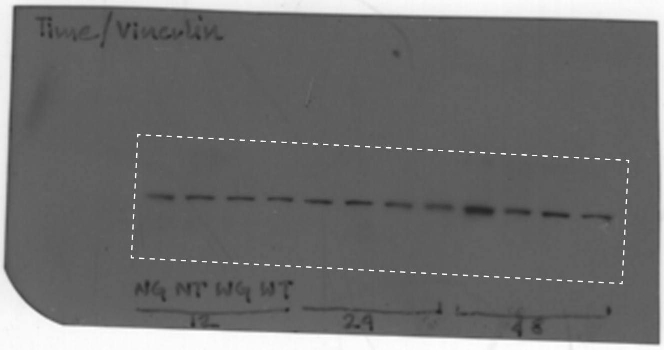


B. Extended data for Figure 5E.
Vinculin (loading control, lower panel)

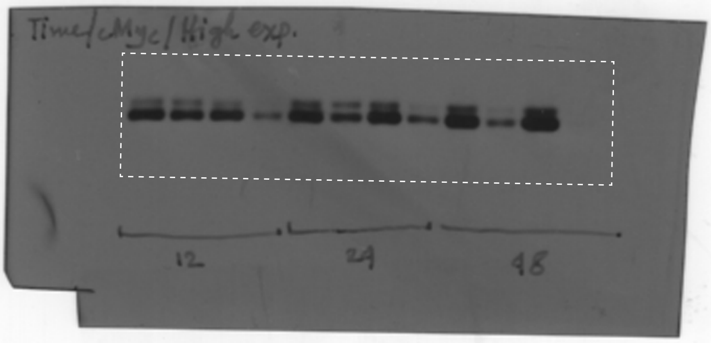


C. Extended data for Figure 5E.
cMyc (high exposure, middle panel)

**Extended data figure 2**

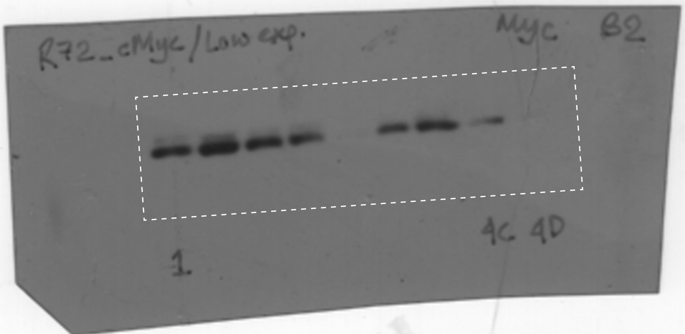


A. Extended data for Figure 6F.
cMyc (low exposure, upper panel)

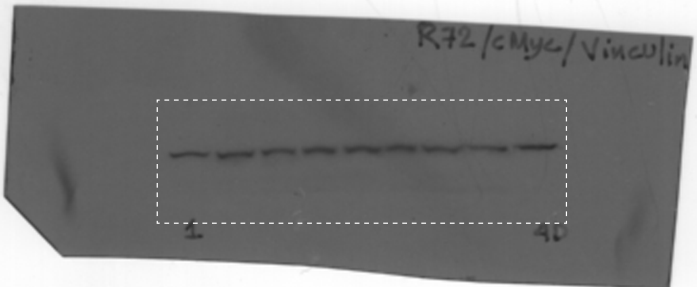


B. Extended data for Figure 6F.
Vinculin (loading control, lower panel)

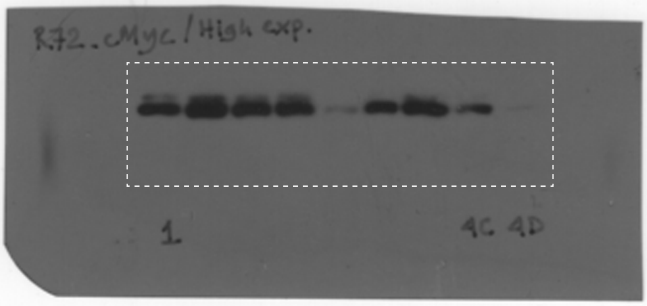


C. Extended data for Figure 6F.
cMyc (high exposure, middle panel)

**Table S1: Authentic standards for metabolite identification**

| Name | Manufacturer | Catalog no./ Identifier |
|---|---|---|
| Adenine | Sigma | A8626 |
| Adenosine | Sigma | A9251 |
| AMP | Sigma | A1752 |
| Citric acid | Sigma | 251275 |
| Cysteine | Sigma | 30089 |
| DL-Ornithine | Sigma | O2375 |
| Glycine | Sigma | G8898 |
| Homovanillic Acid | Fluka | 69673 |
| L-Alanine | Sigma | A7627 |
| L-Aspartic acid | Sigma | A9256 |
| L-Glutamic acid | Sigma | G1251 |
| L-Glutamine | Sigma | G3126 |
| L-Isoleucine | Sigma | 58879 |
| L-Leucine | Sigma | L8000 |
| L-Lysine | Sigma | L5501 |
| L-Phenylalanine | Sigma | P2126 |
| L-Threonine | Sigma | 89179 |
| L-Tyrosine | Sigma | T3754 |
| L-Valine | Sigma | V0500 |
| Malic acid | Supelco | PHR1273 |
| Oleic Acid | Sigma | O1383 |
| Palmitic Acid | Sigma | P0500 |
| Putrescine | Sigma | P5780 |
| Serine | Sigma | S4500 |
| Stearic acid | Sigma | S4751 |
| Succinic acid | Sigma | S7501 |
| Taurine | Sigma | T0625 |
| Tryptophan | Sigma | T0254 |
| D-(-)-Fructose | Sigma | F0127 |
| Hypoxanthine | Sigma | H9377 |
| Pantothenic acid | SRL chemicals | 31269 |

**Table S2: Primers for RT-PCR**

| Gene of Interest | Abv. | Manufacturer | Forward primer (5'-3') | Reverse primer (3'-5') |
|---|---|---|---|---|
| 18S ribosomal RNA | **18S** | Integrated DNA Technologies | TGGTGTTGAGGAAAGCAGAC | ATCTTGTACTGGCGTGGATTC |
| 5-methyltetrahydrofolate-homocysteine methyltransferase | **MTR** | Integrated DNA Technologies | GTAAGCCTAGAGTTCCACCTG | CAAACTTCCTTGATCCTGCAAC |
| Acetyl-CoA carboxylase alpha aka ACACA | **ACC1** | Integrated DNA Technologies | CTGGAGGTGTATGTTCGAAGG | TCTGTTTAGCGTAGGGATGTTC |
| Acetyl-CoA carboxylase beta aka ACACB | **ACC2** | Integrated DNA Technologies | ACCACATCTTCCTCAACTTCG | TGTTGATCTTGACCTCAGCC |
| Adenine phosphoribosyltransferase | **APRT** | Integrated DNA Technologies | TCCTATTCCCTGGAGTACGG | TCACAGGCAGCGTTCATG |
| Adenosine deaminase | **ADA** | Integrated DNA Technologies | CTCATCTTCAAGTCCACCCTG | GTCGAGAAGCTCCCTCTTTTC |
| Adenosine kinase | **ADK** | Integrated DNA Technologies | CAAAGTCGAATATCATGCTGGTG | CTTTTCTCTTCAGGATCTCCCC |
| Adenosylmethionine decarboxylase 1 | **AMD1** | Integrated DNA Technologies | AGTCAGCCAGATCAAACCTTG | GGTATCAGGTCACGAATTCCAC |
| Branched chain amino acid transaminase 1 | **BCAT1** | Integrated DNA Technologies | CCGACGGAACAATGAAGGAT | GGTAGCTGGTGTGACTATTAGG |
| Cellular myelocytomatosis oncogene | **c-Myc** | Integrated DNA Technologies | TTCGGGTAGTGGAAAACCAG | AGTAGAAATACGGCTGCACC |
| Glucose-6-phosphate dehydrogenase | **G6PDH** | Integrated DNA Technologies | AAACGGTCGTACACTTCGG | ATCGCCCTGGAAAAGCTC |
| Glutamate-Cysteine Ligase Catalytic Subunit | **GCLC** | Integrated DNA Technologies | AGACAATGAGATTTAAGCCCC | TGTAGGAAAGGATCACTCTGG |
| Glutamate-Cysteine Ligase Modifier Subunit | **GCLM** | Integrated DNA Technologies | ACTAGAAGTGCAGTTGACATG | GGCTGTAAATGCTCCAAGG |
| Glutamic-oxaloacetic transaminase 1 cytosolic | **GOT1** | Integrated DNA Technologies | CGGATTCTGACCATGAGATCTG | ACCAGATACTCAACCTGCTTG |
| Glutamic-oxaloacetic transaminase 2, mitochondrial | **GOT2** | Integrated DNA Technologies | TTTCTGGAACTGGAGCCTTAAG | ATACCGATAACCTTGTAGCTGC |
| Glutamic-pyruvic transaminase 1 | **GPT 1** | Integrated DNA Technologies | TCGCAGTTCCACTCATTCAAG | GTTCACCACCTCCACATAGC |
| Glutamic-pyruvic transaminase 2 | **GPT2** | Integrated DNA Technologies | CATTCACAGAGGTCATCCGAG | GGCACGTTTCTTAGCATCTTC |
| Glutaminase 1 | **GLS 1** | Integrated DNA Technologies | GAAAGAGTACTGAGCCCTGAAG | GGACAACTAAAAGAATGCCCC |
| Glutaminase 2 | **GLS2** | Integrated DNA Technologies | GCCTGGGTGATTTGCTCTTTT | CCTTTAGTGCAGTGGTGAACTT |
| Hypoxanthine phosphoribosyltransferase 1 | **HPRT1** | Integrated DNA Technologies | TGCTGAGGATTTGGAAAGGG | ACAGAGGGCTACAATGTGATG |
| Laccase Domain Containing 1 | **LACC 1** | Integrated DNA Technologies | ATCGACATCCGTAAAGCCAC | CCAAAATTAAGGCCATCTCGG |
| L-type amino acid transporter 1 (SLC7A5) | **LAT1** | Integrated DNA Technologies | TCCATCCTCTCCATGATCCA | ACGCAGAGCCAGTTGAAG |
| Methionine adenosyltransferase II alpha | **MAT2A** | Integrated DNA Technologies | GCACATTCCTTTTCACCTCAG | CAGTTTCACAAGCTACTTTGGC |
| Methylthioadenosine Phosphorylase | **MTAP** | Integrated DNA Technologies | AGTCATTCTTGTGCCAGAGG | TCCCTCGATTGTGACCATTG |
| Ornithine decarboxylase 1 | **ODC1** | Integrated DNA Technologies | TGATTCCAAAGCAGTCTGTCG | AGGTCTCAGGATCGGTACAG |
| Polyamine Oxidase | **PAOX** | Integrated DNA Technologies | CTTTCCAGTGTCGGTAGAGTG | TCTTCCTGATTGCTTCTGCC |
| S-adenosylhomocysteine hydrolase-like protein 1 | **AHCYL1** | Integrated DNA Technologies | CGGGAGATTGAGATTGCAGAG | AGCCCACTATTTTAGCACCAG |
| Serine hydroxymethyltransferase 1 | **SHMT1** | Integrated DNA Technologies | CCTCCCCATTTGAACACTG | AGAGACTCCAGGTTGTACAG |
| Serine hydroxymethyltransferase 2 | **SHMT2** | Integrated DNA Technologies | AGTGATCCTGAGATGTGGG | GATAACCCTCCGAGTACTTG |
| Solute Carrier Family 1 Member 5 (ASCT2) Glutamine Transporter | **SLC1A5** | Integrated DNA Technologies | CCCCTCATCTACTTCCTCTTC | CATTATTCTCCTCCACGCAC |
| Solute Carrier Family 16 Member 1 (Monocarboxylate transporter 1 | **SLC16A1** | Integrated DNA Technologies | CGTCTGTATTGGAGTCATTGG | AGAGTACAGAGGAACACAGG |
| Solute Carrier Family 16 Member 10 | **SLC16A10** | Integrated DNA Technologies | GCTCCAAAGACGATGACAAG | GCTGTTTTCCGACAACCAA |
| Solute Carrier Family 16 Member 3 | **SLC16A3** | Integrated DNA Technologies | GCATTTCCTGAAGGCTGAG | CTTCTGTACCTCCTCCCTG |
| Solute carrier family 2, facilitated glucose transporter member 1 (GL | **SLC2a1** | Integrated DNA Technologies | GATTGGCTCCTTCTCTGTG | CCAGGATCAGCATCTCAAAG |
| Solute carrier family 2, facilitated glucose transporter member 3(GL | **SLC2a3** | Integrated DNA Technologies | TCATTTCCATTGTGCTCCAG | GATAGTATTAACCACACCCGC |
| Solute carrier family 3 member 2 (CD98) | **SLC3A2** | Integrated DNA Technologies | TGCTCTGGAGTTTTGGCTG | AGTTAGTCCCCGCAATCAAG |
| Solute Carrier Family 43 Member 1 (Large neutral amino acids trans | **SLC43A1** | Integrated DNA Technologies | ACGAGGGCTTCTATTCCAG | AGCACGAAGGAACCAATG |
| Solute Carrier Family 43 Member 2 (Transport of inorganic cations/a | **SLC43A2** | Integrated DNA Technologies | CCTCATTAACACTGCCCAAC | ACGATGAAGGAGACACCAG |
| Solute Carrier Family 7 Member 1 | **SLC7A1** | Integrated DNA Technologies | GCTGAAAACCCCGACATATTC | CCTGACACCATTATGAAGCC |
| Solute Carrier Family 7 Member 11 | **SLC7A11** | Integrated DNA Technologies | GCTGGGCTGATTTATCTTCG | GAATAGAGGGAAAGGGCAAC |

| | | | | |
|---|---|---|---|---|
| Spermidine synthase | **SRM** | Integrated DNA Technologies | GTCCAGTGTGAGATCGACG | ACTCAAAACCGTCACCCAC |
| Spermidine/spermine N1-acetyltransferase 1 (SAT1) | **SSAT1** | Integrated DNA Technologies | CCGAAAGAGCACTGGACTC | TCTGATCCTATGCCAAAGCC |
| Spermine oxidase | **SMOX** | Integrated DNA Technologies | GGCTCCTATTCATACACGCAG | GGTGGAATAGTACTTGCGGTG |
| Spermine synthase | **SMS** | Integrated DNA Technologies | TCTACACTCGAAGCAGTTTGG | CACCTCCCAGAATGAGTACATC |

**Table S3: Metabolite annotation**

| Name | RT (min) | Quantifier m/z | Qualifier m/z | | Identification level** | NIST Match Score |
|---|---|---|---|---|---|---|
| Lactic Acid, 2TMS derivative | 10.305 | 117 | 191 | 147 | 2 | 946 |
| L-Valine, TMS derivative | 10.948 | 72 | 174 | | 1 | 877 |
| L-Alanine, 2TMS derivative | 11.536 | 116 | 190 | | 1 | 820 |
| Glycine, 2TMS derivative | 12.011 | 102 | 204 | 147 | 1 | 923 |
| L-Leucine, TMS derivative | 12.94 | 86 | 188 | 146 | 1 | 864 |
| L-Isoleucine, TMS derivative | 13.526 | 86 | 188 | 146 | 1 | 916 |
| L-Valine, 2TMS derivative | 14.825 | 144 | 218 | | 1 | 719 |
| Isonicotinic Acid, TMS derivative NIST | 16.115 | 180 | 136 | 106 | 2 | 864 |
| L-Leucine, 2TMS derivative | 16.42 | 158 | 232 | 218 | 1 | 908 |
| L-Isoleucine, 2TMS derivative | 17.013 | 158 | 218 | | 1 | 818 |
| Glycine, 3TMS derivative | 17.316 | 174 | 276 | 248 | 1 | 937 |
| Succinic acid 2TMS | 17.472 | 247 | 172 | 147 | 1 | 728 |
| Serine, 3TMS derivative | 18.929 | 204 | 218 | | 1 | 721 |
| L-Threonine, 3TMS derivative | 19.652 | 218 | 291 | | 1 | 929 |
| L-Aspartic acid, 2TMS derivative | 20.362 | 160 | 130 | | 1 | 887 |
| Niacinamide, TMS derivative | 21.558 | 179 | 136 | | 2 | 719 |
| Aminomalonic acid, tris(trimethylsilyl)- | 21.724 | 218 | 320 | | 2 | 824 |
| Malic acid, 3TMS derivative | 22.236 | 233 | 245 | | 1 | 854 |
| L-5-Oxoproline, , 2TMS derivative | 22.848 | 156 | 230 | | 2 | 918 |
| L-Phenylalanine, TMS derivative | 23.253 | 120 | 146 | | 1 | 717 |
| Creatinine - 3TMS Derivative | 23.71 | 115 | 329 | 143 | 2 | 861 |
| Cysteine, 3TMS derivative | 23.767 | 220 | 218 | | 1 | 698 |
| L-Glutamic acid, 3TMS derivative | 25.293 | 246 | 348 | 128 | 1 | 874 |
| L-Phenylalanine, 2TMS derivative | 25.315 | 218 | 192 | | 1 | 846 |
| Taurine, 3TMS derivative | 26.319 | 326 | 188 | 174 | 1 | 888 |
| Putrescine, 4TMS derivative | 27.78 | 174 | 214 | | 1 | 843 |
| Homovanillic Acid, 2TMS derivative | 28.497 | 326 | 311 | 209 | 1 | 901 |
| L-Glutamine, 3TMS derivative | 28.645 | 156 | 245 | | 1 | 759 |
| Hypoxanthine, bis-TMS | 29.177 | 265 | 280 | | 1 | 735 |
| DL-Ornithine, 4TMS derivative | 29.616 | 142 | 200 | 174 | 1 | 739 |
| Citric acid, 4TMS derivative | 29.828 | 273 | 375 | 347 | 1 | 779 |
| Myristic acid, TMS derivative | 29.896 | 285 | 132 | | 2 | 613 |
| Adenine, 2TMS derivative | 30.412 | 264 | 279 | 192 | 1 | 830 |
| Tyrosine, 2TMS derivative | 30.789 | 179 | 310 | 208 | 1 | 779 |
| D-(-)-Fructose, pentakis(trimethylsilyl) ( | 31.119 | 217 | 307 | 103 | 1 | 726 |
| L-Lysine, 4TMS derivative | 31.681 | 174 | 317 | | 1 | 808 |
| L-Tyrosine, 3TMS derivative | 32.005 | 218 | 280 | | 1 | 924 |
| D-Glucitol, 6TMS derivative | 32.385 | 319 | 217 | 205 | 3 | 720 |
| D-(+)-Gluconolactone, 4TMS derivative | 32.676 | 319 | 361 | 129 | 3 | 830 |
| D-Gluconic acid, 6TMS derivative | 32.928 | 292 | 333 | | 3 | 585 |
| Pantothenic acid tritms | 33.115 | 291 | 420 | 103 | 1 | 762 |

| | | | | | | |
|---|---|---|---|---|---|---|
| Palmitic Acid, TMS derivative | 33.723 | 313 | 328 | 117 | 1 | 931 |
| D-Ribofuranose, 1,2,3-tris-O-(trimethyl | 33.805 | 169 | 258 | | 3 | 696 |
| Myo-Inositol, 6TMS derivative | 35.181 | 305 | 318 | 217 | 2 | 885 |
| Linoleic acid, TMS | 36.694 | 337 | 262 | | 2 | 708 |
| Oleic Acid, (Z)-, TMS derivative | 36.784 | 339 | 264 | 117 | 1 | 812 |
| 9-Octadecenoic acid, (E)-, TMS derivati | 36.906 | 339 | 264 | 117 | 2 | 876 |
| Tryptophan, 3TMS derivative | 37.131 | 202 | 291 | | 1 | 856 |
| Stearic acid, TMS derivative | 37.237 | 341 | 132 | 117 | 1 | 931 |
| Ribose phosphate, TMS derivative | 37.384 | 587 | 342 | 299 | 3 | 612 |
| Spermine, 6TMS derivative | 37.766 | 174 | 200 | | 3 | 568 |
| Arachidonic acid, TMS derivative | 39.333 | 117 | 91 | 79 | 2 | 778 |
| Suger_40.209 | 40.209 | 217 | 292 | 204 | 3 | --- |
| Myo-Inositol, 1,3,4,5,6-pentakis-O-(trin | 40.881 | 318 | 387 | 315 | 2 | 910 |
| Steroid_41.886 | 41.885 | 366 | 245 | 169 | 3 | --- |
| Disaccharide_42.057 | 42.057 | 361 | 217 | 204 | 3 | --- |
| Disaccharide_42.366 | 42.366 | 361 | 217 | 204 | 3 | --- |
| 1-Monopalmitin, 2TMS derivative | 42.938 | 371 | 459 | | 2 | 648 |
| Adenosine, 4TMS derivative | 43.893 | 230 | 540 | 245 | 1 | 842 |
| D-Lactose, octakis(trimethylsilyl) ether, | 45.279 | 361 | 319 | 217 | 2 | 935 |
| D-Lactose, octakis(trimethylsilyl) ether, | 45.492 | 361 | 319 | 217 | 2 | 903 |
| Disaccharide_45.760 | 45.76 | 204 | 361 | 217 | 3 | --- |
| D-(+)-Trehalose, octakis(trimethylsilyl) | 45.956 | 361 | 217 | 191 | 3 | 927 |
| Disaccharide_46.713 | 46.71 | 361 | 217 | 204 | 3 | --- |
| AMP | 49.691 | 315 | 230 | 169 | 1 | 823 |
| Cholesterol, TMS derivative | 50.336 | 368 | 353 | 329 | 2 | 925 |
| X_RT | RT | | | | 4 | --- |

** Categorization according to The Chemical Analysis Working Group (CAWG) of the Metabolomics Standards Initiative (MSI) and Core Information for Metabolomics Reporting (CIMR)

**Level 1 (Highest):** "Identified". Matching to Authentic standard

**Level 2 (High):** "Putatively annotated compounds". Annotation verified using additional techniques (NIST library)

**Level 3 (Medium):** "Putatively characterized compound classes". Level 4 plus spectral or chemical properties that indicate class.

**Level 4 (Low):** "Unknown". Are producible and quantifiable MS (or NMR) signal

**Table S4: Enriched pathway in NT compared to NG**

| Pathway name | Total | Expected | Hits | Raw p | FDR |
|---|---|---|---|---|---|
| **Aminoacyl-tRNA biosynthesis** | **48** | **0.83613** | **10** | **1.75E-09** | **1.47E-07** |
| **Glutathione metabolism** | **28** | **0.48774** | **5** | **8.13E-05** | **0.003416** |
| **Pantothenate and CoA biosynthesis** | **19** | **0.33097** | **4** | **0.0002373** | **0.004762** |
| **Valine, leucine and isoleucine biosynthesis** | **8** | **0.13935** | **3** | **0.0002494** | **0.004762** |
| **Biosynthesis of unsaturated fatty acids** | **36** | **0.6271** | **5** | **0.0002835** | **0.004762** |
| **Alanine, aspartate and glutamate metabolism** | **28** | **0.48774** | **4** | **0.0011252** | **0.015753** |
| **Phenylalanine, tyrosine and tryptophan biosynthesis** | **4** | **0.069677** | **2** | **0.0017167** | **0.020601** |
| Phenylalanine metabolism | 10 | 0.17419 | 2 | 0.012069 | 0.12673 |
| Glyoxylate and dicarboxylate metabolism | 32 | 0.55742 | 3 | 0.016704 | 0.15257 |
| Glycine, serine and threonine metabolism | 33 | 0.57484 | 3 | 0.018163 | 0.15257 |
| Arginine biosynthesis | 14 | 0.24387 | 2 | 0.023381 | 0.17854 |
| Butanoate metabolism | 15 | 0.26129 | 2 | 0.026691 | 0.18683 |
| Histidine metabolism | 16 | 0.27871 | 2 | 0.030179 | 0.195 |
| Citrate cycle (TCA cycle) | 20 | 0.34839 | 2 | 0.045787 | 0.27472 |
| Linoleic acid metabolism | 5 | 0.087097 | 1 | 0.08422 | 0.44475 |
| Porphyrin and chlorophyll metabolism | 30 | 0.52258 | 2 | 0.09429 | 0.44475 |
| Nitrogen metabolism | 6 | 0.10452 | 1 | 0.10022 | 0.44475 |
| D-Glutamine and D-glutamate metabolism | 6 | 0.10452 | 1 | 0.10022 | 0.44475 |
| Purine metabolism | 65 | 1.1323 | 3 | 0.1006 | 0.44475 |
| Thiamine metabolism | 7 | 0.12194 | 1 | 0.11596 | 0.48702 |
| Taurine and hypotaurine metabolism | 8 | 0.13935 | 1 | 0.13143 | 0.52571 |
| Arginine and proline metabolism | 38 | 0.66194 | 2 | 0.14011 | 0.53286 |
| Ubiquinone and other terpenoid-quinone biosynthesis | 9 | 0.15677 | 1 | 0.14664 | 0.53286 |
| Valine, leucine and isoleucine degradation | 40 | 0.69677 | 2 | 0.15225 | 0.53286 |
| Primary bile acid biosynthesis | 46 | 0.80129 | 2 | 0.18981 | 0.63776 |
| Nicotinate and nicotinamide metabolism | 15 | 0.26129 | 1 | 0.23264 | 0.75161 |
| Fructose and mannose metabolism | 20 | 0.34839 | 1 | 0.29788 | 0.92673 |
| beta-Alanine metabolism | 21 | 0.36581 | 1 | 0.31027 | 0.93081 |
| Pyruvate metabolism | 22 | 0.38323 | 1 | 0.32245 | 0.93399 |
| Propanoate metabolism | 23 | 0.40065 | 1 | 0.33442 | 0.93638 |
| Glycolysis / Gluconeogenesis | 26 | 0.4529 | 1 | 0.36913 | 0.9983 |
| Galactose metabolism | 27 | 0.47032 | 1 | 0.38031 | 0.9983 |
| Cysteine and methionine metabolism | 33 | 0.57484 | 1 | 0.44347 | 1 |
| Arachidonic acid metabolism | 36 | 0.6271 | 1 | 0.47268 | 1 |
| Amino sugar and nucleotide sugar metabolism | 37 | 0.64452 | 1 | 0.48208 | 1 |
| Fatty acid elongation | 39 | 0.67935 | 1 | 0.50041 | 1 |
| Fatty acid degradation | 39 | 0.67935 | 1 | 0.50041 | 1 |
| Tryptophan metabolism | 41 | 0.71419 | 1 | 0.51811 | 1 |
| Steroid biosynthesis | 42 | 0.73161 | 1 | 0.52673 | 1 |
| Tyrosine metabolism | 42 | 0.73161 | 1 | 0.52673 | 1 |
| Fatty acid biosynthesis | 47 | 0.81871 | 1 | 0.56766 | 1 |
| Steroid hormone biosynthesis | 85 | 1.4806 | 1 | 0.78478 | 1 |

**Table S5: Enriched pathway in WT compared to NG**

| Name | Total | Expected | Hits | Raw p | FDR |
|---|---|---|---|---|---|
| **Aminoacyl-tRNA biosynthesis** | **48** | **0.92903** | **12** | **1.06E-11** | **8.87E-10** |
| **Valine, leucine and isoleucine biosynthesis** | **8** | **0.15484** | **4** | **7.59E-06** | **0.000319** |
| **Pantothenate and CoA biosynthesis** | **19** | **0.36774** | **4** | **0.000362** | **0.009989** |
| **Biosynthesis of unsaturated fatty acids** | **36** | **0.69677** | **5** | **0.000476** | **0.009989** |
| **Phenylalanine, tyrosine and tryptophan biosynthesis** | **4** | **0.077419** | **2** | **0.002122** | **0.03565** |
| Taurine and hypotaurine metabolism | 8 | 0.15484 | 2 | 0.009436 | 0.1321 |
| Phenylalanine metabolism | 10 | 0.19355 | 2 | 0.014803 | 0.14439 |
| Glutathione metabolism | 28 | 0.54194 | 3 | 0.01547 | 0.14439 |
| Alanine, aspartate and glutamate metabolism | 28 | 0.54194 | 3 | 0.01547 | 0.14439 |
| Glyoxylate and dicarboxylate metabolism | 32 | 0.61935 | 3 | 0.022234 | 0.18435 |
| Glycine, serine and threonine metabolism | 33 | 0.63871 | 3 | 0.024141 | 0.18435 |
| Arginine biosynthesis | 14 | 0.27097 | 2 | 0.028532 | 0.19972 |
| Nicotinate and nicotinamide metabolism | 15 | 0.29032 | 2 | 0.032529 | 0.21019 |
| Valine, leucine and isoleucine degradation | 40 | 0.77419 | 3 | 0.039918 | 0.23951 |
| Primary bile acid biosynthesis | 46 | 0.89032 | 3 | 0.056751 | 0.3178 |
| Linoleic acid metabolism | 5 | 0.096774 | 1 | 0.093216 | 0.48938 |
| D-Glutamine and D-glutamate metabolism | 6 | 0.11613 | 1 | 0.11082 | 0.51717 |
| Nitrogen metabolism | 6 | 0.11613 | 1 | 0.11082 | 0.51717 |
| Purine metabolism | 65 | 1.2581 | 3 | 0.12795 | 0.53802 |
| Thiamine metabolism | 7 | 0.13548 | 1 | 0.1281 | 0.53802 |
| Ubiquinone and other terpenoid-quinone biosynthesis | 9 | 0.17419 | 1 | 0.16169 | 0.64674 |
| Biotin metabolism | 10 | 0.19355 | 1 | 0.17801 | 0.67966 |
| Fatty acid biosynthesis | 47 | 0.90968 | 2 | 0.23006 | 0.84023 |
| Histidine metabolism | 16 | 0.30968 | 1 | 0.26966 | 0.94383 |
| Starch and sucrose metabolism | 18 | 0.34839 | 1 | 0.29796 | 1 |
| Fructose and mannose metabolism | 20 | 0.3871 | 1 | 0.3252 | 1 |
| Citrate cycle (TCA cycle) | 20 | 0.3871 | 1 | 0.3252 | 1 |
| beta-Alanine metabolism | 21 | 0.40645 | 1 | 0.33843 | 1 |
| Pyruvate metabolism | 22 | 0.42581 | 1 | 0.35141 | 1 |
| Lysine degradation | 25 | 0.48387 | 1 | 0.38889 | 1 |
| Glycolysis / Gluconeogenesis | 26 | 0.50323 | 1 | 0.40091 | 1 |
| Galactose metabolism | 27 | 0.52258 | 1 | 0.4127 | 1 |
| Porphyrin and chlorophyll metabolism | 30 | 0.58065 | 1 | 0.44675 | 1 |
| Cysteine and methionine metabolism | 33 | 0.63871 | 1 | 0.47889 | 1 |
| Arachidonic acid metabolism | 36 | 0.69677 | 1 | 0.50922 | 1 |
| Amino sugar and nucleotide sugar metabolism | 37 | 0.71613 | 1 | 0.51894 | 1 |
| Arginine and proline metabolism | 38 | 0.73548 | 1 | 0.52848 | 1 |
| Fatty acid elongation | 39 | 0.75484 | 1 | 0.53784 | 1 |
| Fatty acid degradation | 39 | 0.75484 | 1 | 0.53784 | 1 |
| Pyrimidine metabolism | 39 | 0.75484 | 1 | 0.53784 | 1 |
| Tryptophan metabolism | 41 | 0.79355 | 1 | 0.55601 | 1 |
| Steroid biosynthesis | 42 | 0.8129 | 1 | 0.56484 | 1 |
| Tyrosine metabolism | 42 | 0.8129 | 1 | 0.56484 | 1 |
| Steroid hormone biosynthesis | 85 | 1.6452 | 1 | 0.81886 | 1 |

**Table S6: Result of venn diagram analysis**

| Cluster | total | Compounds |
|---|---|---|
| NT WG WT | 15 | X_21.200 |
| | | X_29.345 |
| | | X_16.330 |
| | | Aminomalonic acid |
| | | X_17.099 |
| | | X_32.247 |
| | | X_46.763 |
| | | X_15.939 |
| | | X_15.749 |
| | | Citric acid |
| | | AMP |
| | | Disaccharide_45.760 |
| | | Malic acid |
| | | X_16.323 |
| | | Stearic acid |
| NT WG | 2 | L-5-Oxoproline |
| | | X_29.730 |
| | | X_42.694 |
| | | Suger_40.209 |
| | | Arachidonic acid |
| | | Glycine |
| | | X_32.468 |
| | | X_28.991 |
| | | D-Glucitol |
| | | L-Isoleucine |
| | | X_17.806 |
| | | Cysteine |
| | | X_33.729 |
| | | X_32.878 |
| | | X_14.172 |
| | | L-Valine |
| | | L-Threonine |
| | | Cholesterol |
| | | X_15.459 |
| | | X_25.446 |
| | | X_14.084 |
| | | X_32.310 |
| | | X_20.506 |
| | | X_21.270 |
| | | X_23.515 |
| | | X_28.878 |
| | | X_39.104 |
| | | Suger_24.619 |

Upregulated
Downregulated

| | | |
|---|---|---|
| NT WT | 67 | X_18.279 |
| | | Palmitic Acid |
| | | D-(+)-Gluconolactone |
| | | X_24.278 |
| | | L-Tyrosine |
| | | X_25.320 |
| | | X_44.240 |
| | | X_41.628 |
| | | X_29.565 |
| | | Oleic Acid |
| | | Adenine |
| | | Linoleic acid |
| | | Suger_30.012 |
| | | X_42.126 |
| | | Pantothenic acid |
| | | Steroid_41.886 |
| | | X_26.178 |
| | | Disaccharide_42.057 |
| | | X_27.180 |
| | | Disaccharide_42.366 |
| | | D-(-)-Fructose |
| | | Hypoxanthine |
| | | X_28.815 |
| | | 1-Monopalmitin |
| | | X_28.159 |
| | | Myristic acid |
| | | Suger_30.146 |
| | | X_31.849 |
| | | X_20.866 |
| | | X_26.966 |
| | | Lactic Acid |
| | | D-Lactose |
| | | X_35.540 |
| | | X_40.278 |
| | | L-Tryptophan |
| | | Putrescine |
| | | X_41.489 |
| | | X_20.859 |
| | | L-Aspartic acid |
| | | X_39.849 |
| | | L-Phenylalanine |
| WG WT | 2 | X_40.517 |
| | | L-Glutamine |
| NT | 7 | X_35.092 |
| | | X_21.717 |
| | | Succinic acid |
| | | X_27.925 |

| | | |
|---|---|---|
| | | X_22.569 |
| | | NAc-AA TYPE |
| | | L-Glutamic acid |
| WG | 1 | L-Lysine |
| WT | 15 | X_17.219 |
| | | X_11.819 |
| | | X_11.75 |
| | | X_18.696 |
| | | X_38.530 |
| | | X_33.180 |
| | | D-Ribofuranose phosphate |
| | | Taurine |
| | | X_38.928 |
| | | D-(+)-Trehalose |
| | | X_39.622 |
| | | Niacinamide |
| | | L-Leucine |
| | | Ribose phosphate |
| | | DL-Ornithine |

**Table S7: Enriched pathways affected in both NT and WT**

| Pathway name | Total | Expected | Hits | Raw p | FDR |
|---|---|---|---|---|---|
| **Aminoacyl-tRNA biosynthesis** | **48** | **0.68129** | **9** | **4.44E-09** | **3.73E-07** |
| **Pantothenate and CoA biosynthesis** | **19** | **0.26968** | **4** | **0.000103** | **0.003722** |
| **Valine, leucine and isoleucine biosynthesis** | **8** | **0.11355** | **3** | **0.000133** | **0.003722** |
| **Phenylalanine, tyrosine and tryptophan biosynthesis** | **4** | **0.056774** | **2** | **0.001135** | **0.022383** |
| **Biosynthesis of unsaturated fatty acids** | **36** | **0.51097** | **4** | **0.001332** | **0.022383** |
| Glutathione metabolism | 28 | 0.39742 | 3 | 0.006465 | 0.090503 |
| Phenylalanine metabolism | 10 | 0.14194 | 2 | 0.008081 | 0.096975 |
| Glycine, serine and threonine metabolism | 33 | 0.46839 | 3 | 0.010281 | 0.10795 |
| Linoleic acid metabolism | 5 | 0.070968 | 1 | 0.069068 | 0.64464 |
| Thiamine metabolism | 7 | 0.099355 | 1 | 0.0954 | 0.7593 |
| Taurine and hypotaurine metabolism | 8 | 0.11355 | 1 | 0.1083 | 0.7593 |
| Valine, leucine and isoleucine degradation | 40 | 0.56774 | 2 | 0.10847 | 0.7593 |
| Ubiquinone and other terpenoid-quinone biosynthesis | 9 | 0.12774 | 1 | 0.12102 | 0.78197 |
| Primary bile acid biosynthesis | 46 | 0.6529 | 2 | 0.13685 | 0.79374 |
| Fatty acid biosynthesis | 47 | 0.6671 | 2 | 0.14174 | 0.79374 |
| Arginine biosynthesis | 14 | 0.19871 | 1 | 0.18207 | 0.95589 |
| Nicotinate and nicotinamide metabolism | 15 | 0.2129 | 1 | 0.19379 | 0.95755 |
| Histidine metabolism | 16 | 0.2271 | 1 | 0.20534 | 0.95827 |
| Purine metabolism | 65 | 0.92258 | 2 | 0.23479 | 1 |
| Fructose and mannose metabolism | 20 | 0.28387 | 1 | 0.25 | 1 |
| beta-Alanine metabolism | 21 | 0.29806 | 1 | 0.26079 | 1 |
| Pyruvate metabolism | 22 | 0.31226 | 1 | 0.27142 | 1 |
| Glycolysis / Gluconeogenesis | 26 | 0.36903 | 1 | 0.31252 | 1 |
| Galactose metabolism | 27 | 0.38323 | 1 | 0.32245 | 1 |
| Alanine, aspartate and glutamate metabolism | 28 | 0.39742 | 1 | 0.33224 | 1 |
| Porphyrin and chlorophyll metabolism | 30 | 0.42581 | 1 | 0.35141 | 1 |
| Glyoxylate and dicarboxylate metabolism | 32 | 0.45419 | 1 | 0.37005 | 1 |
| Cysteine and methionine metabolism | 33 | 0.46839 | 1 | 0.37918 | 1 |
| Arachidonic acid metabolism | 36 | 0.51097 | 1 | 0.40582 | 1 |
| Amino sugar and nucleotide sugar metabolism | 37 | 0.52516 | 1 | 0.41445 | 1 |
| Arginine and proline metabolism | 38 | 0.53935 | 1 | 0.42297 | 1 |
| Fatty acid elongation | 39 | 0.55355 | 1 | 0.43136 | 1 |
| Fatty acid degradation | 39 | 0.55355 | 1 | 0.43136 | 1 |
| Tryptophan metabolism | 41 | 0.58194 | 1 | 0.44781 | 1 |
| Steroid biosynthesis | 42 | 0.59613 | 1 | 0.45586 | 1 |
| Tyrosine metabolism | 42 | 0.59613 | 1 | 0.45586 | 1 |
| Steroid hormone biosynthesis | 85 | 1.2065 | 1 | 0.71336 | 1 |

**Table S8: Correlation between BrdU incorporation and metabolites**

| Name | Pearson r | p value |
|---|---|---|
| BrdU | 1.00 | |
| Citric acid | 0.97 | 6.30E-10 |
| AMP | 0.95 | 2.70E-08 |
| Malic acid | 0.87 | 1.20E-05 |
| Pantothenic acid | 0.82 | 9.18E-05 |
| Aminomalonic acid | 0.82 | 1.18E-04 |
| Adenine | 0.73 | 1.43E-03 |
| L-Threonine | 0.71 | 2.07E-03 |
| L-Tyrosine | 0.70 | 2.70E-03 |
| Cystine | 0.68 | 3.85E-03 |
| Taurine | 0.66 | 1.09E-02 |
| Glycine | 0.63 | 8.82E-03 |
| L-Glutamine | 0.56 | 2.40E-02 |
| L-Valine | 0.56 | 2.56E-02 |
| Putrescine | 0.55 | 2.61E-02 |
| L-Phenylalanine | 0.54 | 2.93E-02 |
| L-Tryptophan | 0.52 | 3.96E-02 |
| L-Aspartic acid | 0.52 | 4.07E-02 |
| Isonicotinic Acid | 0.41 | 1.12E-01 |
| DL-Ornithine | 0.39 | 1.40E-01 |
| L-Isoleucine | 0.37 | 1.53E-01 |
| L-Leucine | 0.33 | 2.07E-01 |
| L-Asparagine | 0.30 | 2.53E-01 |
| Adenosine | 0.21 | 4.44E-01 |
| Creatinine | 0.09 | 7.42E-01 |
| Serine | 0.08 | 7.64E-01 |
| L-Lysine | -0.02 | 9.27E-01 |
| 9-Octadecenoic acid(E) | -0.13 | 6.35E-01 |
| L-Glutamic acid | -0.24 | 3.64E-01 |
| L-5-Oxoproline | -0.28 | 2.94E-01 |
| Niacinamide | -0.40 | 1.28E-01 |
| Succinic acid | -0.44 | 9.13E-02 |
| Lactic Acid | -0.49 | 5.40E-02 |
| 1-Monopalmitin | -0.54 | 3.06E-02 |
| D-(-)-Fructose | -0.63 | 8.91E-03 |
| Linoleic acid | -0.64 | 7.32E-03 |
| Palmitic Acid | -0.67 | 4.71E-03 |
| Stearic acid | -0.67 | 4.59E-03 |
| Hypoxanthine | -0.67 | 4.57E-03 |
| Oleic Acid | -0.68 | 4.00E-03 |
| Arachidonic acid | -0.71 | 2.22E-03 |
| D-Lactose | -0.71 | 2.20E-03 |
| Myristic acid | -0.74 | 1.17E-03 |

| |
|---|
| **Positive correlation** |
| **Negative correlation** |

**Table S9: Correlation between BrdU incorporation and gene expression**

| Name | Pearson r | p value |
|---|---|---|
| BrdU | 1.00 | |
| MTR | 0.99 | 3.15E-10 |
| AHCYL1 | 0.97 | 1.30E-07 |
| GOT1 | 0.95 | 2.32E-06 |
| BCAT1 | 0.95 | 2.39E-06 |
| APRT | 0.94 | 4.48E-06 |
| LACC1 | 0.93 | 1.49E-05 |
| SLC43A1 | 0.91 | 3.40E-06 |
| SHMT2 | 0.88 | 6.89E-06 |
| MTAP | 0.88 | 7.24E-06 |
| ADA | 0.88 | 1.75E-04 |
| GLS1 | 0.87 | 2.52E-04 |
| SLC16A1 | 0.86 | 1.77E-05 |
| SLC1A5 | 0.82 | 9.09E-05 |
| ACC2 | 0.79 | 2.42E-03 |
| SLC7A11 | 0.79 | 3.09E-04 |
| GPT2 | 0.78 | 2.82E-03 |
| SLC3A2 | 0.77 | 4.51E-04 |
| C-MYC | 0.76 | 6.53E-04 |
| GLS2 | 0.76 | 4.52E-03 |
| ACC1 | 0.75 | 4.60E-03 |
| LAT1 | 0.75 | 1.22E-03 |
| GCLC | 0.75 | 7.84E-04 |
| SLC7A1 | 0.73 | 1.47E-03 |
| SMOX | 0.71 | 2.11E-03 |
| AMD1 | 0.70 | 2.37E-03 |
| SLC16A10 | 0.68 | 7.90E-03 |
| HPRT | 0.66 | 1.86E-02 |
| GPT1 | 0.66 | 1.97E-02 |
| PAOX | 0.57 | 3.40E-02 |
| SHMT1 | 0.54 | 3.17E-02 |
| GCLM | 0.49 | 5.39E-02 |
| ADK | 0.48 | 1.12E-01 |
| SLC16A3 | 0.45 | 7.70E-02 |
| MAT2A | 0.39 | 1.39E-01 |
| SLC2A3/GLUT3 | 0.38 | 1.42E-01 |
| G6PDH | 0.33 | 3.01E-01 |
| SRM | 0.22 | 4.18E-01 |
| GOT2 | 0.17 | 6.06E-01 |
| ODC1 | 0.12 | 6.52E-01 |
| SLC2A1/GLUT1 | -0.07 | 8.08E-01 |
| SLC43A2 | -0.16 | 5.78E-01 |
| SMS | -0.25 | 3.51E-01 |
| SSAT1 | -0.33 | 2.05E-01 |

| |
|---|
| **Positive correlation** |
| **Negative correlation** |

**Table S10: Correlation coefficient (pearson r) among metabolites**

| | BrdU | Monopalmi | adecenoic a | Adenine | Adenosine | inomalonic | AMP | achidonic a | Citric acid | Creatinine | Cystine | -(-)-Fructos | D-Lactose | DL-Ornithin |
|---|---|---|---|---|---|---|---|---|---|---|---|---|---|---|
| BrdU | 1 | -0.54056 | -0.12867 | 0.72655 | 0.206067 | 0.815526 | 0.947096 | -0.70644 | 0.96934 | 0.089435 | 0.678591 | -0.62997 | -0.70687 | 0.385669 |
| 1-Monopal | -0.54056 | 1 | 0.763631 | -0.59267 | -0.27792 | -0.69358 | -0.67527 | 0.879566 | -0.53084 | -0.43756 | -0.68411 | 0.798098 | 0.663496 | -0.50636 |
| 9-Octadec | -0.12867 | 0.763631 | 1 | -0.37681 | -0.08508 | -0.49022 | -0.38438 | 0.770463 | -0.19126 | -0.38553 | -0.51767 | 0.60621 | 0.489809 | -0.43445 |
| Adenine | 0.72655 | -0.59267 | -0.37681 | 1 | 0.536254 | 0.657186 | 0.67457 | -0.64437 | 0.577708 | 0.280794 | 0.593371 | -0.57011 | -0.68926 | 0.44258 |
| Adenosine | 0.206067 | -0.27792 | -0.08508 | 0.536254 | 1 | -0.01831 | 0.10341 | -0.12022 | 0.054057 | -0.0694 | 0.033715 | -0.14047 | -0.00056 | -0.00342 |
| Aminomalo | 0.815526 | -0.69358 | -0.49022 | 0.657186 | -0.01831 | 1 | 0.924007 | -0.88465 | 0.848002 | 0.474617 | 0.826809 | -0.7984 | -0.90437 | 0.574621 |
| AMP | 0.947096 | -0.67527 | -0.38438 | 0.67457 | 0.10341 | 0.924007 | 1 | -0.83987 | 0.907413 | 0.354438 | 0.847796 | -0.79885 | -0.88058 | 0.522859 |
| Arachidoni | -0.70644 | 0.879566 | 0.770463 | -0.64437 | -0.12022 | -0.88465 | -0.83987 | 1 | -0.67036 | -0.55076 | -0.865 | 0.904938 | 0.881644 | -0.66486 |
| Citric acid | 0.96934 | -0.53084 | -0.19126 | 0.577708 | 0.054057 | 0.848002 | 0.907413 | -0.67036 | 1 | 0.175876 | 0.703635 | -0.59718 | -0.72222 | 0.259142 |
| Creatinine | 0.089435 | -0.43756 | -0.38553 | 0.280794 | -0.0694 | 0.474617 | 0.354438 | -0.55076 | 0.175876 | 1 | 0.679309 | -0.65144 | -0.60084 | 0.887454 |
| Cystine | 0.678591 | -0.68411 | -0.51767 | 0.593371 | 0.033715 | 0.826809 | 0.847796 | -0.865 | 0.703635 | 0.679309 | 1 | -0.89829 | -0.91214 | 0.742712 |
| D-(-)-Fruct | -0.62997 | 0.798098 | 0.60621 | -0.57011 | -0.14047 | -0.7984 | -0.79885 | 0.904938 | -0.59718 | -0.65144 | -0.89829 | 1 | 0.884114 | -0.7637 |
| D-Lactose | -0.70687 | 0.663496 | 0.489809 | -0.68926 | -0.00056 | -0.90437 | -0.88058 | 0.881644 | -0.72222 | -0.60084 | -0.91214 | 0.884114 | 1 | -0.72889 |
| DL-Ornithi | 0.385669 | -0.50636 | -0.43445 | 0.44258 | -0.00342 | 0.574621 | 0.522859 | -0.66486 | 0.259142 | 0.887454 | 0.742712 | -0.7637 | -0.72889 | 1 |
| Glycine | 0.630575 | -0.69345 | -0.54221 | 0.625606 | 0.008214 | 0.866976 | 0.789862 | -0.88142 | 0.601962 | 0.798161 | 0.895137 | -0.88962 | -0.91052 | 0.891108 |
| Hypoxanthi | -0.66933 | 0.801241 | 0.606085 | -0.60695 | -0.19069 | -0.82663 | -0.80275 | 0.91951 | -0.61557 | -0.67921 | -0.8737 | 0.941342 | 0.863633 | -0.80907 |
| Isonicotinic | 0.412857 | -0.37748 | -0.25895 | 0.262904 | 0.226316 | 0.402647 | 0.433019 | -0.42062 | 0.407537 | -0.28119 | 0.232343 | -0.36684 | -0.38425 | -0.11513 |
| L-5-Oxopro | -0.27998 | 0.321306 | 0.053418 | 0.101398 | -0.03844 | -0.34059 | -0.37081 | 0.369046 | -0.2961 | -0.41538 | -0.4225 | 0.513681 | 0.313801 | -0.45539 |
| Lactic Acid | -0.49003 | 0.725742 | 0.531695 | -0.42922 | -0.23939 | -0.64113 | -0.65344 | 0.788588 | -0.47934 | -0.55572 | -0.74577 | 0.892613 | 0.691987 | -0.69823 |
| L-Asparagi | 0.303635 | -0.26067 | -0.29502 | 0.205797 | -0.31435 | 0.405048 | 0.42038 | -0.39401 | 0.294444 | 0.202054 | 0.341108 | -0.36306 | -0.47757 | 0.403441 |
| L-Aspartic | 0.516184 | -0.65715 | -0.46052 | 0.450243 | -0.00823 | 0.807264 | 0.718959 | -0.80227 | 0.553894 | 0.782014 | 0.789277 | -0.84394 | -0.81678 | 0.848764 |
| L-Glutamic | -0.24313 | 0.269722 | -0.09497 | 0.008746 | -0.44788 | -0.12122 | -0.23158 | 0.178219 | -0.18108 | -0.18644 | -0.19682 | 0.348808 | 0.088447 | -0.25051 |
| L-Glutamin | 0.560292 | -0.35017 | -0.35806 | 0.458612 | -0.13052 | 0.518058 | 0.582695 | -0.46823 | 0.485823 | -0.16076 | 0.41302 | -0.32212 | -0.54103 | 0.012427 |
| Linoleic aci | -0.64208 | 0.848709 | 0.701845 | -0.56 | -0.2182 | -0.82195 | -0.77546 | 0.92919 | -0.65835 | -0.44112 | -0.72321 | 0.789036 | 0.754431 | -0.548 |
| L-Isoleucin | 0.374532 | -0.643 | -0.47505 | 0.394782 | -0.05683 | 0.712319 | 0.654595 | -0.76979 | 0.516769 | 0.890003 | 0.892464 | -0.83937 | -0.80412 | 0.848418 |
| L-Leucine | 0.333184 | -0.56928 | -0.4218 | 0.40416 | -0.07788 | 0.639811 | 0.577729 | -0.69793 | 0.407412 | 0.949795 | 0.833739 | -0.80081 | -0.76575 | 0.918587 |
| L-Lysine | -0.02482 | -0.26596 | -0.39903 | 0.264249 | -0.08588 | 0.269599 | 0.167334 | -0.42533 | -0.07858 | 0.83452 | 0.555015 | -0.53295 | -0.4935 | 0.853438 |
| L-Phenylal | 0.544257 | -0.64067 | -0.43953 | 0.540086 | 0.060303 | 0.73967 | 0.727545 | -0.79868 | 0.532665 | 0.85799 | 0.904114 | -0.89559 | -0.86497 | 0.929504 |
| L-Threonin | 0.709722 | -0.69595 | -0.454 | 0.642086 | 0.10158 | 0.848775 | 0.842938 | -0.86116 | 0.672119 | 0.736477 | 0.906601 | -0.90482 | -0.90961 | 0.87201 |
| L-Tryptoph | 0.518648 | -0.63685 | -0.44152 | 0.493458 | 0.123039 | 0.646984 | 0.678187 | -0.76209 | 0.437848 | 0.826991 | 0.868823 | -0.88784 | -0.80569 | 0.935293 |
| L-Tyrosine | 0.696865 | -0.70537 | -0.44253 | 0.595208 | 0.156614 | 0.808558 | 0.827404 | -0.85498 | 0.646755 | 0.720514 | 0.911619 | -0.92893 | -0.89685 | 0.849792 |
| L-Valine | 0.555165 | -0.62471 | -0.40359 | 0.521005 | 0.060504 | 0.747773 | 0.722756 | -0.78241 | 0.539792 | 0.868326 | 0.884368 | -0.8735 | -0.8474 | 0.925901 |
| Malic acid | 0.86979 | -0.6833 | -0.3891 | 0.665645 | 0.09965 | 0.939251 | 0.942913 | -0.84882 | 0.872909 | 0.51618 | 0.858499 | -0.82155 | -0.87678 | 0.649838 |
| Myristic aci | -0.73538 | 0.697101 | 0.598936 | -0.69239 | -0.08951 | -0.79919 | -0.78234 | 0.864442 | -0.60255 | -0.51185 | -0.85296 | 0.807609 | 0.852197 | -0.69417 |
| Niacinamid | -0.39713 | 0.526026 | 0.596448 | -0.29409 | 0.371511 | -0.70066 | -0.59144 | 0.736803 | -0.42377 | -0.60761 | -0.72949 | 0.734776 | 0.734056 | -0.72297 |
| Oleic Acid | -0.67655 | 0.90175 | 0.864265 | -0.62271 | -0.21254 | -0.78387 | -0.74672 | 0.950612 | -0.58723 | -0.37184 | -0.74118 | 0.804771 | 0.74319 | -0.49865 |
| Palmitic Ac | -0.66773 | 0.825842 | 0.728589 | -0.75501 | -0.14022 | -0.8399 | -0.79793 | 0.938781 | -0.59479 | -0.59872 | -0.87026 | 0.885724 | 0.907227 | -0.73663 |
| Pantotheni | 0.822507 | -0.71151 | -0.43884 | 0.635971 | 0.20865 | 0.853614 | 0.896553 | -0.87346 | 0.715036 | 0.558114 | 0.884365 | -0.92156 | -0.89815 | 0.74354 |
| Putrescine | 0.553652 | -0.59131 | -0.4514 | 0.430115 | 0.315903 | 0.575682 | 0.563013 | -0.69371 | 0.326749 | 0.657294 | 0.685129 | -0.77117 | -0.60227 | 0.798619 |
| Serine | 0.081538 | -0.333 | -0.42137 | 0.219094 | -0.20619 | 0.400167 | 0.28228 | -0.5081 | 0.072158 | 0.83922 | 0.585237 | -0.58775 | -0.55768 | 0.888656 |
| Stearic aci | -0.66915 | 0.820459 | 0.704575 | -0.7624 | -0.14121 | -0.84168 | -0.80299 | 0.929636 | -0.60422 | -0.59876 | -0.8666 | 0.881726 | 0.911042 | -0.73585 |
| Succinic ac | -0.43613 | 0.330006 | 0.091883 | 0.047291 | -0.09005 | -0.31798 | -0.40087 | 0.326464 | -0.36791 | -0.03694 | -0.29315 | 0.317986 | 0.213005 | -0.06081 |
| Taurine | 0.655606 | -0.76668 | -0.47464 | 0.48661 | -0.0092 | 0.898894 | 0.849144 | -0.83155 | 0.817172 | 0.495956 | 0.821985 | -0.77445 | -0.84999 | 0.563806 |

| | Glycine | Hypoxanthin | onicotinic A | -5-Oxoprolir | Lactic Acid | -Asparagin | -Aspartic ac | Glutamic ac | L-Glutamine | Linoleic aci | L-Isoleucine | L-Leucine | L-Lysine | Phenylalani | L-Threonine |
|---|---|---|---|---|---|---|---|---|---|---|---|---|---|---|---|
| BrdU | 0.630575 | -0.66933 | 0.412857 | -0.27998 | -0.49003 | 0.303635 | 0.516184 | -0.24313 | 0.560292 | -0.64208 | 0.374532 | 0.333184 | -0.02482 | 0.544257 | 0.709722 |
| 1-Monopal | -0.69345 | 0.801241 | -0.37748 | 0.321306 | 0.725742 | -0.26067 | -0.65715 | 0.269722 | -0.35017 | 0.848709 | -0.643 | -0.56928 | -0.26596 | -0.64067 | -0.69595 |
| 9-Octadec | -0.54221 | 0.606085 | -0.25895 | 0.053418 | 0.531695 | -0.29502 | -0.46052 | -0.09497 | -0.35806 | 0.701845 | -0.47505 | -0.4218 | -0.39903 | -0.43953 | -0.454 |
| Adenine | 0.625606 | -0.60695 | 0.262904 | 0.101398 | -0.42922 | 0.205797 | 0.450243 | 0.008746 | 0.458612 | -0.56 | 0.394782 | 0.40416 | 0.264249 | 0.540086 | 0.642086 |
| Adenosine | 0.008214 | -0.19069 | 0.226316 | -0.03844 | -0.23939 | -0.31435 | -0.00823 | -0.44788 | -0.13052 | -0.2182 | -0.05683 | -0.07788 | -0.08588 | 0.060303 | 0.10158 |
| Aminomalo | 0.866976 | -0.82663 | 0.402647 | -0.34059 | -0.64113 | 0.405048 | 0.807264 | -0.12122 | 0.518058 | -0.82195 | 0.712319 | 0.639811 | 0.269599 | 0.73967 | 0.848775 |
| AMP | 0.789862 | -0.80275 | 0.433019 | -0.37081 | -0.65344 | 0.42038 | 0.718959 | -0.23158 | 0.582695 | -0.77546 | 0.654595 | 0.577729 | 0.167334 | 0.727545 | 0.842938 |
| Arachidoni | -0.88142 | 0.91951 | -0.42062 | 0.369046 | 0.788588 | -0.39401 | -0.80227 | 0.178219 | -0.46823 | 0.92919 | -0.76979 | -0.69793 | -0.42533 | -0.79868 | -0.86116 |
| Citric acid | 0.601962 | -0.61557 | 0.407537 | -0.2961 | -0.47934 | 0.294444 | 0.553894 | -0.18108 | 0.485823 | -0.65835 | 0.516769 | 0.407412 | -0.07858 | 0.532665 | 0.672119 |
| Creatinine | 0.798161 | -0.67921 | -0.28119 | -0.41538 | -0.55572 | 0.202054 | 0.782014 | -0.18644 | -0.16076 | -0.44112 | 0.890003 | 0.949795 | 0.83452 | 0.85799 | 0.736477 |
| Cystine | 0.895137 | -0.8737 | 0.232343 | -0.4225 | -0.74577 | 0.341108 | 0.789277 | -0.19682 | 0.41302 | -0.72321 | 0.892464 | 0.833739 | 0.555015 | 0.904114 | 0.906601 |
| D-(-)-Fruct | -0.88962 | 0.941342 | -0.36684 | 0.513681 | 0.892613 | -0.36306 | -0.84394 | 0.348808 | -0.32212 | 0.789036 | -0.83937 | -0.80081 | -0.53295 | -0.89559 | -0.90482 |
| D-Lactose | -0.91052 | 0.863633 | -0.38425 | 0.313801 | 0.691987 | -0.47757 | -0.81678 | 0.088447 | -0.54103 | 0.754431 | -0.80412 | -0.76575 | -0.4935 | -0.86497 | -0.90961 |
| DL-Ornithi | 0.891108 | -0.80907 | -0.11513 | -0.45539 | -0.69823 | 0.403441 | 0.848764 | -0.25051 | 0.012427 | -0.548 | 0.848418 | 0.918587 | 0.853438 | 0.929504 | 0.87201 |
| Glycine | 1 | -0.92764 | 0.123785 | -0.42528 | -0.75749 | 0.39226 | 0.916936 | -0.19316 | 0.271193 | -0.76845 | 0.889775 | 0.897742 | 0.674085 | 0.950418 | 0.965439 |
| Hypoxanthi | -0.92764 | 1 | -0.32053 | 0.557781 | 0.869928 | -0.40008 | -0.90018 | 0.392641 | -0.2578 | 0.873157 | -0.85037 | -0.80849 | -0.544 | -0.91357 | -0.94853 |
| Isonicotinic | 0.123785 | -0.32053 | 1 | -0.27503 | -0.34964 | 0.092601 | 0.144467 | -0.28551 | 0.512504 | -0.52445 | 0.019347 | -0.14676 | -0.28677 | 0.070912 | 0.17994 |
| L-5-Oxopro | -0.42528 | 0.557781 | -0.27503 | 1 | 0.617571 | -0.02838 | -0.6072 | 0.856411 | 0.283122 | 0.50066 | -0.54703 | -0.45538 | -0.18328 | -0.55176 | -0.53131 |
| Lactic Acid | -0.75749 | 0.869928 | -0.34964 | 0.617571 | 1 | -0.35036 | -0.81011 | 0.552513 | -0.04325 | 0.749024 | -0.74585 | -0.69222 | -0.47829 | -0.78497 | -0.79688 |
| L-Asparagi | 0.39226 | -0.40008 | 0.092601 | -0.02838 | -0.35036 | 1 | 0.499044 | 0.164942 | 0.366137 | -0.30135 | 0.342919 | 0.342913 | 0.1666 | 0.364747 | 0.431666 |
| L-Aspartic | 0.916936 | -0.90018 | 0.144467 | -0.6072 | -0.81011 | 0.499044 | 1 | -0.39003 | 0.073641 | -0.76148 | 0.891742 | 0.866533 | 0.546321 | 0.905446 | 0.924499 |
| L-Glutamic | -0.19316 | 0.392641 | -0.28551 | 0.856411 | 0.552513 | 0.164942 | -0.39003 | 1 | 0.406344 | 0.364547 | -0.29167 | -0.21257 | 0.011667 | -0.34175 | -0.34124 |
| L-Glutamin | 0.271193 | -0.2578 | 0.512504 | 0.283122 | -0.04325 | 0.366137 | 0.073641 | 0.406344 | 1 | -0.33979 | 0.088733 | 0.022798 | -0.0841 | 0.140549 | 0.250629 |
| Linoleic aci | -0.76845 | 0.873157 | -0.52445 | 0.50066 | 0.749024 | -0.30135 | -0.76148 | 0.364547 | -0.33979 | 1 | -0.67428 | -0.56514 | -0.25424 | -0.69516 | -0.78205 |
| L-Isoleucin | 0.889775 | -0.85037 | 0.019347 | -0.54703 | -0.74585 | 0.342919 | 0.891742 | -0.29167 | 0.088733 | -0.67428 | 1 | 0.966956 | 0.673466 | 0.943324 | 0.887886 |
| L-Leucine | 0.897742 | -0.80849 | -0.14676 | -0.45538 | -0.69222 | 0.342913 | 0.866533 | -0.21257 | 0.022798 | -0.56514 | 0.966956 | 1 | 0.782763 | 0.951876 | 0.874978 |
| L-Lysine | 0.674085 | -0.544 | -0.28677 | -0.18328 | -0.47829 | 0.1666 | 0.546321 | 0.011667 | -0.0841 | -0.25424 | 0.673466 | 0.782763 | 1 | 0.714278 | 0.57764 |
| L-Phenylal | 0.950418 | -0.91357 | 0.070912 | -0.55176 | -0.78497 | 0.364747 | 0.905446 | -0.34175 | 0.140549 | -0.69516 | 0.943324 | 0.951876 | 0.714278 | 1 | 0.968001 |
| L-Threonin | 0.965439 | -0.94853 | 0.17994 | -0.53131 | -0.79688 | 0.431666 | 0.924499 | -0.34124 | 0.250629 | -0.78205 | 0.887886 | 0.874978 | 0.57764 | 0.968001 | 1 |
| L-Tryptoph | 0.898813 | -0.90044 | 0.074989 | -0.58172 | -0.81663 | 0.362808 | 0.862888 | -0.41683 | 0.113738 | -0.6635 | 0.910091 | 0.921847 | 0.731926 | 0.978629 | 0.933158 |
| L-Tyrosine | 0.930857 | -0.95405 | 0.288455 | -0.62168 | -0.84165 | 0.357375 | 0.898362 | -0.45165 | 0.235383 | -0.79777 | 0.891192 | 0.857028 | 0.573642 | 0.966401 | 0.978465 |
| L-Valine | 0.951301 | -0.90978 | 0.05377 | -0.57976 | -0.77502 | 0.332996 | 0.918725 | -0.37388 | 0.099347 | -0.69903 | 0.945695 | 0.952992 | 0.70349 | 0.994101 | 0.964962 |
| Malic acid | 0.873397 | -0.87186 | 0.300352 | -0.47911 | -0.73166 | 0.452434 | 0.863232 | -0.31572 | 0.352178 | -0.80627 | 0.767179 | 0.701072 | 0.278584 | 0.820419 | 0.92422 |
| Myristic aci | -0.85347 | 0.818373 | -0.37056 | 0.297881 | 0.675525 | -0.2333 | -0.65614 | 0.116014 | -0.5026 | 0.770712 | -0.67485 | -0.64773 | -0.56222 | -0.78014 | -0.82211 |
| Niacinamid | -0.79643 | 0.755004 | -0.14895 | 0.383601 | 0.55463 | -0.47811 | -0.71096 | -0.02931 | -0.35844 | 0.610554 | -0.70541 | -0.703 | -0.60247 | -0.74003 | -0.74567 |
| Oleic Acid | -0.73202 | 0.816005 | -0.49822 | 0.259046 | 0.731039 | -0.30564 | -0.64302 | 0.142951 | -0.49965 | 0.915614 | -0.60231 | -0.51062 | -0.30779 | -0.62343 | -0.70308 |
| Palmitic Ac | -0.8984 | 0.890665 | -0.33285 | 0.201668 | 0.715188 | -0.40213 | -0.74534 | 0.043964 | -0.5446 | 0.810743 | -0.75685 | -0.73574 | -0.5639 | -0.82221 | -0.86231 |
| Pantotheni | 0.886917 | -0.93758 | 0.447875 | -0.606 | -0.82866 | 0.337767 | 0.841158 | -0.46358 | 0.358923 | -0.832 | 0.778105 | 0.722659 | 0.423389 | 0.886564 | 0.942858 |
| Putrescine | 0.782477 | -0.84039 | 0.193495 | -0.70005 | -0.79766 | 0.090923 | 0.767815 | -0.6152 | -0.06159 | -0.70525 | 0.69739 | 0.686571 | 0.576353 | 0.805708 | 0.799754 |
| Serine | 0.734994 | -0.63455 | -0.29685 | -0.30582 | -0.60261 | 0.502362 | 0.754494 | -0.04659 | -0.14373 | -0.37335 | 0.767977 | 0.840605 | 0.853874 | 0.76416 | 0.684592 |
| Stearic aci | -0.89766 | 0.887753 | -0.32537 | 0.192647 | 0.702814 | -0.41332 | -0.7459 | 0.040704 | -0.5513 | 0.797708 | -0.75676 | -0.73875 | -0.55431 | -0.82358 | -0.86518 |
| Succinic ac | -0.19741 | 0.30557 | -0.45459 | 0.70049 | 0.278454 | 0.28291 | -0.25555 | 0.608996 | -0.08645 | 0.471932 | -0.2307 | -0.09864 | 0.195924 | -0.24315 | -0.28406 |
| Taurine | 0.823796 | -0.80453 | 0.252185 | -0.41368 | -0.65924 | 0.434268 | 0.798509 | -0.27935 | 0.342869 | -0.79406 | 0.786309 | 0.699121 | 0.186494 | 0.759918 | 0.844636 |

| | -Tryptopha | L-Tyrosine | L-Valine | Malic acid | Myristic aci | Niacinamide | Oleic Acid | Palmitic Aci | ntothenic a | Putrescine | Serine | Stearic aci | Succinic aci | Taurine |
|---|---|---|---|---|---|---|---|---|---|---|---|---|---|---|
| BrdU | 0.518648 | 0.696865 | 0.555165 | 0.86979 | -0.73538 | -0.39713 | -0.67655 | -0.66773 | 0.822507 | 0.553652 | 0.081538 | -0.66915 | -0.43613 | 0.655606 |
| 1-Monopal | -0.63685 | -0.70537 | -0.62471 | -0.6833 | 0.697101 | 0.526026 | 0.90175 | 0.825842 | -0.71151 | -0.59131 | -0.333 | 0.820459 | 0.330006 | -0.76668 |
| 9-Octadec | -0.44152 | -0.44253 | -0.40359 | -0.3891 | 0.598936 | 0.596448 | 0.864265 | 0.728589 | -0.43884 | -0.4514 | -0.42137 | 0.704575 | 0.091883 | -0.47464 |
| Adenine | 0.493458 | 0.595208 | 0.521005 | 0.665645 | -0.69239 | -0.29409 | -0.62271 | -0.75501 | 0.635971 | 0.430115 | 0.219094 | -0.7624 | 0.047291 | 0.48661 |
| Adenosine | 0.123039 | 0.156614 | 0.060504 | 0.09965 | -0.08951 | 0.371511 | -0.21254 | -0.14022 | 0.20865 | 0.315903 | -0.20619 | -0.14121 | -0.09005 | -0.0092 |
| Aminomalo | 0.646984 | 0.808558 | 0.747773 | 0.939251 | -0.79919 | -0.70066 | -0.78387 | -0.8399 | 0.853614 | 0.575682 | 0.400167 | -0.84168 | -0.31798 | 0.898894 |
| AMP | 0.678187 | 0.827404 | 0.722756 | 0.942913 | -0.78234 | -0.59144 | -0.74672 | -0.79793 | 0.896553 | 0.563013 | 0.28228 | -0.80299 | -0.40087 | 0.849144 |
| Arachidoni | -0.76209 | -0.85498 | -0.78241 | -0.84882 | 0.864442 | 0.736803 | 0.950612 | 0.938781 | -0.87346 | -0.69371 | -0.5081 | 0.929636 | 0.326464 | -0.83155 |
| Citric acid | 0.437848 | 0.646755 | 0.539792 | 0.872909 | -0.60255 | -0.42377 | -0.58723 | -0.59479 | 0.715036 | 0.326749 | 0.072158 | -0.60422 | -0.36791 | 0.817172 |
| Creatinine | 0.826991 | 0.720514 | 0.868326 | 0.51618 | -0.51185 | -0.60761 | -0.37184 | -0.59872 | 0.558114 | 0.657294 | 0.83922 | -0.59876 | -0.03694 | 0.495956 |
| Cystine | 0.868823 | 0.911619 | 0.884368 | 0.858499 | -0.85296 | -0.72949 | -0.74118 | -0.87026 | 0.884365 | 0.685129 | 0.585237 | -0.8666 | -0.29315 | 0.821985 |
| D-(-)-Fruct | -0.88784 | -0.92893 | -0.8735 | -0.82155 | 0.807609 | 0.734776 | 0.804771 | 0.885724 | -0.92156 | -0.77117 | -0.58775 | 0.881726 | 0.317986 | -0.77445 |
| D-Lactose | -0.80569 | -0.89685 | -0.8474 | -0.87678 | 0.852197 | 0.734056 | 0.74319 | 0.907227 | -0.89815 | -0.60227 | -0.55768 | 0.911042 | 0.213005 | -0.84999 |
| DL-Ornithi | 0.935293 | 0.849792 | 0.925901 | 0.649838 | -0.69417 | -0.72297 | -0.49865 | -0.73663 | 0.74354 | 0.798619 | 0.888656 | -0.73585 | -0.06081 | 0.563806 |
| Glycine | 0.898813 | 0.930857 | 0.951301 | 0.873397 | -0.85347 | -0.79643 | -0.73202 | -0.8984 | 0.886917 | 0.782477 | 0.734994 | -0.89766 | -0.19741 | 0.823796 |
| Hypoxanthi | -0.90044 | -0.95405 | -0.90978 | -0.87186 | 0.818373 | 0.755004 | 0.816005 | 0.890665 | -0.93758 | -0.84039 | -0.63455 | 0.887753 | 0.30557 | -0.80453 |
| Isonicotinic | 0.074989 | 0.288455 | 0.05377 | 0.300352 | -0.37056 | -0.14895 | -0.49822 | -0.33285 | 0.447875 | 0.193495 | -0.29685 | -0.32537 | -0.45459 | 0.252185 |
| L-5-Oxopro | -0.58172 | -0.62168 | -0.57976 | -0.47911 | 0.297881 | 0.383601 | 0.259046 | 0.201668 | -0.606 | -0.70005 | -0.30582 | 0.192647 | 0.70049 | -0.41368 |
| Lactic Acid | -0.81663 | -0.84165 | -0.77502 | -0.73166 | 0.675525 | 0.55463 | 0.731039 | 0.715188 | -0.82866 | -0.79766 | -0.60261 | 0.702814 | 0.278454 | -0.65924 |
| L-Asparagi | 0.362808 | 0.357375 | 0.332996 | 0.452434 | -0.2333 | -0.47811 | -0.30564 | -0.40213 | 0.337767 | 0.090923 | 0.502362 | -0.41332 | 0.28291 | 0.434268 |
| L-Aspartic | 0.862888 | 0.898362 | 0.918725 | 0.863232 | -0.65614 | -0.71096 | -0.64302 | -0.74534 | 0.841158 | 0.767815 | 0.754494 | -0.7459 | -0.25555 | 0.798509 |
| L-Glutamic | -0.41683 | -0.45165 | -0.37388 | -0.31572 | 0.116014 | -0.02931 | 0.142951 | 0.043964 | -0.46358 | -0.6152 | -0.04659 | 0.040704 | 0.608996 | -0.27935 |
| L-Glutamin | 0.113738 | 0.235383 | 0.099347 | 0.352178 | -0.5026 | -0.35844 | -0.49965 | -0.5446 | 0.358923 | -0.06159 | -0.14373 | -0.5513 | -0.08645 | 0.342869 |
| Linoleic aci | -0.6635 | -0.79777 | -0.69903 | -0.80627 | 0.770712 | 0.610554 | 0.915614 | 0.810743 | -0.832 | -0.70525 | -0.37335 | 0.797708 | 0.471932 | -0.79406 |
| L-Isoleucin | 0.910091 | 0.891192 | 0.945695 | 0.767179 | -0.67485 | -0.70541 | -0.60231 | -0.75685 | 0.778105 | 0.69739 | 0.767977 | -0.75676 | -0.2307 | 0.786309 |
| L-Leucine | 0.921847 | 0.857028 | 0.952992 | 0.701072 | -0.64773 | -0.703 | -0.51062 | -0.73574 | 0.722659 | 0.686571 | 0.840605 | -0.73875 | -0.09864 | 0.699121 |
| L-Lysine | 0.731926 | 0.573642 | 0.70349 | 0.278584 | -0.56222 | -0.60247 | -0.30779 | -0.5639 | 0.423389 | 0.576353 | 0.853874 | -0.55431 | 0.195924 | 0.186494 |
| L-Phenylal | 0.978629 | 0.966401 | 0.994101 | 0.820419 | -0.78014 | -0.74003 | -0.62343 | -0.82221 | 0.886564 | 0.805708 | 0.76416 | -0.82358 | -0.24315 | 0.759918 |
| L-Threonin | 0.933158 | 0.978465 | 0.964962 | 0.92422 | -0.82211 | -0.74567 | -0.70308 | -0.86231 | 0.942858 | 0.799754 | 0.684592 | -0.86518 | -0.28406 | 0.844636 |
| L-Tryptoph | 1 | 0.953454 | 0.969494 | 0.752563 | -0.75084 | -0.69051 | -0.60459 | -0.79396 | 0.873706 | 0.836386 | 0.761303 | -0.79427 | -0.24108 | 0.676064 |
| L-Tyrosine | 0.953454 | 1 | 0.963279 | 0.885557 | -0.82819 | -0.71116 | -0.70779 | -0.84732 | 0.96832 | 0.833484 | 0.631695 | -0.84818 | -0.3679 | 0.802817 |
| L-Valine | 0.969494 | 0.963279 | 1 | 0.82589 | -0.76096 | -0.71904 | -0.60093 | -0.79742 | 0.880734 | 0.815392 | 0.758152 | -0.7998 | -0.25728 | 0.759593 |
| Malic acid | 0.752563 | 0.885557 | 0.82589 | 1 | -0.7599 | -0.64593 | -0.72149 | -0.78941 | 0.906774 | 0.665535 | 0.468695 | -0.79342 | -0.33837 | 0.900585 |
| Myristic aci | -0.75084 | -0.82819 | -0.76096 | -0.7599 | 1 | 0.739657 | 0.824444 | 0.910715 | -0.85158 | -0.70685 | -0.47395 | 0.896717 | 0.293544 | -0.68282 |
| Niacinamid | -0.69051 | -0.71116 | -0.71904 | -0.64593 | 0.739657 | 1 | 0.632819 | 0.743821 | -0.67928 | -0.57674 | -0.66557 | 0.73341 | 0.186945 | -0.72871 |
| Oleic Acid | -0.60459 | -0.70779 | -0.60093 | -0.72149 | 0.824444 | 0.632819 | 1 | 0.883548 | -0.75679 | -0.61554 | -0.35257 | 0.865169 | 0.329056 | -0.71476 |
| Palmitic Ac | -0.79396 | -0.84732 | -0.79742 | -0.78941 | 0.910715 | 0.743821 | 0.883548 | 1 | -0.8454 | -0.65736 | -0.54746 | 0.998309 | 0.141091 | -0.77856 |
| Pantotheni | 0.873706 | 0.96832 | 0.880734 | 0.906774 | -0.85158 | -0.67928 | -0.75679 | -0.8454 | 1 | 0.827552 | 0.482275 | -0.8444 | -0.46343 | 0.787443 |
| Putrescine | 0.836386 | 0.833484 | 0.815392 | 0.665535 | -0.70685 | -0.57674 | -0.61554 | -0.65736 | 0.827552 | 1 | 0.585043 | -0.64249 | -0.42922 | 0.500457 |
| Serine | 0.761303 | 0.631695 | 0.758152 | 0.468695 | -0.47395 | -0.66557 | -0.35257 | -0.54746 | 0.482275 | 0.585043 | 1 | -0.53944 | 0.193209 | 0.398852 |
| Stearic aci | -0.79427 | -0.84818 | -0.7998 | -0.79342 | 0.896717 | 0.73341 | 0.865169 | 0.998309 | -0.8444 | -0.64249 | -0.53944 | 1 | 0.126406 | -0.7889 |
| Succinic ac | -0.24108 | -0.3679 | -0.25728 | -0.33837 | 0.293544 | 0.186945 | 0.329056 | 0.141091 | -0.46343 | -0.42922 | 0.193209 | 0.126406 | 1 | -0.29719 |
| Taurine | 0.676064 | 0.802817 | 0.759593 | 0.900585 | -0.68282 | -0.72871 | -0.71476 | -0.77856 | 0.787443 | 0.500457 | 0.398852 | -0.7889 | -0.29719 | 1 |

**Table S10: Siginificance value of correlation coefficient (pearson r) among metabolites**

| | BrdU | Monopalmi | adecenoic a | Adenine | Adenosine | nomalonic | AMP | achidonic a | Citric acid | Creatinine | Cystine | -(-)-Fructos | D-Lactose | DL-Ornithin | Glycine | Hypoxanthir | onicotinic A | -5-Oxoproli | Lactic Acid | -Asparagin | Aspartic ac | Glutamic ac | L-Glutamine | Linoleic aci | L-Isoleucine | L-Leucine | L-Lysine | Phenylalan | L-Threonine | -Tryptopha | L-Tyrosine | L-Valine | Malic acid | Myristic aci | Niacinamid | Oleic Acid | Palmitic Aci | ntothenic a | Putrescine | Serine | Stearic aci | Succinic aci | Taurine |
|---|---|---|---|---|---|---|---|---|---|---|---|---|---|---|---|---|---|---|---|---|---|---|---|---|---|---|---|---|---|---|---|---|---|---|---|---|---|---|---|---|---|---|---|
| BrdU | | 0.030628 | 0.634843 | 0.001434 | 0.443868 | 0.000118 | 2.7E-08 | 0.00222 | 6.3E-10 | 0.741866 | 0.003852 | 0.008909 | 0.0022 | 0.140125 | 0.008824 | 0.004571 | 0.111987 | 0.293603 | 0.053993 | 0.252926 | 0.040666 | 0.364225 | 0.023984 | 0.007324 | 0.152932 | 0.207294 | 0.927313 | 0.029287 | 0.002072 | 0.039552 | 0.002701 | 0.025592 | 1.2E-05 | 0.001169 | 0.12773 | 0.004002 | 0.004705 | 9.18E-05 | 0.026082 | 0.764028 | 0.004586 | 0.091262 | 0.01091 |
| 1-Monopal | 0.030628 | | 1.41E-05 | 0.002274 | 0.18853 | 0.000171 | 0.000294 | 1.51E-08 | 0.007611 | 0.032491 | 0.000227 | 2.97E-06 | 0.000409 | 0.011575 | 0.000172 | 2.54E-06 | 0.068988 | 0.125777 | 5.98E-05 | 0.218596 | 0.000485 | 0.202449 | 0.093446 | 1.6E-07 | 0.000702 | 0.003691 | 0.209058 | 0.000744 | 0.000159 | 0.000819 | 0.000118 | 0.001101 | 0.000233 | 0.000153 | 0.008285 | 1.79E-09 | 6.72E-07 | 9.68E-05 | 0.002342 | 0.111827 | 9.14E-07 | 0.115285 | 3.16E-05 |
| 9-Octadec | 0.634843 | 1.41E-05 | | 0.069519 | 0.692627 | 0.015021 | 0.063668 | 1.06E-05 | 0.370646 | 0.062811 | 0.009572 | 0.001689 | 0.015118 | 0.03389 | 0.006198 | 0.001694 | 0.221768 | 0.804212 | 0.007496 | 0.161661 | 0.023536 | 0.65892 | 0.085797 | 0.000132 | 0.018984 | 0.040073 | 0.05341 | 0.03163 | 0.025847 | 0.030778 | 0.030353 | 0.050495 | 0.06021 | 0.001985 | 0.002096 | 5.23E-08 | 5.41E-05 | 0.031931 | 0.026821 | 0.040298 | 0.000121 | 0.669372 | 0.025615 |
| Adenine | 0.001434 | 0.002274 | 0.069519 | | 0.006908 | 0.000485 | 0.0003 | 0.000677 | 0.003113 | 0.183817 | 0.00224 | 0.00363 | 0.000195 | 0.030333 | 0.001078 | 0.001662 | 0.214538 | 0.637322 | 0.036345 | 0.334667 | 0.027261 | 0.967647 | 0.024196 | 0.004431 | 0.056242 | 0.05014 | 0.212116 | 0.006444 | 0.000718 | 0.014269 | 0.002153 | 0.00904 | 0.000385 | 0.000177 | 0.163043 | 0.001155 | 2.01E-05 | 0.000837 | 0.035916 | 0.303667 | 1.49E-05 | 0.826314 | 0.021645 |
| Adenosine | 0.443868 | 0.18853 | 0.692627 | 0.006908 | | 0.932316 | 0.630622 | 0.57579 | 0.801914 | 0.747264 | 0.87572 | 0.512659 | 0.99792 | 0.987336 | 0.969612 | 0.372099 | 0.287599 | 0.858452 | 0.259896 | 0.134646 | 0.969541 | 0.02818 | 0.543249 | 0.305684 | 0.791981 | 0.717564 | 0.689875 | 0.779549 | 0.636714 | 0.566801 | 0.46489 | 0.778833 | 0.643165 | 0.677463 | 0.073862 | 0.318707 | 0.513431 | 0.327862 | 0.132633 | 0.333714 | 0.510418 | 0.675607 | 0.967583 |
| Aminomal | 0.000118 | 0.000171 | 0.015021 | 0.000485 | 0.932316 | | 1.18E-10 | 9.64E-09 | 1.68E-07 | 0.019108 | 6.35E-07 | 2.93E-06 | 1.35E-09 | 0.003315 | 4.24E-08 | 6.42E-07 | 0.051088 | 0.103402 | 0.000736 | 0.049591 | 1.87E-06 | 0.5726 | 0.009509 | 8.4E-07 | 9.43E-05 | 0.00076 | 0.202663 | 3.62E-05 | 1.59E-07 | 0.000634 | 1.75E-06 | 2.67E-05 | 1.08E-11 | 2.82E-06 | 0.000137 | 5.86E-06 | 2.85E-07 | 1.14E-07 | 0.003244 | 0.052672 | 2.55E-07 | 0.129969 | 1.32E-08 |
| AMP | 2.7E-08 | 0.000294 | 0.063668 | 0.0003 | 0.630622 | 1.18E-10 | | 2.86E-07 | 9.57E-10 | 0.08925 | 1.7E-07 | 2.87E-06 | 1.38E-08 | 0.008755 | 4.43E-06 | 2.36E-06 | 0.034548 | 0.074454 | 0.000535 | 0.040818 | 7.55E-05 | 0.27623 | 0.002808 | 8.54E-06 | 0.000519 | 0.003112 | 0.434485 | 5.61E-05 | 2.35E-07 | 0.00027 | 6.13E-07 | 6.63E-05 | 5.55E-12 | 6.28E-06 | 0.002335 | 2.77E-05 | 3E-06 | 3.08E-09 | 0.004178 | 0.181409 | 2.33E-06 | 0.052221 | 5.82E-07 |
| Arachidoni | 0.00222 | 1.51E-08 | 1.06E-05 | 0.000677 | 0.57579 | 9.64E-09 | 2.86E-07 | | 0.000338 | 0.005286 | 4.94E-08 | 1.26E-09 | 1.26E-08 | 0.000394 | 1.28E-08 | 2.17E-10 | 0.040689 | 0.07595 | 4.7E-06 | 0.056769 | 2.41E-06 | 0.40474 | 0.021023 | 5.56E-11 | 1.09E-05 | 0.00015 | 0.038266 | 2.89E-06 | 6.6E-08 | 1.51E-05 | 1.03E-07 | 6.26E-06 | 1.59E-07 | 5.16E-08 | 4.02E-05 | 1.17E-12 | 1.17E-11 | 2.53E-08 | 0.00017 | 0.011247 | 5.2E-11 | 0.119476 | 1.62E-06 |
| Citric acid | 6.3E-10 | 0.007611 | 0.370646 | 0.003113 | 0.801914 | 1.68E-07 | 9.57E-10 | 0.000338 | | 0.411045 | 0.000125 | 0.002063 | 6.75E-05 | 0.221412 | 0.001857 | 0.001365 | 0.048075 | 0.160051 | 0.017785 | 0.162516 | 0.004982 | 0.39711 | 0.016092 | 0.00047 | 0.009721 | 0.04815 | 0.715136 | 0.007367 | 0.000322 | 0.032364 | 0.000637 | 0.006478 | 2.64E-08 | 0.001833 | 0.039055 | 0.002554 | 0.002173 | 8.61E-05 | 0.119136 | 0.737577 | 0.001766 | 0.076926 | 3.46E-06 |
| Creatinine | 0.741866 | 0.032491 | 0.062811 | 0.183817 | 0.747264 | 0.019108 | 0.08925 | 0.005286 | 0.411045 | | 0.000262 | 0.000564 | 0.001904 | 7.45E-09 | 2.97E-06 | 0.000263 | 0.183172 | 0.043531 | 0.004811 | 0.343723 | 6.37E-06 | 0.383045 | 0.453004 | 0.030949 | 5.86E-09 | 1.39E-12 | 4E-07 | 8.34E-08 | 4.07E-05 | 6.28E-07 | 7.16E-05 | 3.82E-08 | 0.009819 | 0.010564 | 0.001637 | 0.073589 | 0.001995 | 0.004595 | 0.000483 | 2.98E-07 | 0.001993 | 0.863936 | 0.018903 |
| Cystine | 0.003852 | 0.000227 | 0.009572 | 0.00224 | 0.87572 | 6.35E-07 | 1.7E-07 | 4.94E-08 | 0.000125 | 0.000262 | | 2.58E-09 | 5.51E-10 | 3.23E-05 | 3.55E-09 | 2.48E-08 | 0.274606 | 0.039707 | 2.88E-05 | 0.102843 | 4.55E-06 | 0.356631 | 0.044862 | 6.53E-05 | 4.62E-09 | 4.19E-07 | 0.004876 | 1.39E-09 | 1.05E-09 | 3.67E-08 | 5.86E-10 | 9.88E-09 | 8.03E-08 | 1.19E-07 | 5.24E-05 | 3.42E-05 | 3.28E-08 | 9.89E-09 | 0.000221 | 0.002663 | 4.37E-08 | 0.164453 | 2.7E-06 |
| D-(-)-Fruct | 0.008909 | 2.97E-06 | 0.001689 | 0.00363 | 0.512659 | 2.93E-06 | 2.87E-06 | 1.26E-09 | 0.002063 | 0.000564 | 2.58E-09 | | 1.01E-08 | 1.41E-05 | 6.08E-09 | 7.42E-12 | 0.077856 | 0.010243 | 4.56E-09 | 0.081201 | 2.2E-07 | 0.094816 | 0.12477 | 4.61E-06 | 2.95E-07 | 2.6E-06 | 0.007329 | 3.4E-09 | 1.28E-09 | 7.19E-09 | 5.78E-11 | 2.52E-08 | 8.6E-07 | 1.83E-06 | 4.33E-05 | 2.13E-06 | 8.74E-09 | 1.65E-10 | 1.03E-05 | 0.002526 | 1.25E-08 | 0.129959 | 2.33E-05 |
| D-Lactose | 0.0022 | 0.000409 | 0.015118 | 0.000195 | 0.99792 | 1.35E-09 | 1.38E-08 | 1.26E-08 | 6.75E-05 | 0.001904 | 5.51E-10 | 1.01E-08 | | 5.35E-05 | 6.67E-10 | 5.49E-08 | 0.063764 | 0.135371 | 0.00018 | 0.018273 | 1.12E-06 | 0.681089 | 0.006334 | 2.05E-05 | 2.2E-06 | 1.3E-05 | 0.01426 | 4.95E-08 | 7.43E-10 | 2.03E-06 | 2.99E-09 | 1.75E-07 | 1.92E-08 | 1.26E-07 | 4.45E-05 | 3.17E-05 | 9.78E-10 | 2.61E-09 | 0.001845 | 0.004633 | 6.28E-10 | 0.317636 | 5.52E-07 |
| DL-Ornithi | 0.140125 | 0.011575 | 0.03389 | 0.030333 | 0.987336 | 0.003315 | 0.008755 | 0.000394 | 0.221412 | 7.45E-09 | 3.23E-05 | 1.41E-05 | 5.35E-05 | | 5.27E-09 | 1.7E-06 | 0.592165 | 0.02534 | 0.000148 | 0.050589 | 1.59E-07 | 0.237736 | 0.954042 | 0.005567 | 1.63E-07 | 2.45E-10 | 1.15E-07 | 5.3E-11 | 2.84E-08 | 2.12E-11 | 1.49E-07 | 9.02E-11 | 0.000588 | 0.000168 | 6.58E-05 | 0.013128 | 4.05E-05 | 3.13E-05 | 2.9E-06 | 6.66E-09 | 4.17E-05 | 0.777724 | 0.00628 |
| Glycine | 0.008824 | 0.000172 | 0.006198 | 0.001078 | 0.969612 | 4.24E-08 | 4.43E-06 | 1.28E-08 | 0.001857 | 2.97E-06 | 3.55E-09 | 6.08E-09 | 6.67E-10 | 5.27E-09 | | 7E-11 | 0.564432 | 0.038288 | 1.82E-05 | 0.057976 | 3.04E-10 | 0.365809 | 0.199904 | 1.16E-05 | 5.99E-09 | 2.73E-09 | 0.000304 | 1.22E-12 | 2.47E-14 | 2.44E-09 | 4.31E-11 | 1E-12 | 2.54E-08 | 1.15E-07 | 3.23E-06 | 4.78E-05 | 2.55E-09 | 7.83E-09 | 6.24E-06 | 4.3E-05 | 2.75E-09 | 0.355155 | 2.46E-06 |
| Hypoxanth | 0.004571 | 2.54E-06 | 0.001694 | 0.001662 | 0.372099 | 6.42E-07 | 2.36E-06 | 2.17E-10 | 0.001365 | 0.000263 | 2.48E-08 | 7.42E-12 | 5.49E-08 | 1.7E-06 | 7E-11 | | 0.126742 | 0.004625 | 3.36E-08 | 0.052729 | 2.12E-09 | 0.057711 | 0.223905 | 2.59E-08 | 1.43E-07 | 1.75E-06 | 0.005997 | 4.63E-10 | 1.82E-12 | 2.06E-09 | 5.37E-13 | 7.29E-10 | 2.88E-08 | 1.03E-06 | 2.01E-05 | 1.17E-06 | 5.5E-09 | 1.44E-11 | 2.76E-07 | 0.000867 | 7.25E-09 | 0.146484 | 6.38E-06 |
| Isonicotini | 0.111987 | 0.068988 | 0.221768 | 0.214538 | 0.287599 | 0.051088 | 0.034548 | 0.040689 | 0.048075 | 0.183172 | 0.274606 | 0.077856 | 0.063764 | 0.592165 | 0.564432 | 0.126742 | | 0.193365 | 0.093975 | 0.666934 | 0.500627 | 0.176243 | 0.010448 | 0.008517 | 0.928504 | 0.493785 | 0.174273 | 0.741954 | 0.400146 | 0.727653 | 0.171637 | 0.802947 | 0.153857 | 0.074665 | 0.48729 | 0.013221 | 0.112001 | 0.028182 | 0.364974 | 0.158949 | 0.120792 | 0.025634 | 0.257539 |
| L-5-Oxopr | 0.293603 | 0.125777 | 0.804212 | 0.637322 | 0.858452 | 0.103402 | 0.074454 | 0.07595 | 0.160051 | 0.043531 | 0.039707 | 0.010243 | 0.135371 | 0.02534 | 0.038288 | 0.004625 | 0.193365 | | 0.001303 | 0.895284 | 0.001652 | 9.34E-08 | 0.180054 | 0.012709 | 0.005669 | 0.025345 | 0.391302 | 0.005188 | 0.007547 | 0.002866 | 0.001183 | 0.002985 | 0.017849 | 0.157437 | 0.064249 | 0.221589 | 0.344668 | 0.001697 | 0.00014 | 0.146134 | 0.367121 | 0.000138 | 0.055646 |
| Lactic Acid | 0.053993 | 5.98E-05 | 0.007496 | 0.036345 | 0.259896 | 0.000736 | 0.000535 | 4.7E-06 | 0.017785 | 0.004811 | 2.88E-05 | 4.56E-09 | 0.00018 | 0.000148 | 1.82E-05 | 3.36E-08 | 0.093975 | 0.001303 | | 0.093259 | 1.61E-06 | 0.005114 | 0.840948 | 2.54E-05 | 2.87E-05 | 0.000178 | 0.018071 | 5.57E-06 | 3.16E-06 | 1.13E-06 | 2.55E-07 | 8.71E-06 | 4.85E-05 | 0.000292 | 0.004912 | 4.96E-05 | 8.57E-05 | 5.7E-07 | 3.04E-06 | 0.001831 | 0.000128 | 0.187653 | 0.000847 |
| L-Asparag | 0.252926 | 0.218596 | 0.161661 | 0.334667 | 0.134646 | 0.049591 | 0.040818 | 0.056769 | 0.162516 | 0.343723 | 0.102843 | 0.081201 | 0.018273 | 0.050589 | 0.057976 | 0.052729 | 0.666934 | 0.895284 | 0.093259 | | 0.013046 | 0.441175 | 0.07847 | 0.152429 | 0.100912 | 0.100917 | 0.436532 | 0.079697 | 0.03518 | 0.081431 | 0.086445 | 0.111831 | 0.02643 | 0.272588 | 0.018123 | 0.146385 | 0.051416 | 0.106477 | 0.67264 | 0.012361 | 0.044694 | 0.180395 | 0.043438 |
| L-Aspartic | 0.040666 | 0.000485 | 0.023536 | 0.027261 | 0.969541 | 1.87E-06 | 7.55E-05 | 2.41E-06 | 0.004982 | 6.37E-06 | 4.55E-06 | 2.2E-07 | 1.12E-06 | 1.59E-07 | 3.04E-10 | 2.12E-09 | 0.500627 | 0.001652 | 1.61E-06 | 0.013046 | | 0.059542 | 0.732371 | 1.55E-05 | 4.96E-09 | 4.39E-08 | 0.005744 | 1.2E-09 | 1.1E-10 | 5.8E-08 | 2.56E-09 | 2.41E-10 | 5.66E-08 | 0.000498 | 9.86E-05 | 0.000701 | 2.93E-05 | 2.63E-07 | 1.19E-05 | 2.05E-05 | 2.86E-05 | 0.228104 | 8.41E-06 |
| L-Glutamic | 0.364225 | 0.202449 | 0.65892 | 0.967647 | 0.02818 | 0.5726 | 0.27623 | 0.40474 | 0.39711 | 0.383045 | 0.356631 | 0.094816 | 0.681089 | 0.237736 | 0.365809 | 0.057711 | 0.176243 | 9.34E-08 | 0.005114 | 0.441175 | 0.059542 | | 0.048797 | 0.079874 | 0.166691 | 0.318643 | 0.956849 | 0.102158 | 0.102702 | 0.04273 | 0.026723 | 0.071899 | 0.132875 | 0.589307 | 0.891859 | 0.505178 | 0.838368 | 0.022511 | 0.001376 | 0.828842 | 0.850216 | 0.001587 | 0.208026 |
| L-Glutamin | 0.023984 | 0.093446 | 0.085797 | 0.024196 | 0.543249 | 0.009509 | 0.002808 | 0.021023 | 0.016092 | 0.453004 | 0.044862 | 0.12477 | 0.006334 | 0.954042 | 0.199904 | 0.223905 | 0.010448 | 0.180054 | 0.840948 | 0.07847 | 0.732371 | 0.048797 | | 0.104265 | 0.680112 | 0.915791 | 0.696015 | 0.512435 | 0.237503 | 0.596681 | 0.268196 | 0.644179 | 0.091454 | 0.012314 | 0.085443 | 0.012919 | 0.005931 | 0.084994 | 0.774959 | 0.502828 | 0.005233 | 0.687934 | 0.118261 |
| Linoleic ac | 0.007324 | 1.6E-07 | 0.000132 | 0.004431 | 0.305684 | 8.4E-07 | 8.54E-06 | 5.56E-11 | 0.00047 | 0.030949 | 6.53E-05 | 4.61E-06 | 2.05E-05 | 0.005567 | 1.16E-05 | 2.59E-08 | 0.008517 | 0.012709 | 2.54E-05 | 0.152429 | 1.55E-05 | 0.079874 | 0.104265 | | 0.000302 | 0.004007 | 0.230581 | 0.000163 | 6.36E-06 | 0.000409 | 3.02E-06 | 0.000145 | 1.97E-06 | 1.05E-05 | 0.001531 | 3.59E-10 | 1.56E-06 | 4.66E-07 | 0.000119 | 0.072335 | 3.03E-06 | 0.019894 | 1.03E-05 |
| L-Isoleucin | 0.152932 | 0.000702 | 0.018984 | 0.056242 | 0.791981 | 9.43E-05 | 0.000519 | 1.09E-05 | 0.009721 | 5.86E-09 | 4.62E-09 | 2.95E-07 | 2.2E-06 | 1.63E-07 | 5.99E-09 | 1.43E-07 | 0.928504 | 0.005669 | 2.87E-05 | 0.100912 | 4.96E-09 | 0.166691 | 0.680112 | 0.000302 | | 1.52E-14 | 0.00031 | 5.13E-12 | 7.16E-09 | 7.02E-10 | 5.23E-09 | 3.24E-12 | 1.22E-05 | 0.000298 | 0.000118 | 0.001843 | 1.87E-05 | 7.6E-06 | 0.000152 | 1.18E-05 | 1.87E-05 | 0.278121 | 1.43E-05 |
| L-Leucine | 0.207294 | 0.003691 | 0.040073 | 0.05014 | 0.717564 | 0.00076 | 0.003112 | 0.00015 | 0.04815 | 1.39E-12 | 4.19E-07 | 2.6E-06 | 1.3E-05 | 2.45E-10 | 2.73E-09 | 1.75E-06 | 0.493785 | 0.025345 | 0.000178 | 0.100917 | 4.39E-08 | 0.318643 | 0.915791 | 0.004007 | 1.52E-14 | | 6.16E-06 | 8.84E-13 | 2.23E-08 | 1.59E-10 | 8.94E-08 | 6.86E-13 | 0.000136 | 0.000621 | 0.000128 | 0.010784 | 4.18E-05 | 6.65E-05 | 0.000211 | 2.73E-07 | 3.75E-05 | 0.646565 | 0.000294 |
| L-Lysine | 0.927313 | 0.209058 | 0.05341 | 0.212116 | 0.689875 | 0.202663 | 0.434485 | 0.038266 | 0.715136 | 4E-07 | 0.004876 | 0.007329 | 0.01426 | 1.15E-07 | 0.000304 | 0.005997 | 0.174273 | 0.391302 | 0.018071 | 0.436532 | 0.005744 | 0.956849 | 0.696015 | 0.230581 | 0.00031 | 6.16E-06 | | 8.83E-05 | 0.003117 | 4.8E-05 | 0.003382 | 0.000126 | 0.187437 | 0.004243 | 0.001837 | 0.143425 | 0.004106 | 0.03925 | 0.0032 | 1.12E-07 | 0.004942 | 0.358867 | 0.405973 |
| L-Phenylal | 0.029287 | 0.000744 | 0.03163 | 0.006444 | 0.779549 | 3.62E-05 | 5.61E-05 | 2.89E-06 | 0.007367 | 8.34E-08 | 1.39E-09 | 3.4E-09 | 4.95E-08 | 5.3E-11 | 1.22E-12 | 4.63E-10 | 0.741954 | 0.005188 | 5.57E-06 | 0.079697 | 1.2E-09 | 0.102158 | 0.512435 | 0.000163 | 5.13E-12 | 8.84E-13 | 8.83E-05 | | 1.07E-14 | 1.33E-16 | 1.82E-14 | 1.01E-22 | 9.16E-07 | 6.94E-06 | 3.57E-05 | 0.001135 | 8.28E-07 | 8.09E-09 | 2.03E-06 | 1.38E-05 | 7.66E-07 | 0.252263 | 4.08E-05 |
| L-Threonin | 0.002072 | 0.000159 | 0.025847 | 0.000718 | 0.636714 | 1.59E-07 | 2.35E-07 | 6.6E-08 | 0.000322 | 4.07E-05 | 1.05E-09 | 1.28E-09 | 7.43E-10 | 2.84E-08 | 2.47E-14 | 1.82E-12 | 0.400146 | 0.007547 | 3.16E-06 | 0.03518 | 1.1E-10 | 0.102702 | 0.237503 | 6.36E-06 | 7.16E-09 | 2.23E-08 | 0.003117 | 1.07E-14 | | 3E-11 | 1.44E-16 | 2.86E-14 | 1.15E-10 | 8.33E-07 | 2.89E-05 | 0.000127 | 6.06E-08 | 5.6E-12 | 2.74E-06 | 0.000224 | 4.88E-08 | 0.178552 | 7.66E-07 |
| L-Tryptoph | 0.039552 | 0.000819 | 0.030778 | 0.014269 | 0.566801 | 0.000634 | 0.00027 | 1.51E-05 | 0.032364 | 6.28E-07 | 3.67E-08 | 7.19E-09 | 2.03E-06 | 2.12E-11 | 2.44E-09 | 2.06E-09 | 0.727653 | 0.002866 | 1.13E-06 | 0.081431 | 5.8E-08 | 0.04273 | 0.596681 | 0.000409 | 7.02E-10 | 1.59E-10 | 4.8E-05 | 1.33E-16 | 3E-11 | | 6.17E-13 | 6.37E-15 | 2.21E-05 | 2.37E-05 | 0.000188 | 0.001752 | 3.64E-06 | 2.48E-08 | 3.56E-07 | 1.56E-05 | 3.59E-06 | 0.256451 | 0.000553 |
| L-Tyrosine | 0.002701 | 0.000118 | 0.030353 | 0.002153 | 0.46489 | 1.75E-06 | 6.13E-07 | 1.03E-07 | 0.000637 | 7.16E-05 | 5.86E-10 | 5.78E-11 | 2.99E-09 | 1.49E-07 | 4.31E-11 | 5.37E-13 | 0.171637 | 0.001183 | 2.55E-07 | 0.086445 | 2.56E-09 | 0.026723 | 0.268196 | 3.02E-06 | 5.23E-09 | 8.94E-08 | 0.003382 | 1.82E-14 | 1.44E-16 | 6.17E-13 | | 4.76E-14 | 8.87E-09 | 5.85E-07 | 9.8E-05 | 0.000109 | 1.76E-07 | 9.6E-15 | 4.26E-07 | 0.00093 | 1.66E-07 | 0.076932 | 6.91E-06 |
| L-Valine | 0.025592 | 0.001101 | 0.050495 | 0.00904 | 0.778833 | 2.67E-05 | 6.63E-05 | 6.26E-06 | 0.006478 | 3.82E-08 | 9.88E-09 | 2.52E-08 | 1.75E-07 | 9.02E-11 | 1E-12 | 7.29E-10 | 0.802947 | 0.002985 | 8.71E-06 | 0.111831 | 2.41E-10 | 0.071899 | 0.644179 | 0.000145 | 3.24E-12 | 6.86E-13 | 0.000126 | 1.01E-22 | 2.86E-14 | 6.37E-15 | 4.76E-14 | | 6.7E-07 | 1.58E-05 | 7.53E-05 | 0.0019 | 3.08E-06 | 1.37E-08 | 1.21E-06 | 1.77E-05 | 2.73E-06 | 0.224868 | 4.13E-05 |
| Malic acid | 1.2E-05 | 0.000233 | 0.06021 | 0.000385 | 0.643165 | 1.08E-11 | 5.55E-12 | 1.59E-07 | 2.64E-08 | 0.009819 | 8.03E-08 | 8.6E-07 | 1.92E-08 | 0.000588 | 2.54E-08 | 2.88E-08 | 0.153857 | 0.017849 | 4.85E-05 | 0.02643 | 5.66E-08 | 0.132875 | 0.091454 | 1.97E-06 | 1.22E-05 | 0.000136 | 0.187437 | 9.16E-07 | 1.15E-10 | 2.21E-05 | 8.87E-09 | 6.7E-07 | | 1.65E-05 | 0.000651 | 6.92E-05 | 4.53E-06 | 1.03E-09 | 0.000386 | 0.020878 | 3.74E-06 | 0.105819 | 1.12E-08 |
| Myristic ac | 0.001169 | 0.000153 | 0.001985 | 0.000177 | 0.677463 | 2.82E-06 | 6.28E-06 | 5.16E-08 | 0.001833 | 0.010564 | 1.19E-07 | 1.83E-06 | 1.26E-07 | 0.000168 | 1.15E-07 | 1.03E-06 | 0.074665 | 0.157437 | 0.000292 | 0.272588 | 0.000498 | 0.589307 | 0.012314 | 1.05E-05 | 0.000298 | 0.000621 | 0.004243 | 6.94E-06 | 8.33E-07 | 2.37E-05 | 5.85E-07 | 1.58E-05 | 1.65E-05 | | 3.62E-05 | 7.29E-07 | 6.52E-10 | 1.31E-07 | 0.000113 | 0.019301 | 3.03E-09 | 0.163865 | 0.000462 |
| Niacinamid | 0.12773 | 0.008285 | 0.002096 | 0.163043 | 0.073862 | 0.000137 | 0.002335 | 4.02E-05 | 0.039055 | 0.001637 | 5.24E-05 | 4.33E-05 | 4.45E-05 | 6.58E-05 | 3.23E-06 | 2.01E-05 | 0.48729 | 0.064249 | 0.004912 | 0.018123 | 9.86E-05 | 0.891859 | 0.085443 | 0.001531 | 0.000118 | 0.000128 | 0.001837 | 3.57E-05 | 2.89E-05 | 0.000188 | 9.8E-05 | 7.53E-05 | 0.000651 | 3.62E-05 | | 0.000905 | 3.1E-05 | 0.000262 | 0.003175 | 0.000386 | 4.55E-05 | 0.381739 | 0.00012 |
| Oleic Acid | 0.004002 | 1.79E-09 | 5.23E-08 | 0.001155 | 0.318707 | 5.86E-06 | 2.77E-05 | 1.17E-12 | 0.002554 | 0.073589 | 3.42E-05 | 2.13E-06 | 3.17E-05 | 0.013128 | 4.78E-05 | 1.17E-06 | 0.013221 | 0.221589 | 4.96E-05 | 0.146385 | 0.000701 | 0.505178 | 0.012919 | 3.59E-10 | 0.001843 | 0.010784 | 0.143425 | 0.001135 | 0.000127 | 0.001752 | 0.000109 | 0.0019 | 6.92E-05 | 7.29E-07 | 0.000905 | | 1.06E-08 | 1.87E-05 | 0.001366 | 0.091073 | 4.88E-08 | 0.116398 | 0.000186 |
| Palmitic A | 0.004705 | 6.72E-07 | 5.41E-05 | 2.01E-05 | 0.513431 | 2.85E-07 | 3E-06 | 1.17E-11 | 0.002173 | 0.001995 | 3.28E-08 | 8.74E-09 | 9.78E-10 | 4.05E-05 | 2.55E-09 | 5.5E-09 | 0.112001 | 0.344668 | 8.57E-05 | 0.051416 | 2.93E-05 | 0.838368 | 0.005931 | 1.56E-06 | 1.87E-05 | 4.18E-05 | 0.004106 | 8.28E-07 | 6.06E-08 | 3.64E-06 | 1.76E-07 | 3.08E-06 | 4.53E-06 | 6.52E-10 | 3.1E-05 | 1.06E-08 | | 2E-07 | 0.000482 | 0.005624 | 1.11E-28 | 0.510792 | 1.98E-05 |
| Pantotheni | 9.18E-05 | 9.68E-05 | 0.031931 | 0.000837 | 0.327862 | 1.14E-07 | 3.08E-09 | 2.53E-08 | 8.61E-05 | 0.004595 | 9.89E-09 | 1.65E-10 | 2.61E-09 | 3.13E-05 | 7.83E-09 | 1.44E-11 | 0.028182 | 0.001697 | 5.7E-07 | 0.106477 | 2.63E-07 | 0.022511 | 0.084994 | 4.66E-07 | 7.6E-06 | 6.65E-05 | 0.03925 | 8.09E-09 | 5.6E-12 | 2.48E-08 | 9.6E-15 | 1.37E-08 | 1.03E-09 | 1.31E-07 | 0.000262 | 1.87E-05 | 2E-07 | | 6.08E-07 | 0.017002 | 2.13E-07 | 0.022561 | 1.37E-05 |
| Putrescine | 0.026082 | 0.002342 | 0.026821 | 0.035916 | 0.132633 | 0.003244 | 0.004178 | 0.00017 | 0.119136 | 0.000483 | 0.000221 | 1.03E-05 | 0.001845 | 2.9E-06 | 6.24E-06 | 2.76E-07 | 0.364974 | 0.00014 | 3.04E-06 | 0.67264 | 1.19E-05 | 0.001376 | 0.774959 | 0.000119 | 0.000152 | 0.000211 | 0.0032 | 2.03E-06 | 2.74E-06 | 3.56E-07 | 4.26E-07 | 1.21E-06 | 0.000386 | 0.000113 | 0.003175 | 0.001366 | 0.000482 | 6.08E-07 | | 0.002674 | 0.000711 | 0.036348 | 0.017686 |
| Serine | 0.764028 | 0.111827 | 0.040298 | 0.303667 | 0.333714 | 0.052672 | 0.181409 | 0.011247 | 0.737577 | 2.98E-07 | 0.002663 | 0.002526 | 0.004633 | 6.66E-09 | 4.3E-05 | 0.000867 | 0.158949 | 0.146134 | 0.001831 | 0.012361 | 2.05E-05 | 0.828842 | 0.502828 | 0.072335 | 1.18E-05 | 2.73E-07 | 1.12E-07 | 1.38E-05 | 0.000224 | 1.56E-05 | 0.00093 | 1.77E-05 | 0.020878 | 0.019301 | 0.000386 | 0.091073 | 0.005624 | 0.017002 | 0.002674 | | 0.00652 | 0.365696 | 0.065953 |
| Stearic aci | 0.004586 | 9.14E-07 | 0.000121 | 1.49E-05 | 0.510418 | 2.55E-07 | 2.33E-06 | 5.2E-11 | 0.001766 | 0.001993 | 4.37E-08 | 1.25E-08 | 6.28E-10 | 4.17E-05 | 2.75E-09 | 7.25E-09 | 0.120792 | 0.367121 | 0.000128 | 0.044694 | 2.86E-05 | 0.850216 | 0.005233 | 3.03E-06 | 1.87E-05 | 3.75E-05 | 0.004942 | 7.66E-07 | 4.88E-08 | 3.59E-06 | 1.66E-07 | 2.73E-06 | 3.74E-06 | 3.03E-09 | 4.55E-05 | 4.88E-08 | 1.11E-28 | 2.13E-07 | 0.000711 | 0.00652 | | 0.556148 | 1.28E-05 |
| Succinic a | 0.091262 | 0.115285 | 0.669372 | 0.826314 | 0.675607 | 0.129969 | 0.052221 | 0.119476 | 0.076926 | 0.863936 | 0.164453 | 0.129959 | 0.317636 | 0.777724 | 0.355155 | 0.146484 | 0.025634 | 0.000138 | 0.187653 | 0.180395 | 0.228104 | 0.001587 | 0.687934 | 0.019894 | 0.278121 | 0.646565 | 0.358867 | 0.252263 | 0.178552 | 0.256451 | 0.076932 | 0.224868 | 0.105819 | 0.163865 | 0.381739 | 0.116398 | 0.510792 | 0.022561 | 0.036348 | 0.365696 | 0.556148 | | 0.179216 |
| Taurine | 0.01091 | 3.16E-05 | 0.025615 | 0.021645 | 0.967583 | 1.32E-08 | 5.82E-07 | 1.62E-06 | 3.46E-06 | 0.018903 | 2.7E-06 | 2.33E-05 | 5.52E-07 | 0.00628 | 2.46E-06 | 6.38E-06 | 0.257539 | 0.055646 | 0.000847 | 0.043438 | 8.41E-06 | 0.208026 | 0.118261 | 1.03E-05 | 1.43E-05 | 0.000294 | 0.405973 | 4.08E-05 | 7.66E-07 | 0.000553 | 6.91E-06 | 4.13E-05 | 1.12E-08 | 0.000462 | 0.00012 | 0.000186 | 1.98E-05 | 1.37E-05 | 0.017686 | 0.065953 | 1.28E-05 | 0.179216 | |

**Table S11: Correlation between cMYC and metabolites**

| Name | Pearson r | p value |
|---|---|---|
| C-MYC mRNA | 1 | |
| AMP | 0.954247823 | 5.12482E-13 |
| Aminomalonic acid | 0.917091909 | 2.97619E-10 |
| Malic acid | 0.89173314 | 4.96498E-09 |
| Citric acid | 0.874051793 | 2.40731E-08 |
| Pantothenic acid | 0.872948798 | 2.63552E-08 |
| Cystine | 0.852093038 | 1.26729E-07 |
| Taurine | 0.83091785 | 1.68124E-06 |
| L-Threonine | 0.818653194 | 1.01165E-06 |
| L-Tyrosine | 0.803263776 | 2.29531E-06 |
| Glycine | 0.787805917 | 4.88073E-06 |
| Adenine | 0.758209488 | 1.76631E-05 |
| L-Phenylalanine | 0.713334563 | 9.11598E-05 |
| L-Valine | 0.696497318 | 0.000156381 |
| L-Glutamine | 0.658996106 | 0.000461729 |
| L-Aspartic acid | 0.65004041 | 0.000585301 |
| L-Tryptophan | 0.643856348 | 0.000686449 |
| L-Isoleucine | 0.628592186 | 0.001002856 |
| L-Leucine | 0.557574471 | 0.004642945 |
| Putrescine | 0.531033979 | 0.007584534 |
| DL-Ornithine | 0.497974936 | 0.013273114 |
| Isonicotinic Acid | 0.476924456 | 0.018451999 |
| Creatinine | 0.356781118 | 0.087006899 |
| L-Asparagine | 0.315997385 | 0.132510574 |
| Serine | 0.225902778 | 0.288502979 |
| L-Lysine | 0.199733786 | 0.349410974 |
| Adenosine | 0.135874046 | 0.526696948 |
| L-Glutamic acid | -0.126531806 | 0.555752935 |
| L-5-Oxoproline | -0.266034137 | 0.208930005 |
| Succinic acid | -0.369129628 | 0.075877866 |
| 9-Octadecenoic acid(E) | -0.430197331 | 0.035876519 |
| Lactic Acid | -0.571980435 | 0.003496855 |
| Niacinamide | -0.612681032 | 0.001458746 |
| 1-Monopalmitin | -0.690162983 | 0.000189838 |
| Linoleic acid | -0.759334089 | 1.68767E-05 |
| Oleic Acid | -0.762121063 | 1.50597E-05 |
| Hypoxanthine | -0.783806689 | 5.87392E-06 |
| D-(-)-Fructose | -0.79567163 | 3.35127E-06 |
| Myristic acid | -0.832630149 | 4.48664E-07 |
| Arachidonic acid | -0.844943747 | 2.0567E-07 |
| Palmitic Acid | -0.853506395 | 1.14817E-07 |
| Stearic acid | -0.860090385 | 7.14962E-08 |
| D-Lactose | -0.902105179 | 1.72354E-09 |

| |
|---|
| **Positive correlation** |
| **Negative correlation** |

**Table S12: Correlation between cMYC and other gene expression**

| Name | Pearsor r | p value |
|---|---|---|
| MYC | 1 | |
| SHMT2 | 0.85 | 1.58E-07 |
| LAT1 | 0.83 | 8.76E-07 |
| APRT | 0.82 | 1.14E-03 |
| LACC1 | 0.82 | 1.19E-03 |
| BCAT1 | 0.81 | 1.56E-03 |
| GLS1 | 0.80 | 1.75E-03 |
| AHCYL1 | 0.80 | 1.78E-03 |
| SLC1A5 | 0.79 | 4.07E-06 |
| MTR | 0.79 | 2.17E-03 |
| GPT2 | 0.77 | 3.12E-03 |
| SLC16A1 | 0.77 | 1.22E-05 |
| SMOX | 0.76 | 1.50E-05 |
| ADA | 0.76 | 4.34E-03 |
| SLC7A1 | 0.76 | 1.93E-05 |
| SLC3A2 | 0.74 | 3.95E-05 |
| MTAP | 0.73 | 4.90E-05 |
| GCLC | 0.73 | 6.02E-05 |
| GPT1 | 0.69 | 1.29E-02 |
| SLC2A3/GLUT3 | 0.69 | 2.19E-04 |
| SLC43A1 | 0.68 | 3.20E-04 |
| SLC16A10 | 0.68 | 5.50E-04 |
| ACC2 | 0.65 | 2.28E-02 |
| GCLM | 0.65 | 6.56E-04 |
| GOT1 | 0.63 | 2.65E-02 |
| G6PDH | 0.63 | 2.82E-02 |
| AMD1 | 0.62 | 1.10E-03 |
| SLC7A11 | 0.62 | 1.12E-03 |
| MAT2A | 0.62 | 1.22E-03 |
| HPRT | 0.60 | 3.86E-02 |
| ACC1 | 0.60 | 3.97E-02 |
| GLS2 | 0.57 | 5.45E-02 |
| SRM | 0.49 | 1.58E-02 |
| SHMT1 | 0.45 | 2.56E-02 |
| PAOX | 0.43 | 5.27E-02 |
| SLC16A3 | 0.43 | 3.73E-02 |
| ODC1 | 0.33 | 1.10E-01 |
| SLC2A1/GLUT1 | 0.31 | 1.37E-01 |
| SLC43A2 | 0.29 | 1.75E-01 |
| GOT2 | 0.26 | 4.12E-01 |
| ADK | 0.21 | 5.16E-01 |
| SSAT1 | -0.10 | 6.40E-01 |
| SMS | -0.29 | 1.65E-01 |

| |
|---|
| **Positive correlation** |
| **Negative correlation** |

**Table S13: Enriched pathay in 4B compared to 3B**

| Name | Total | Expected | Hits | Raw p | FDR |
|---|---|---|---|---|---|
| **Aminoacyl-tRNA biosynthesis** | **48** | **0.77419** | **10** | **7.09E-10** | **5.96E-08** |
| **Pantothenate and CoA biosynthesis** | **19** | **0.30645** | **5** | **7.16E-06** | **0.000301** |
| **Valine, leucine and isoleucine biosynthesis** | **8** | **0.12903** | **3** | **0.000197** | **0.005518** |
| **Phenylalanine, tyrosine and tryptophan biosynthesis** | **4** | **0.064516** | **2** | **0.00147** | **0.030867** |
| **Biosynthesis of unsaturated fatty acids** | **36** | **0.58065** | **4** | **0.002192** | **0.036824** |
| Taurine and hypotaurine metabolism | 8 | 0.12903 | 2 | 0.006592 | 0.092292 |
| Glutathione metabolism | 28 | 0.45161 | 3 | 0.00931 | 0.096945 |
| Alanine, aspartate and glutamate metabolism | 28 | 0.45161 | 3 | 0.00931 | 0.096945 |
| Phenylalanine metabolism | 10 | 0.16129 | 2 | 0.010387 | 0.096945 |
| Glycine, serine and threonine metabolism | 33 | 0.53226 | 3 | 0.014702 | 0.12349 |
| beta-Alanine metabolism | 21 | 0.33871 | 2 | 0.043495 | 0.33214 |
| Linoleic acid metabolism | 5 | 0.080645 | 1 | 0.078183 | 0.54728 |
| Glyoxylate and dicarboxylate metabolism | 32 | 0.51613 | 2 | 0.092276 | 0.59625 |
| Thiamine metabolism | 7 | 0.1129 | 1 | 0.10778 | 0.6467 |
| Valine, leucine and isoleucine degradation | 40 | 0.64516 | 2 | 0.13431 | 0.71647 |
| Ubiquinone and other terpenoid-quinone biosynthesis | 9 | 0.14516 | 1 | 0.13647 | 0.71647 |
| Primary bile acid biosynthesis | 46 | 0.74194 | 2 | 0.16824 | 0.8122 |
| Fatty acid biosynthesis | 47 | 0.75806 | 2 | 0.17404 | 0.8122 |
| Arginine biosynthesis | 14 | 0.22581 | 1 | 0.20437 | 0.90351 |
| Nicotinate and nicotinamide metabolism | 15 | 0.24194 | 1 | 0.21732 | 0.91272 |
| Histidine metabolism | 16 | 0.25806 | 1 | 0.23006 | 0.92025 |
| Selenocompound metabolism | 20 | 0.32258 | 1 | 0.27909 | 0.94879 |
| Fructose and mannose metabolism | 20 | 0.32258 | 1 | 0.27909 | 0.94879 |
| Citrate cycle (TCA cycle) | 20 | 0.32258 | 1 | 0.27909 | 0.94879 |
| Purine metabolism | 65 | 1.0484 | 2 | 0.28238 | 0.94879 |
| Pentose phosphate pathway | 22 | 0.35484 | 1 | 0.30246 | 0.97718 |
| Galactose metabolism | 27 | 0.43548 | 1 | 0.35776 | 1 |
| Porphyrin and chlorophyll metabolism | 30 | 0.48387 | 1 | 0.38889 | 1 |
| Cysteine and methionine metabolism | 33 | 0.53226 | 1 | 0.41857 | 1 |
| Arachidonic acid metabolism | 36 | 0.58065 | 1 | 0.44686 | 1 |
| Amino sugar and nucleotide sugar metabolism | 37 | 0.59677 | 1 | 0.45599 | 1 |
| Arginine and proline metabolism | 38 | 0.6129 | 1 | 0.46498 | 1 |
| Fatty acid elongation | 39 | 0.62903 | 1 | 0.47383 | 1 |
| Fatty acid degradation | 39 | 0.62903 | 1 | 0.47383 | 1 |
| Pyrimidine metabolism | 39 | 0.62903 | 1 | 0.47383 | 1 |
| Tryptophan metabolism | 41 | 0.66129 | 1 | 0.4911 | 1 |
| Tyrosine metabolism | 42 | 0.67742 | 1 | 0.49953 | 1 |

**Table S14: Enriched Pathway in 4B compared to 4D**

| Pathway name | Total | Expected | Hits | Raw p | FDR |
|---|---|---|---|---|---|
| **Aminoacyl-tRNA biosynthesis** | **48** | **0.77419** | **11** | **2.46E-11** | **2.06E-09** |
| **Valine, leucine and isoleucine biosynthesis** | **8** | **0.12903** | **4** | **3.54E-06** | **0.000149** |
| **Pantothenate and CoA biosynthesis** | **19** | **0.30645** | **5** | **7.16E-06** | **0.000201** |
| **Phenylalanine, tyrosine and tryptophan biosynthesis** | **4** | **0.064516** | **2** | **0.00147** | **0.030867** |
| Taurine and hypotaurine metabolism | 8 | 0.12903 | 2 | 0.006592 | 0.10906 |
| Glutathione metabolism | 28 | 0.45161 | 3 | 0.00931 | 0.10906 |
| Alanine, aspartate and glutamate metabolism | 28 | 0.45161 | 3 | 0.00931 | 0.10906 |
| Phenylalanine metabolism | 10 | 0.16129 | 2 | 0.010387 | 0.10906 |
| Glycine, serine and threonine metabolism | 33 | 0.53226 | 3 | 0.014702 | 0.13721 |
| Biosynthesis of unsaturated fatty acids | 36 | 0.58065 | 3 | 0.018635 | 0.15653 |
| Valine, leucine and isoleucine degradation | 40 | 0.64516 | 3 | 0.024713 | 0.18872 |
| beta-Alanine metabolism | 21 | 0.33871 | 2 | 0.043495 | 0.30446 |
| Glyoxylate and dicarboxylate metabolism | 32 | 0.51613 | 2 | 0.092276 | 0.59625 |
| Thiamine metabolism | 7 | 0.1129 | 1 | 0.10778 | 0.6467 |
| Ubiquinone and other terpenoid-quinone biosynthesi | 9 | 0.14516 | 1 | 0.13647 | 0.76423 |
| Primary bile acid biosynthesis | 46 | 0.74194 | 2 | 0.16824 | 0.85997 |
| Fatty acid biosynthesis | 47 | 0.75806 | 2 | 0.17404 | 0.85997 |
| Arginine biosynthesis | 14 | 0.22581 | 1 | 0.20437 | 0.95371 |
| Nicotinate and nicotinamide metabolism | 15 | 0.24194 | 1 | 0.21732 | 0.96076 |
| Histidine metabolism | 16 | 0.25806 | 1 | 0.23006 | 0.96626 |
| Selenocompound metabolism | 20 | 0.32258 | 1 | 0.27909 | 0.98833 |
| Fructose and mannose metabolism | 20 | 0.32258 | 1 | 0.27909 | 0.98833 |
| Citrate cycle (TCA cycle) | 20 | 0.32258 | 1 | 0.27909 | 0.98833 |
| Purine metabolism | 65 | 1.0484 | 2 | 0.28238 | 0.98833 |
| Pentose phosphate pathway | 22 | 0.35484 | 1 | 0.30246 | 1 |
| Galactose metabolism | 27 | 0.43548 | 1 | 0.35776 | 1 |
| Porphyrin and chlorophyll metabolism | 30 | 0.48387 | 1 | 0.38889 | 1 |
| Cysteine and methionine metabolism | 33 | 0.53226 | 1 | 0.41857 | 1 |
| Arachidonic acid metabolism | 36 | 0.58065 | 1 | 0.44686 | 1 |
| Amino sugar and nucleotide sugar metabolism | 37 | 0.59677 | 1 | 0.45599 | 1 |
| Arginine and proline metabolism | 38 | 0.6129 | 1 | 0.46498 | 1 |
| Fatty acid elongation | 39 | 0.62903 | 1 | 0.47383 | 1 |
| Fatty acid degradation | 39 | 0.62903 | 1 | 0.47383 | 1 |
| Pyrimidine metabolism | 39 | 0.62903 | 1 | 0.47383 | 1 |
| Tryptophan metabolism | 41 | 0.66129 | 1 | 0.4911 | 1 |
| Tyrosine metabolism | 42 | 0.67742 | 1 | 0.49953 | 1 |

**Table S15: Enriched pathway in 4B compared to 4C**

| Name | Total | Expected | Hits | Raw p | FDR |
|---|---|---|---|---|---|
| **Aminoacyl-tRNA biosynthesis** | **48** | **0.71226** | **9** | **7.13E-09** | **5.99E-07** |
| **Pantothenate and CoA biosynthesis** | **19** | **0.28194** | **5** | **4.61E-06** | **0.000193** |
| **Phenylalanine, tyrosine and tryptophan biosynthesis** | **4** | **0.059355** | **2** | **0.001242** | **0.034769** |
| Valine, leucine and isoleucine biosynthesis | 8 | 0.11871 | 2 | 0.005588 | 0.088136 |
| Taurine and hypotaurine metabolism | 8 | 0.11871 | 2 | 0.005588 | 0.088136 |
| Glutathione metabolism | 28 | 0.41548 | 3 | 0.007345 | 0.088136 |
| Alanine, aspartate and glutamate metabolism | 28 | 0.41548 | 3 | 0.007345 | 0.088136 |
| Phenylalanine metabolism | 10 | 0.14839 | 2 | 0.00882 | 0.092614 |
| Glycine, serine and threonine metabolism | 33 | 0.48968 | 3 | 0.011653 | 0.10876 |
| beta-Alanine metabolism | 21 | 0.31161 | 2 | 0.037281 | 0.31316 |
| Glyoxylate and dicarboxylate metabolism | 32 | 0.47484 | 2 | 0.079815 | 0.6095 |
| Biosynthesis of unsaturated fatty acids | 36 | 0.53419 | 2 | 0.097841 | 0.64321 |
| Thiamine metabolism | 7 | 0.10387 | 1 | 0.099544 | 0.64321 |
| Ubiquinone and other terpenoid-quinone biosynthesis | 9 | 0.13355 | 1 | 0.1262 | 0.75718 |
| Primary bile acid biosynthesis | 46 | 0.68258 | 2 | 0.14716 | 0.79988 |
| Fatty acid biosynthesis | 47 | 0.69742 | 2 | 0.15236 | 0.79988 |
| Arginine biosynthesis | 14 | 0.20774 | 1 | 0.18957 | 0.93669 |
| Nicotinate and nicotinamide metabolism | 15 | 0.22258 | 1 | 0.2017 | 0.94128 |
| Histidine metabolism | 16 | 0.23742 | 1 | 0.21367 | 0.94462 |
| Purine metabolism | 65 | 0.96452 | 2 | 0.25062 | 0.94723 |
| Selenocompound metabolism | 20 | 0.29677 | 1 | 0.25982 | 0.94723 |
| Fructose and mannose metabolism | 20 | 0.29677 | 1 | 0.25982 | 0.94723 |
| Citrate cycle (TCA cycle) | 20 | 0.29677 | 1 | 0.25982 | 0.94723 |
| Pyruvate metabolism | 22 | 0.32645 | 1 | 0.28191 | 0.94723 |
| Pentose phosphate pathway | 22 | 0.32645 | 1 | 0.28191 | 0.94723 |
| Glycolysis / Gluconeogenesis | 26 | 0.38581 | 1 | 0.32422 | 1 |
| Galactose metabolism | 27 | 0.40065 | 1 | 0.33442 | 1 |
| Porphyrin and chlorophyll metabolism | 30 | 0.44516 | 1 | 0.36414 | 1 |
| Cysteine and methionine metabolism | 33 | 0.48968 | 1 | 0.39259 | 1 |
| Amino sugar and nucleotide sugar metabolism | 37 | 0.54903 | 1 | 0.42863 | 1 |
| Arginine and proline metabolism | 38 | 0.56387 | 1 | 0.43732 | 1 |
| Fatty acid elongation | 39 | 0.57871 | 1 | 0.44588 | 1 |
| Fatty acid degradation | 39 | 0.57871 | 1 | 0.44588 | 1 |
| Pyrimidine metabolism | 39 | 0.57871 | 1 | 0.44588 | 1 |
| Valine, leucine and isoleucine degradation | 40 | 0.59355 | 1 | 0.45431 | 1 |
| Tryptophan metabolism | 41 | 0.60839 | 1 | 0.46262 | 1 |
| Tyrosine metabolism | 42 | 0.62323 | 1 | 0.47081 | 1 |
